# A competitive disinhibitory network for robust optic flow processing in *Drosophila*

**DOI:** 10.1101/2023.08.06.552150

**Authors:** Mert Erginkaya, Tomás Cruz, Margarida Brotas, Kathrin Steck, Aljoscha Nern, Filipa Torrão, Nélia Varela, Davi Bock, Michael Reiser, M Eugenia Chiappe

## Abstract

Many animals rely on optic flow for navigation, using differences in eye image velocity to detect deviations from their intended direction of travel. However, asymmetries in image velocity between the eyes are often overshadowed by strong, symmetric translational optic flow during navigation. Yet, the brain efficiently extracts these asymmetries for course control. While optic flow sensitive-neurons have been found in many animal species, far less is known about the postsynaptic circuits that support such robust optic flow processing. In the fly *Drosophila melanogaster*, a group of neurons called the horizontal system (HS) are involved in course control during high-speed translation. To understand how HS cells facilitate robust optic flow processing, we identified central networks that connect to HS cells using full brain electron microscopy datasets. These networks comprise three layers: convergent inputs from different, optic flow-sensitive cells, a middle layer with reciprocal, and lateral inhibitory interactions among different interneuron classes, and divergent output projecting to both the ventral nerve cord (equivalent to the vertebrate spinal cord), and to deeper regions of the fly brain. By combining two-photon optical imaging to monitor free calcium dynamics, manipulating GABA receptors and modeling, we found that lateral disinhibition between brain hemispheres enhance the selectivity to rotational visual flow at the output layer of the network. Moreover, asymmetric manipulations of interneurons and their descending outputs induce drifts during high-speed walking, confirming their contribution to steering control. Together, these findings highlight the importance of competitive disinhibition as a critical circuit mechanism for robust processing of optic flow, which likely influences course control and heading perception, both critical functions supporting navigation.

## Introduction

As we navigate through space, our senses are continuously bombarded with a variety of stimuli from the environment and our own movements. Even for seemingly straightforward binary information, such as determining if we are veering to the left or right, our brains must extract and process the relevant signals from all incoming inputs to reorient according to our goals. Sensorimotor circuits that combine feedforward excitation with lateral inhibition have frequently been associated with this type of binary categorization^1,2^ and behavioral choice^3–5^. However, how inhibitory interactions transform sensory information into categorical signals for continuous movement control is poorly understood.

Many animals rely on coherent patterns of image velocity across the retina to navigate according to goals. These patterns result from the relative motion between the moving observer and the surrounding scene, commonly known as optic flow^6,7^. The specific optic flow experienced by the observer depends on the direction and speed of the eyes, head, and body. Therefore, optic flow provides valuable information about gaze, heading and traveling directions, allowing for a comparison with the intended trajectories for course control. Indeed, optic flow-sensitive neurons, which poses large monocular receptive fields sensitive to different axes of movement^8–11^ are found across the animal kingdom^12–17^ and have been implicated in heading perception, and gaze and course control^18–23^. Additionally, optic-flow sensitive neurons integrate multimodal information to distinguish between self-generated and externally generated information^13,24–27^. However, for accurate extraction of self-motion information, binocular integration between the eyes is also necessary. While some lateral interactions within optic flow-sensitive neurons have been identified^17,28–31^, the precise mechanisms through which postsynaptic circuits integrate bilateral optic flow information to disambiguate self-motion and contribute to goal-directed navigation remain largely unknown. The fly *Drosophila melanogaster* offers a valuable opportunity to investigate the circuit mechanisms involved in processing optic flow robustly. With its impressive visually guided behaviors, versatile genetic toolkit, comprehensive brain electron microscopy datasets, and well-characterized populations of optic flow-sensitive neurons, *Drosophila* allows for a systematic identification, and precise dissection of these mechanisms. In *Drosophila*, optic flow-sensitive neurons called lobula plate tangential cells (LPTCs) are found in deep layers of the fly’s optic lobe and project to premotor regions of the brain. Certain LPTCs, like the horizontal system (HS) cells and H2 cells are sensitive to horizontal optic flow^32–34^. They receive non-visual signals related to angular velocity, which enhance their selectivity to the fly’s rotations during spontaneous walking^27,33^. HS and H2 cells can also drive rotations of the fly^27,33,35^, implicating them in course control. Recent work has shown that HS cells play a significant role in steering an exploratory fly, particularly during high-speed walking^35,36^, when optic flow is dominated by symmetric translational components, which may overshadow the asymmetric ones caused by the fly’s unexpected rotations. Thus, investigating HS cells and their postsynaptic pathways provides an opportunity to understand how the network process optic flow robustly, even in situations with a low signal-to-noise ratio.

In this study, we conducted a systematic analysis of the structure and function of synaptic partners of HS and H2 at their central axon terminals. Our findings revealed that HS and H2 cells form a subnetwork comprising three layers, including an input layer of converging LPTCs, a middle layer of several classes of GABAergic interneurons, and divergent outputs projecting back to the lobula plate (feedback), to the ventral nerve cord (VNC, the insect analogous of the vertebrate spinal cord), or to deeper premotor regions of the fly brain. The inhibitory layer forms recurrent connections with LPTCs and provides lateral inhibition across the two brain hemispheres. Using two-photon calcium imaging of the newly identified interneurons along with manipulating GABA receptors and modeling, we found that a lateral disinhibition motif enhances rotational sensitivity at the network’s output layer, supporting steering control in freely exploring flies. Overall, our study unveils the layered anatomical structure of optic flow networks and their strong modulation by inhibitory feedback and elucidates how feedforward excitation combined with lateral disinhibition enables robust extraction of asymmetries in optic flow signals between the eyes to guide reorientation of a roaming fly.

## Results

### A class of optic flow-sensitive descending neuron contributes to steering during high-speed walking

Course deviations can be determined by analyzing the differences in the pattern of optic flow between the fly’s eyes using binocular integration. However, these differences vary with the fly’s walking speed. Even when the deviation from a straight course is the same, flies moving at low versus high speed induce distinct optic flow patterns (**Fig. 1a**, colored rectangles). At low walking speeds, asymmetries in the optic flow can be easily detected, but at high speed, these asymmetries are largely reduced due to the prominent translational components in the optic flow (**Fig. 1a**). We currently lack a complete understanding of how the networks responsible for processing rotational optic flow estimate asymmetries that remain consistent regardless of the fly speed, enabling reliable walking control.

**Figure 1:**
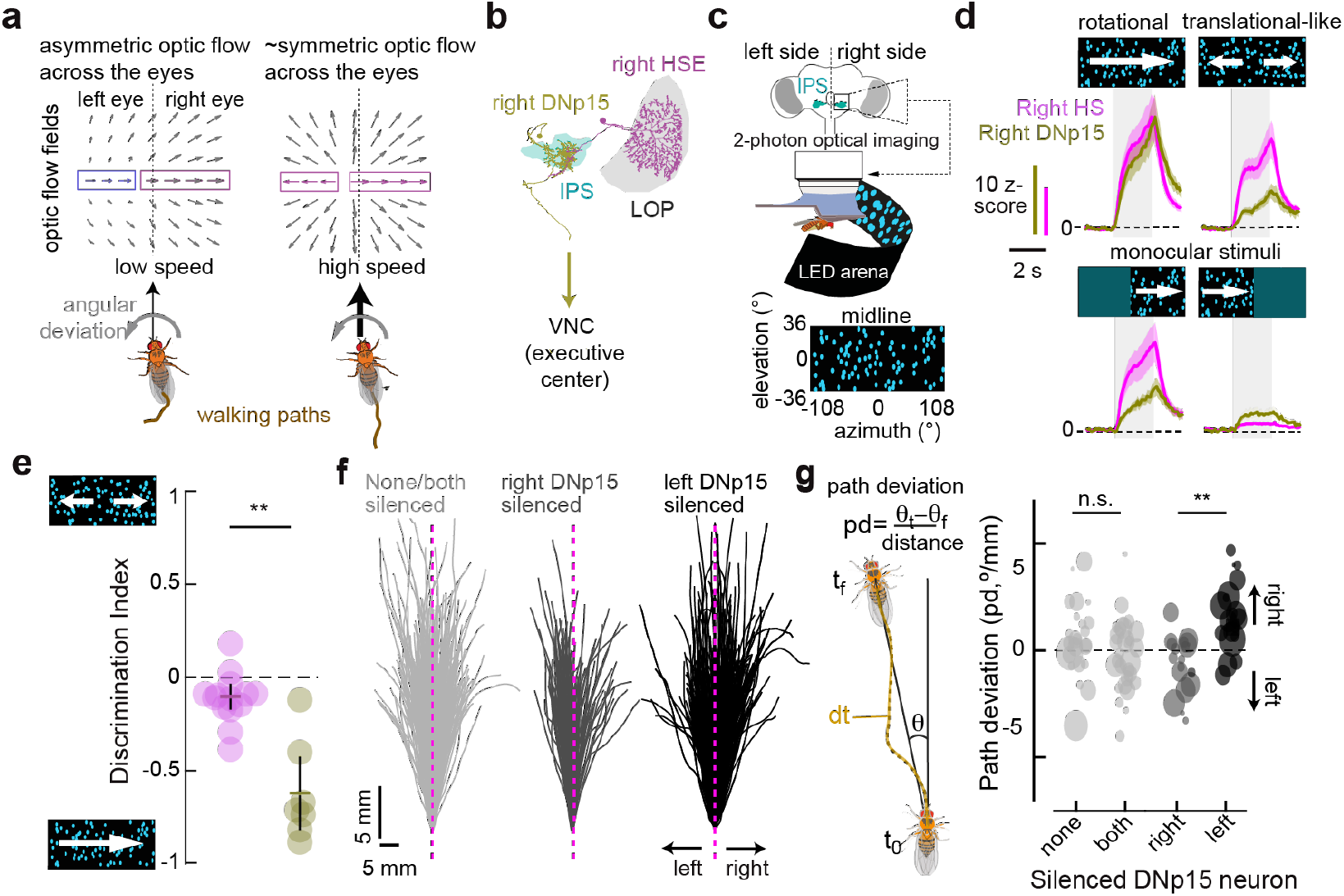
DNp15 exhibit enhanced sensitivity to rotational optic flow compared to HS and plays a role in steering during high-speed translation. (a) Simulated binocular optic flow of a fly walking at low (left) versus high (right) forward speed and rotating with the same angular velocity. Rectangles indicate the presence of regressive (blue) or progressive (magenta) motion detected by a horizontal motion-sensitive system. (b) EM-reconstructed right HS (magenta) and postsynaptic right DNp15 (olive) cells. The chemical and electrical synapses between the two cells allocated within the inferior posterior slope (IPS, cyan), a premotor region. The dendrites of HS cells are located within the lobula plate (LOP). (c) Top, schematic of the fly brain and the IPS, where optical imaging was performed. Bottom, 2-photon calcium imaging setup and the visual LED display. (d) HS (magenta) and DNp15 (olive) calcium responses to horizontal binocular rotational or translational-like optic flow (top), and to monocular stimuli (bottom). (e) Discrimination index (see Methods) between translational-like and rotational stimuli in HS (magenta) and DNp15 (olive). Each circle represents an individual ROI. Colored horizontal line and black vertical line represent the mean and 95% CI. (HS: N=12 flies, n=16 ROIs, DNp15: N=6 flies n=7 ROIs, *P < 0.05; **P < 0.01 ***P < 0.001, Wilcoxon’s rank-sum test). (f) The walking paths of forward runs of length ≥ 4 body lengths, aligned at their origin, in flies with either both or none DNP15 silenced (left, light gray traces), right DNp15 silenced (middle, gray traces), and left DNp15 silenced (right, black traces). (g) Left, definition of path deviation, the difference in body orientation between the end and the beginning of the walking path over traveled distance. Right, mean path deviation from per fly in flies with none, single or both DNp15 silenced. The size of the circle indicates the number of walking bouts per fly, with larger sizes representing larger number of walking bouts. Number of flies per condition, none n=25; both n=31; left n=22; right n=22; *p < 0.05 **p < 0.01 ***p < 0.001, Wilcoxon’s rank-sum test with Bonferroni correction for multiple comparisons.

The horizontal system cells (HS cells), whose receptive fields are predominately monocular^32^, play a role in steering, especially during high-speed walking^35,36^. Recent work has shown that these cells interact laterally (across hemispheres) via gap junction coupling through an unidentified interneuron^30^, potentially improving the detection of asymmetric optic flow. If these lateral interactions enhanced the detection of asymmetric optic flow, output neurons from the HS network should display increased sensitivity to rotational optic flow and binocular activity compared to HS cells.

To explore this idea, we compared the response properties of the only known descending neuron postsynaptic to HS cells, DNp15^30,37^ to those of HS cells concerning optic flow (**Fig. 1b,c**). We conducted *in vivo*, 2-photon calcium imaging in immobilized flies within the inferior posterior slope (IPS) region of the fly brain (**Fig. 1c**, see **Methods**) while presenting binocular asymmetric horizontal motion (i.e., stimuli representing rotations along the yaw axis) or binocular symmetric horizontal motion (i.e., stimuli representing translational-like movement along the forward direction (**Supplementary Video 1**, “Front-to-back/Back-to-front” and “Yaw left/right” stimuli, respectively). In addition, to identify binocular interactions, we compare the binocular responses of HS and DNp15 cells to those induce by monocular stimuli.

We found that DNp15 exhibited response profiles and velocity tuning to binocular asymmetric optic flow like HS cells, likely due to their electrical coupling^37^ (**Fig. 1-S1a-c, Supplementary Video 2**). However, DNp15 activity was more influenced by binocular interactions compared to HS cells and showed a small but detectable response to contralateral back-to-front visual motion (**Fig. 1d, 1-S1c**), suggesting an increased selectivity to rotational optic flow. To describe the relative selectivity of HS and DNp15 to rotational vs. translational stimuli, we defined a qualitative parameter called the discrimination index, which compares the magnitude of the responses of a cell to rotational vs translational stimuli (see **Methods**, **Fig. 1-S1e**). Overall, DNp15 demonstrated enhanced selectivity to rotational stimuli than translational-like stimuli, unlike HS cells (**Fig. 1d,e, 1-S1d,e**). Thus, the network associated with HS cells seems well suited to robustly detect asymmetries in binocular optic flow for course control.

To test this idea, we investigated the role of DNp15 in walking control. DNp15 is thought to contribute to gaze stabilization^37^ and directly projects to dorsal neuropiles within the tectulum of the ventral nerve cord^38^ (VNC, the insect analogous of the spinal cord). Gaze control depends on head-body coordination and course control^39^. Thus, we directly tested the impact of DNp15 silencing on steering (course control) during highspeed walking by expressing the inward rectifying potassium channel Kir2.1 in different configurations and observing the walking paths of exploratory flies^39^. We created flies with unperturbed activity of DNp15 (controls, i.e., no expression of Kir2.1), with balanced silenced DNp15 (bilateral expression of Kir2.1), and with asymmetry in the activity of DNp15 (unilateral expression of Kir2.1) (**Fig. 1-S1f**, see **Methods**). Flies typically exhibit runs of high forward velocity or turns of low forward velocity (body saccades) under these experimental conditions (**Fig. 1f**, see also **Fig. 6a**). We evaluated course control by calculating path deviation per run bout, or by examining any biases during body saccades (**Fig. 1f,g**, **Fig. 1-S1g**, see **Methods**). Flies with both DNp15 silenced or with no silencing did not show any bias during body saccades or runs (**Fig. 1f,g** and **Fig. 1-S1g**, none or both silenced DNp15). However, unilateral silencing of DNp15 resulted in contralateral path deviation during runs (**Fig. 1f,g**), while no biases were observed during body saccades (**Fig. 1-S1g**, left or right silenced DNp15). These findings suggest that asymmetry in activity of left vs. right DNp15 leads to steering biases during forward walking. Furthermore, the enhanced selectivity of DNp15 to rotational vs. translational optic flow implies that these neurons could detect minor differences in optic flow across the eyes, thereby contributing to steering at high forward speed. However, the specific circuits and mechanisms responsible for conveying such selectivity supporting robust optic flow processing in DNp15 remain unknown.

### HS, H2 and VS cells connect distinctively to two parallel central networks

To uncover the circuit and mechanisms responsible for robust rotational optic flow processing, we investigated the downstream central synaptic partners of well-characterized rotational optic flow sensitive cells, such as HS, H2, and VS cells. We used a complete female adult fly brain (FAFB) electron microscopy (EM) dataset^40^ (**Fig. 1-S2a-b**, see **Methods**) to map the location of chemical synapses at the axon terminals of these LPTCs. By manually tracing and identifying pre and postsynaptic partners (**Fig. 1-S2c**, see **Methods** for tracing strategy), we found that HS, H2 and VS cells establish both input and output synaptic sites, revealing central modulation of axonal activity in HS, H2 and VS cells (**Fig. 1-S2c-d**). We also observed that these cells form few strong connections and numerous weak connections with their synaptic inputs and outputs (**Fig. 1-S3a,b**). These results were confirmed using two additional EM datasets with semi-automatically reconstructed neurons, FlyWire^41^ and Hemibrain^42^ (**Fig. 1-S2a,c,d, S3**, **Supplementary Table 1**). Hence, our manual tracing approach effectively identified important synaptic partners of HS, VS, and H2 cells. We defined neurons providing more than 10 synaptic inputs to or receiving more than 35 synaptic outputs from LPTCs as strong synaptic partners (**Fig. 1-S2d**, black lines).

To analyze the central networks involving these LPTCs, we computed a connectivity matrix among all identified strong partners in the FAFB dataset (**Fig. 2-S1**, see **Methods**). While the matrix displayed rather sparse connections with weak or no connections between most neuron pairs, certain pairs exhibited strong unidirectional connections with hundreds of synapses (**Fig. 2-S1**). We performed a hierarchical cluster analysis on the pairwise number of synapses (**Fig. 2-S1**, see **Methods**), and identified three primary clusters: HS and VS cells with the interneuron lVLPT8, and two clusters forming separate networks with HS and VS cells (**Fig. 2-S1** and **2a**). Importantly, this network structure remains mostly unchanged by thresholding (**Fig. 2-S2**).

The VS network demonstrated strong connections between VS and central interneurons (**Fig. 2a**), with output pathways diverging into descending pathways, including neck motor neurons, and descending neurons (DNs), and feedback pathways to the ocelli, the secondary optical system of flies^43,44^. On the other hand, the H2-HS network exhibited strong connections among central interneurons, H2 and HS cells, and the GABAergic feedback CH cells^29,45,46^ (**Fig. 2a**). Like the VS network, the outputs of the HS-H2 network diverged, with a descending pathway defined by several DNs projecting to leg, neck, and haltere neuropiles in the VNC^38,47^, and an ascending pathway of interneurons projecting to the lateral accessory lobes (LAL).

**Figure 2.**
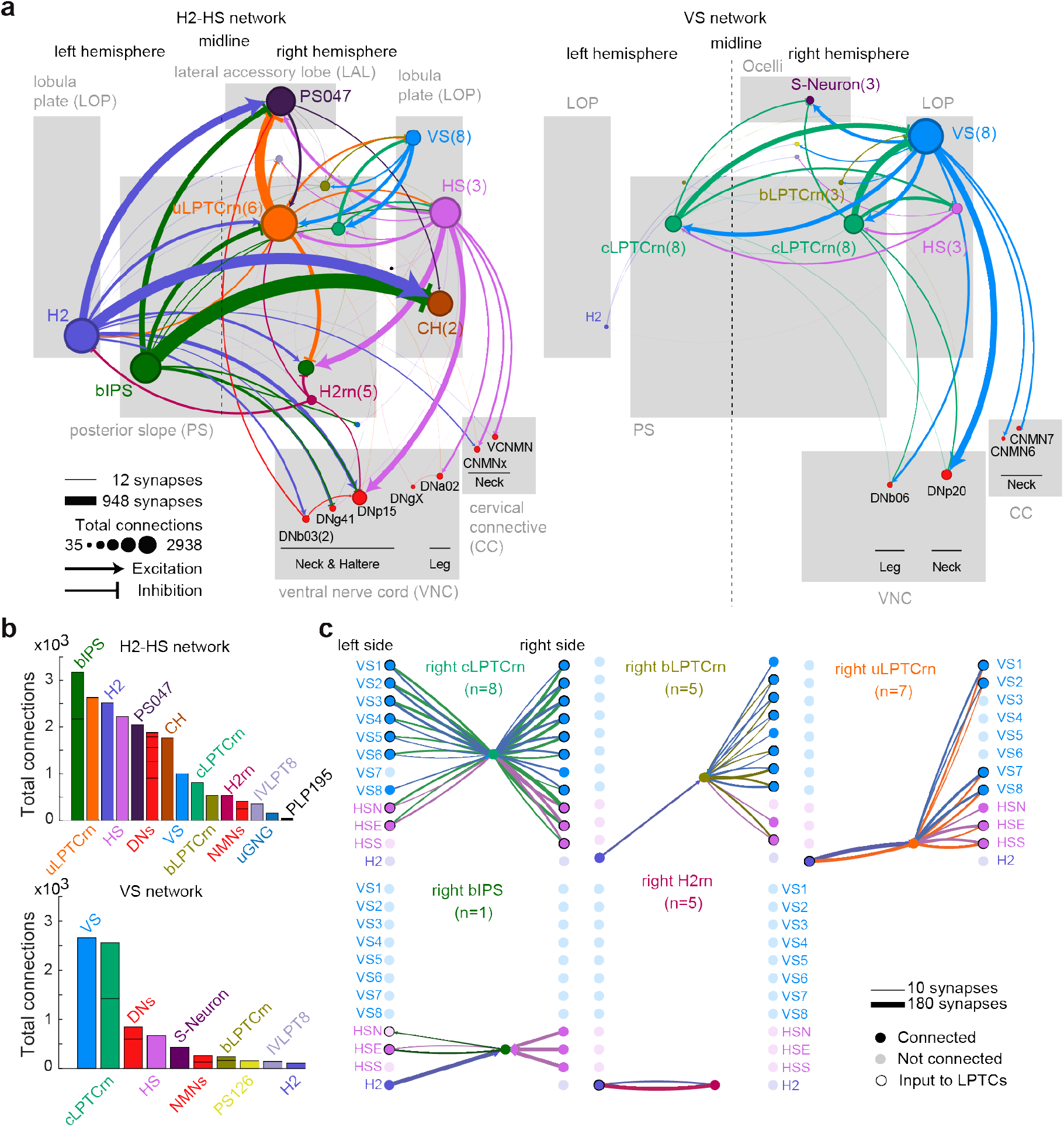
H2, HS and VS cells form two distinctive networks in the central brain. (a) Connectivity graph of the H2-HS (left) and VS (right) networks. Neurons are represented as nodes (circles), with their synaptic connections indicated by arrows (predicted to be excitatory) or tee-shaped edges (predicted to be inhibitory). Thickness of the arrows/tee-shaped edges represents number of synapses, and color indicates the presynaptic neuron. For clarity, connections with fewer than 10 synapses are not shown; connections with fewer than 50 synapses are shaded. Node diameters represent the total number of connections that each node has within the network (including both inputs and outputs). Except for LPTCs, nodes are placed within the brain regions (grey rectangles) where their axons reside. (b) The total number of synaptic connections for each neuron within H2-HS network (Top), and VS network (bottom). See Supplementary Table 2 for the complete list of neurons and nomenclature. (c) Connectivity graph between each LPTC input cell type and individual HS, H2 and VS cells. Edges indicate synaptic connection between cells, with thickness indicating number of synapses and color indicating the presynaptic cell. LPTCs without input are transparent for clarity.

The VS connections represented only a small proportion of total connections within the H2-HS network and vice versa (**Fig. 2b**). This suggests that both networks largely operate independently from each other. Notably, the DNs of the HS-H2 network are different from those of the VS network, pointing towards parallel operation. Two DN outputs of the HS-H2 network, DNa02 and DNp15 have been previously characterized (**Fig. 1**)^37,47^. These neurons are sensitive to fly rotations (or their visual consequence) and contribute to steering. The LAL, the other output region of the HS-H2 network, is a higher-order premotor area of the insect brain, thought to contribute to steering^48,49^. Together, these findings strongly support the involvement the HS-H2 network in steering control.

### Recurrent connections define the intermediate layer structure of both networks

The two networks, though largely non-overlapping, converge at the level of LPTC cells, connecting to a small set of interneurons within the intermediate layers of the networks (i.e., between LPTCs and their outputs). We identified five distinct classes of LPTC input cells within these intermediate layers that provide specific, recurrent input to HS, H2 and VS cells, thereby defining the structure of the two networks (**Fig. 2-S3a-c**). Three of the five classes are collectively defined as lobula plate tangential cell recurrent neurons (LPTCrns), due to their shared soma location and tract, indicative of a common developmental origin. We differentiate them based on anatomical features (**Fig. 2-S3b-d**).

Crab LPTCrn cells (**cLPTCrn**), central components of the VS network (**Fig. 2b**), extend prominent projections to both hemispheres of the PS and connect reciprocally with most VS and HS cells (**Fig. 2c**). Bilateral LPTCrn cells (bLPTCrn) also project to both hemispheres of the PS, but their contralateral projections are limited to the dorsal area of the PS (**Fig. 2-S3b**). Unilateral LPTCrn cells (uLPTCrn) are prominent components of the H2-HS network (**Fig. 2b**). They project to the ipsilateral side and exhibit strong reciprocal connections with HS, H2, and VS1-2, and VS7,8 cells (**Fig. 2c**).

The remaining two LPTC input classes specifically connect to HS and H2 cells. The first class involves bilateral inferior posterior slope neurons (bIPS, **Fig 2-S3b**), which receive strong convergent projections from ipsilateral HS and contralateral H2 cells and send projections to the contralateral HSN and HSE cells (**Fig. 2c**). Notably, the two contralateral CH cells are the strongest downstream partners of bIPS, receiving over 450 synapses from bIPS as their strongest input (**Fig. 2-S1**). This provides strong anatomical evidence for a central feedback pathway within this network. The second class of interneurons selectively connects to the contralateral H2 cells at their axon terminals (**Fig. 2c**), referred as H2 recurrent neurons (H2rn) as they receive input back from contralateral H2 cells.

The primary brain regions that send information to the intermediate layers of the VS and HS-H2 are the PS, gnathal ganglion (GNG), and optic lobes. Additionally, all five cell types receive direct ascending input from the VNC (**Fig. 2-S3e, Fig. 3b**). Thus, the intermediate layers may operate as both integrators and broadcasters: they likely integrate multimodal signals to modulate the optic flow sensitive networks, and broadcast optic flow signals to numerous regions of the fly’s CNS. The recurrent connections between LPTCs and different classes of LPTC input neurons in the central brain suggest that these interneurons possess distinct optic flow sensitivities based on their LPTC inputs, integrate extraretinal signals in addition to optic flow, broadcast modulatory signals from premotor centers, and shape the optic flow sensitivity of output neurons. Different sets of central interneurons within the intermediate layers of the two networks may contribute in different visual-motor contexts. In the following sessions, we focus on the functional properties of neurons within the different layers of the HS-H2 network, and how optic flow signals are transformed to enhance DNp15’s selectivity to horizontal optic flow compared to HS cells.

**Figure 3:**
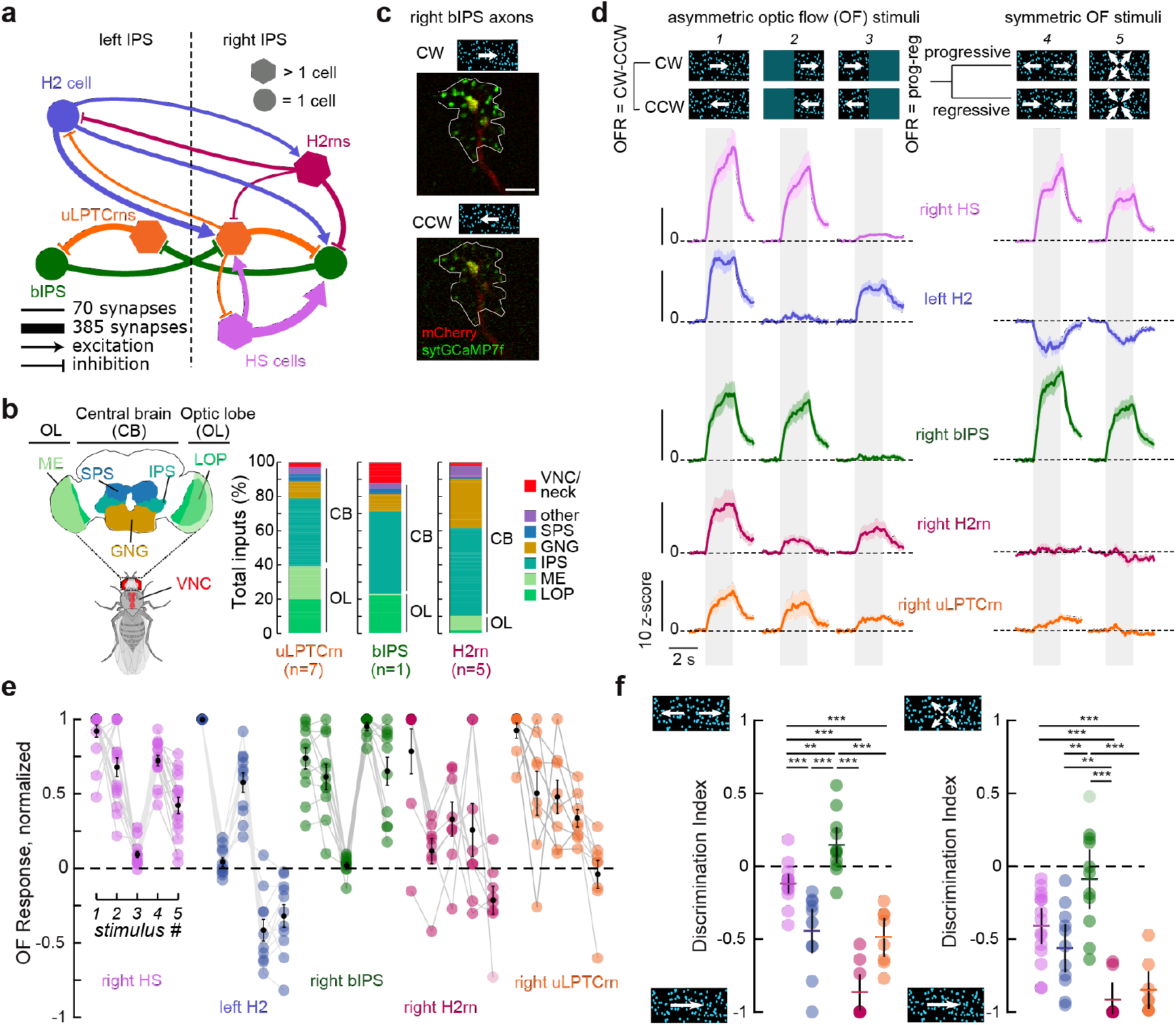
Binocular interactions define optic flow sensitivity within the H2-HS network. (a) Local connectivity between the input and middle layers of the H2-HS network. (b) Left, schematic of the central nervous system (CNS) regions from which uLPTCrns, bIPS and H2rns receive inputs. LOP, lobula plate; ME, medulla; GNG, gnathal ganglia; IPS, inferior posterior slope; SPS, superior posterior slope. Right, proportion of inputs per CNS region (color code) for each interneuron. (c) Single plane fluorescence image of an example recording from bIPS axons during binocular CW (top) and CCW (bottom) horizontal motion. We defined a single region of interest (ROI), which covers the entire axon is outlined in white. The red signal corresponds to mCherry fluorescence, whereas the green signal corresponds to sytGCaMP7f fluorescence. Scale bar, 10 μm. (d) Top: visual stimuli presented to the fly, with direction of motion as indicated by arrows. Bottom: optic flow calcium responses (OFR) of each neuron to rotational (stimuli 1, 2, 3) or translational (stimuli 4, 5) optic flow. OFR for rotational stimuli is defined as CW (rightwards) minus CCW (leftwards) direction of visual motion. For the translational stimuli, OFR is defined as progressive minus regressive direction of visual motion. Each trace corresponds to the grand mean of the z-scored calcium response across flies ± SEM (shade). Per fly, we obtained mean responses to 5-10 repetitions per stimulus. Light-gray rectangles indicate the stimulation window. For this and the remaining panels: HS: N=12 flies n=16 ROIs, H2: N =8 flies n=11 ROIs, bIPS: N=10 flies n=11 ROIs, H2rn: N=5 flies n=8 ROIs, uLPTCrn: N=6 flies n=8 ROIs. (e) Normalized OFRs (relative to the preferred stimulus per neuron). Error bars represent SEM of the grand mean across flies. (f) Left, discrimination index between translation-like vs. rotational stimuli (see Methods). Each circle represents an individual ROI. The horizontal line and black vertical line represent mean and 95% CI. Right, discrimination index between translation vs. rotational stimuli. *: p < 0.05; **p < 0.01; ***p < 0.001, Wilcoxon’s rank-sum test with Bonferroni correction for multiple comparisons.

### The H2-HS network exhibit responses to optic flow that are not entirely predicted by their LPTC inputs

To understand how the enhanced selectivity to rotational optic flow in DNp15 emerges from the structure of the HS-H2 network, we characterized the optic flow responses of the intermediate layer of the H2-HS network, which connect to DNp15 (**Fig. 2-S3f, Fig. 3a**). To achieve cell-type-specific genetic access, we used traced EM skeletons to identify neurons in transgenic line libraries (see **Methods, Fig. 3-S1a**). In vivo, neural recordings were conducted under non-behaving conditions, focusing specifically on optic flow processing while minimizing the potential impact of non-visual signals present in both LPTCs^25,27,33,50,51^ and the network’s interneurons (**Fig. 3b**). We characterized neural responses to various patterns of optic flow (**Supplementary Video 1**) by measuring calcium dynamics at axon terminals (e.g., **Fig. 3c**, see **Methods** for choice of calcium indicators).

bIPS, uLPTCrn, and H2rn cells displayed responses to horizontal optic flow and velocity tuning that showed similarities and differences compared to their inputs HS and H2 cells, (**Fig. 3d, Fig. 3-S2, Supplementary Video 2)**. First, uLPTCrn cells, which receive direct input from VS (**Fig. 2c**), exhibited minimal or negative calcium signals to pitch and roll optic flow, which would be expected to excite their presynaptic inputs from VS cells (**Fig. 3-S2a**). Second, each class of interneurons displayed characteristic velocity tuning. While uLPTCrns are more sensitive to slower than faster speeds, bIPS exhibit larger responses to intermediate speeds, and H2rns are more sensitive to faster than slower speeds, in contrast to their presynaptic H2 cells. These results suggest that cell autonomous properties or additional inputs to these interneurons modulate visual responses of uLPTCrns and H2rns, either through direct connections from other optic lobe neurons, or indirectly through inputs from the central brain (**Fig. 3b**).

Further evidence for network-based modulation of visual inputs came from the analysis of the responses of interneurons to monocular stimuli. bIPS and uLPTCrn cells receive converging inputs from ipsilateral HS and contralateral H2 neurons (**Fig. 3a**), which respond to front-to-back (FTB) and back-to-front (BTF) monocular motion, respectively (**Fig. 3-S2c**). Accordingly, one would expect both cell types to respond to ipsilateral progressive and contralateral regressive stimuli. However, while both cell types responded to progressive stimuli, they showed minimal responses to contralateral regressive motion (**Fig. 3-S2c**), unlike their H2 inputs. Conversely, H2rn cells responded to contralateral regressive motion, as excepted from their inputs from H2 cells, but they also responded to progressive motion (**Fig. 3-S2c**) despite not receiving direct input from HS cells. Thus, additional inputs from the central brain likely convey visual information indirectly to H2rn cells. Nevertheless, responses to binocular asymmetric stimuli were larger than monocular stimuli in all cells (**Fig. 3d-e**), suggesting binocular interactions at the intermediate layer. Together, these findings demonstrate that the sensitivity to optic flow in different classes of interneurons within the H2-HS network are modulated by network activity and their central inputs, with strong binocular interactions. Specifically, both uLPTCrns and bIPS display binocular optic flow selectivity that cannot be entirely explained by their direct H2 inputs.

### Neurons within the H2-HS network show distinctive optic flow selectivity

The lack of responses to contralateral regressive motion in bIPS and uLPTCrn suggests that these interneurons might be equally sensitive to translational and rotational optic flow. To test this idea, we examined their responses to binocular symmetric and asymmetric stimuli (**Fig. 3-S2d**). We presented binocular translational motion or binocular symmetrical horizontal motion (translation-like stimulus). The former stimuli contain progressive or regressive horizontal components combined with vertical downward or upward components, respectively, to mimic the visual consequence of the body’s forward or backward translation. The latter stimuli contain only horizontal progressive or regressive components, representing what a horizontal-selective system may respond to in the context of translation (**Supplementary Video 1,** “Progressive/regressive” and “Front-to-back/Back-to-front”, respectively). The three interneurons exhibited characteristic response profiles to these stimuli (**Fig. 3d,e, Fig. 3-S2d**). bIPS was most active with binocular symmetric horizontal stimuli compared to translational progressive stimuli (**Fig. 3d,e**). uLPTCrn cells showed minimal or negative responses to translational patterns, while H2rns showed positive response to both regardless of the direction of motion, albeit weakly (**Fig. 3d,e, 3-S2d**).

To describe the relative selectivity of HS, H2, bIPS, uLPTCrns, and H2rns to rotational vs. translational stimuli by calculating the discrimination index (see **Methods**, **Fig. 1-S1e**, **3f**). HS and H2 responded to both rotational and translational optic flow. However, they were relatively, albeit weakly, more selective to rotational optic flow, resulting in a slightly negative discrimination index closer to zero compared to uLPTCrn and H2rn cells, which showed enhanced selectivity for rotational optic flow, particularly when compared to the more naturalistic translational optic flow. bIPS also responded to both rotational and translational patterns in a selective manner; however, bIPS exhibited a relatively stronger sensitivity to binocular horizontal symmetric stimuli and thus have a discrimination index skewed more towards translations compared to rest of the H2-HS network cells (**Fig. 3f**).

We expanded our translational optic flow characterization (**Fig. 3-S3a**, **Supplementary Video 1**) and observed that bIPS, like HS and H2 cells, exhibited its strongest translational sensitivity to compound optic flow-fields, with sideslip and forward components (**Fig. 3-S3b-d**). However, translational patterns with backwards motion which elicited activity in H2 cells (**Fig. 3-S3c**) have minimal impact on bIPS. H2rns and uLPTCrns on the other hand, minimally increased or decreased their activity during forward and backward translations (**Fig. 3-S3e-f**), becoming mostly sensitive to changes in sideslip translations (**Fig. 3-S3g**). Taken together, these results demonstrate that neurons downstream of HS and H2 cells have increased selectivities to characteristic patterns of optic flow. The question remains as to how a neuron like bIPS, which is embedded in a rotational-sensitive network, enhances its selectivity for translational or translational-like optic flow patterns, and what function this response property provides to the network.

### Inhibitory interactions fine tune optic flow sensitivity in bIPS

We hypothesized that the lack of responses to contralateral regressive motion in bIPS, and its enhanced selectivity to translational optic flow could arise from recurrent interactions between excitatory LPTCs and inhibitory interneurons. For example, excitation provided by H2 to bIPS could be suppressed by indirect, H2-dependent inhibition. To identify possible candidates, we analyzed the predicted neurotransmitter^52^ for bIPS input neurons (**Fig. 4a**). The two main inputs to bIPS are uLPTCrn and H2rn, both of which are GABAergic (**Fig. 4-S1**) and receive prominent inputs from H2 (**Fig. 4b**). The subcellular organization of H2, HS, uLPTCrn and H2rn inputs relative to the root point of bIPS (**Fig. 4-S2a**) suggests that these neurons tend to make synapses closer to the root point than other bIPS inputs (**Fig. 4-S2b-c**). H2rn synaptic inputs onto bIPS tend to be closer to H2 inputs than HS inputs (**Fig. 4-S2d**), often forming microcircuit motifs with H2 (**Fig. 4-S2e, g**) but not with HS cells (**Fig. 4-S2f, h**). Thus, this local connectivity structure could suppress H2-related excitation through H2rns, but it may not fully account for the lack of responses to contralateral regressive motion in bIPS (**Fig. 3-S2c**). Another possibility is that both H2rns and uLPTCrns contribute to this suppression.

**Figure 4:**
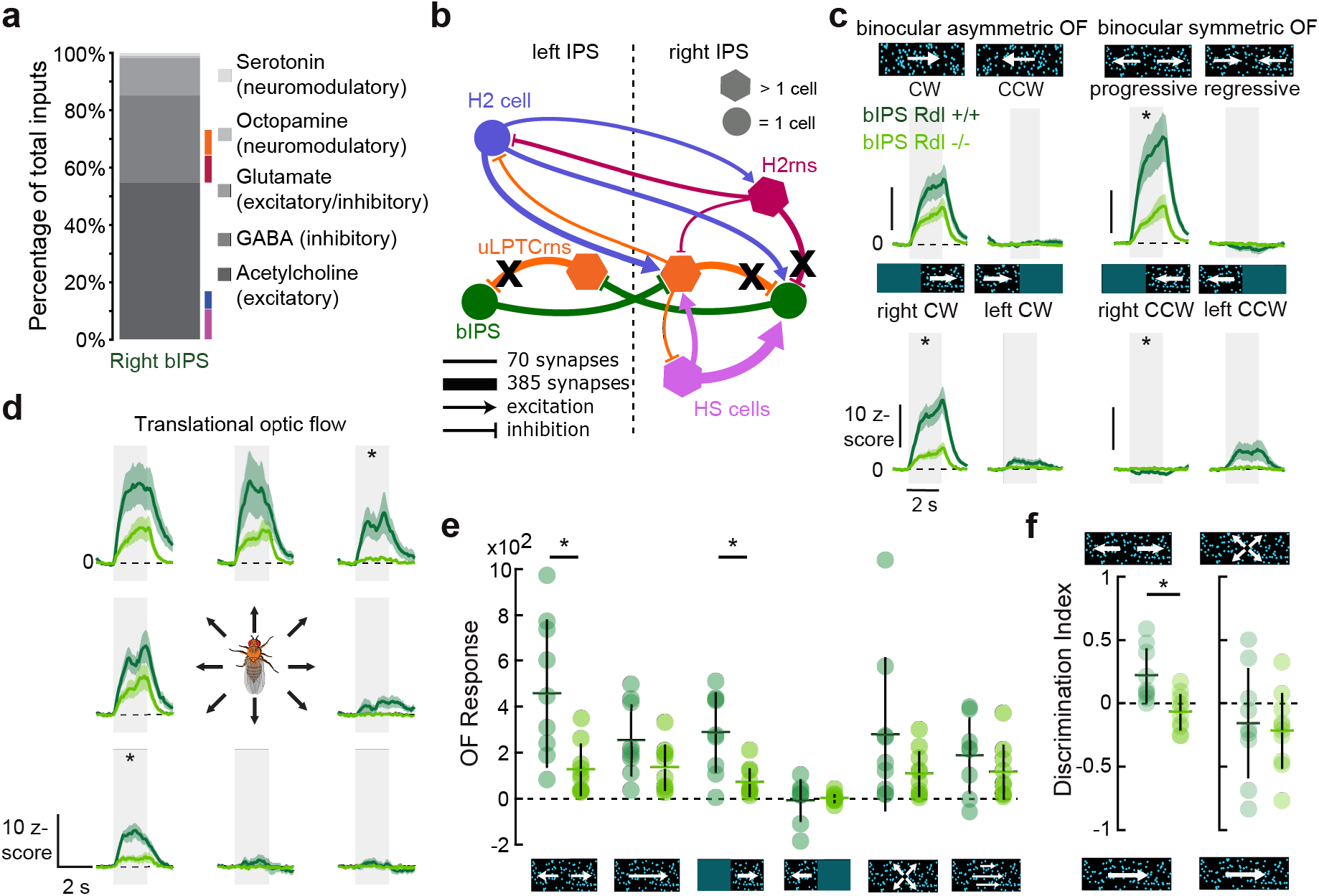
GABAergic inhibition fine-tunes optic flow responses in bIPS. (a) Right bIPS input profile, grouped by their predicted primary neurotransmitter. Neurons shown in b) are highlighted with colored lines. (b) Schematic of the local connectivity of the input and middle layers of the H2-HS network. Inhibitory connections are shown as tee shaped edges. Blocking Rdl receptors in bIPS using FLPStop disrupts fast GABAergic signaling onto bIPS (black crosses). (c) Neural responses of wild-type (dark green) and Rdl-disrupted (light green) bIPS cells to binocular asymmetric (left, rotational) and symmetric (right, translational-like) optic flow (OF). Each trace corresponds to the grand mean ± SEM (shaded area) of z-scored calcium responses obtained across flies. For each fly, mean responses to 5-10 repetitions per stimulus were obtained from a single ROI at the axon terminals. Stimuli with a star indicate area under the curve (auc) responses that are statistically different from wild type bIPS recordings. (d) Same as c), but for translational optic flow. (e) OF responses, the difference between the preferred and nulled direction responses (auc) for horizontal and progressive visual motion. (f) Left, discrimination index between translational-like and rotational optic flow of recorded neurons in c) (see Methods). Each circle represents an individual ROI. Colored horizontal line and black vertical line represents mean and 95% CI. Right, discrimination index between progressive and rotational optic flow. For all panels: bIPS: N=7 flies n=9 ROIs, bIPS Rdl-FLPStop: N=7 flies n=10 ROIs. *P < 0.05; **P < 0.01 ***P < 0.001, Wilcoxon’s rank-sum test.

To test this idea, we perturbed fast-acting GABA signaling in bIPS using the FlpStop genetic strategy^53^. We targeted the Resistance to dieldrin (Rdl) gene^54^ since it is essential for functional GABA_A_ receptors^55,56^ (GABAARs). Overall, bIPS’ responses to optic flow Rdl-FlpStop flies were decreased relative to control flies (**Fig. 4c-e**), including lack of negative calcium responses to non-preferred directions of stimulus motion (**Fig. 4c-d**). Notably, the effect of disrupting GABA_A_ signaling was most evident for translational and translational-like optic flow (**Fig. 4c-e**). This caused the discrimination index between translational-like and rotational optic flow responses to skew towards zero (i.e., equal discriminability), while the discrimination index between translation and rotational optic flow remained unaffected (**Fig. 4f**). These results suggest that the enhanced selectivity to horizontal binocular symmetric motion, as well as the responses to the non-preferred directions of motion are mediated by GABAARs in bIPS. However, contralateral regressive responses that drive H2 activity remain unaffected under Rdl-FlpStop (**Fig. 4c**, “left CW”), indicating that signals from H2 are gated by other mechanisms. Taken together, these findings show that GABAergic inhibition modulates bIPS’ selectivity to binocular symmetric optic flow within the network. We propose that this selectivity may play a key role in enhancing the asymmetries in the activity of left versus right postsynaptic partners, particularly in the sensory context of translational optic flow where the visual input becomes more symmetric (**Fig. 1**).

### Reciprocal inhibitory connections in the middle layer of the H2-HS network promote competition between the left and right sides for steering control

Our Rdl-FlpStop experiments suggest that inhibitory input onto bIPS enhances its responses to binocular symmetric stimuli. Because bIPS is an inhibitory neuron, its selectivity may facilitate the detection of minor binocular asymmetries at the output layer of the network. However, bIPS is embedded in a reciprocal motif with contralateral uLPTCrns (**Fig. 3a**), making it challenging to understand the role of this motif in a dynamic setting like locomotion. Therefore we implemented a phenomenological model agent^33^ constrained by the connectivity structure of the HS-H2 network (**Fig. 5, 5-S1**, see **Methods**), to study the role of bIPS and uLPTCrns on steering control. Our agent processes horizontal motion to guide steering of a virtual fly, like an exploratory real fly^39^. We built a model with four input channels, each containing classical visual motion detectors^28^, and the core cross-inhibitory circuit architecture of the H2-HS network (**Fig. 5a**). The model was tuned to match the real neural data for horizontal motion stimuli (see **Methods**, **Fig. 5-S1a-c** and **5b**).

**Figure 5:**
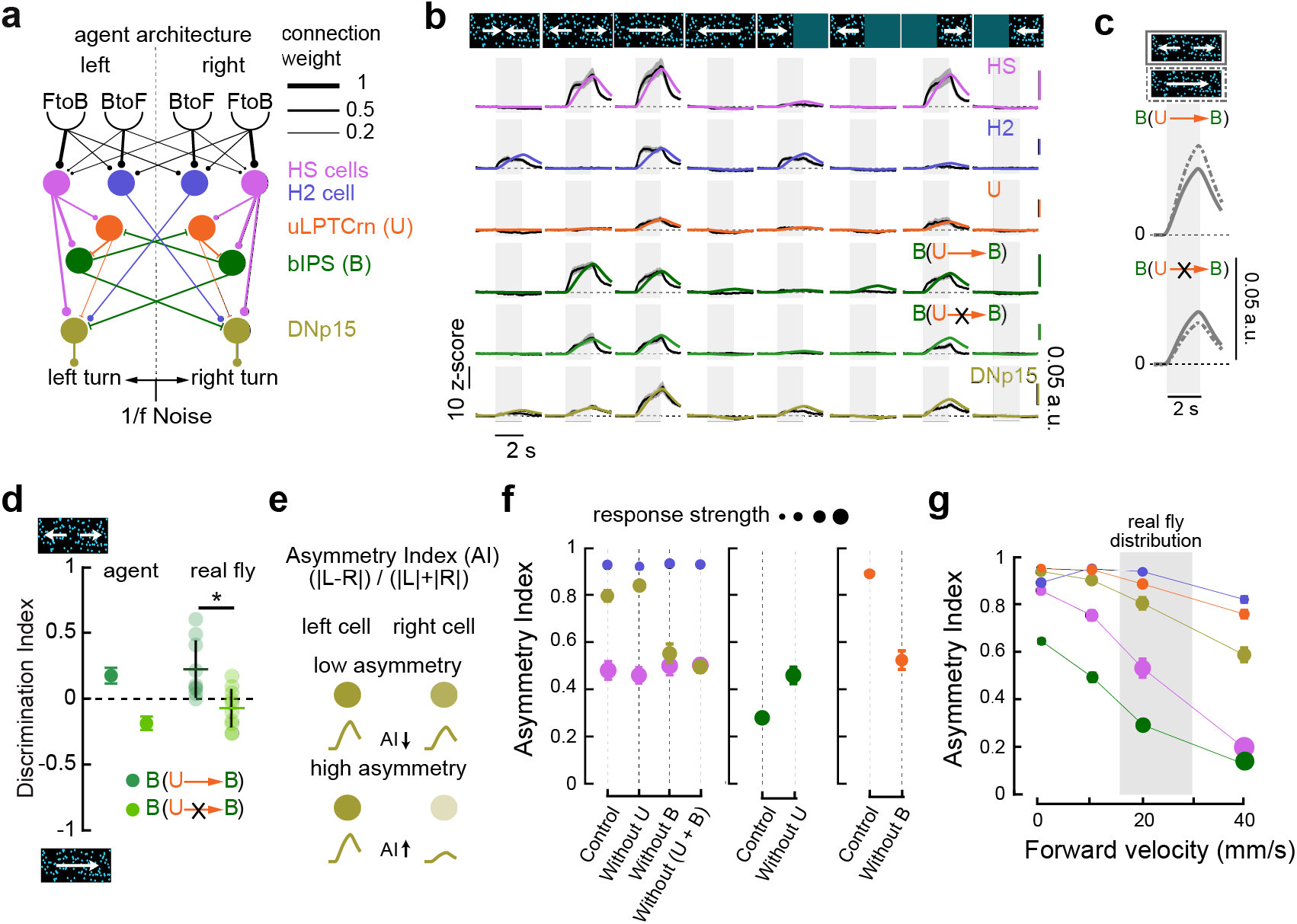
A lateral disinhibitory motif enhance asymmetries in neural activity across the right versus left hemisphere. (a) Schematic of the agent architecture: two sets of Reichardt detectors tuned to front-to-back (FtoB) or back-to-front (BtoF) motion detect the optic flow generated by a simulated walking fly (See Methods, Fig. 5-S1). The detected visual motion is processed by a network of nodes that mimic the core network motif of H2-HS network (**Fig. 3a**). The weights that connect the nodes are tuned based on a subset of measured visual responses (Fig. 5-S1). (b) Simulated and observed visual responses to monocular and binocular patterns of optic flow. Responses were matched to the maximal observed response. Observed responses are replotted from Fig. 3-S2 and **Fig. 4c** (c) Visual response of modeled bIPS to symmetric or asymmetric binocular optic flow, and with or without a functional input from uLPTCrns. (d) Left: discrimination index of modeled and observed bIPS with or without a functional input from uLPTCrns. Right: observed bIPS’ discrimination index with or without functional GABAergic input (replotted from **Fig. 4f**). (e) Definition of the Asymmetry Index (AI) and two example conditions with low and high AI values. (f) Asymmetry index of simulated neurons (colored) in the full model (Control), under bIPS silencing (Without B), under uLPTCrn silencing (Without U) and under both bIPS and uLPTCrn silencing (Without U+B). The dot size corresponds to the magnitude of the neural response. (g) Asymmetry index of simulated neurons (colored) in the full model varying the forward velocity of the agent. The shaded area corresponds to the range of velocities observed for forward runs in real exploratory flies (Cruz et al. 2021)^39^.

We tested the effect of removing the inhibitory input to bIPS in the model by removing ipsilateral uLPTCrns inputs. This manipulation abolished the enhanced responses of bIPS to binocular symmetric stimuli (**Fig. 5c**) and affected the discriminability of the modeled cell to rotational vs translational-like stimuli, like in the real fly (**Fig. 5d**). Based on these results, we concluded that this simplified core model architecture, combining direct excitation with lateral disinhibition, can replicate the visual responses of the input and middle layers of the H2-HS subnetwork.

Next, we tested the contribution of the lateral inhibition provided by bIPS to the output responses of the network in the context of course control. This agent fly compares the activity of a left and right pair of DNs, akin DNp15, to impose steering control (**Fig. 5e**). Removing bIPS from the model reduced the asymmetry between left and right DNp15 cells, leading to a reduced ability to discriminate asymmetries in binocular horizontal motion during translations at high speed (**Fig. 5f**). Removing both uLPTCrns and bIPS further reduced the asymmetry between left and right DNp15 activity down to the level of HS. Like DNp15, activity of uLPTCrn cells across the two eyes also became more symmetric upon removing bIPS (**Fig. 5f**). Conversely, removing uLPTCrn cells made bIPS less sensitive to horizontal symmetric binocular motion (**Fig. 5c**). Thus, the reciprocal inhibitory and lateral interactions within the middle layer of the H2-HS network support enhanced sensitivity to differences between the eyes at the output layer in the context of translations. Indeed, the asymmetry in DNp15 and uLPTCrn cells remained high in ranges of forward velocities typically observed in exploratory flies, and we propose that this is aided by the increased symmetric activity in bIPS as a function of translation speed (**Fig. 5g**).

To test the direct contribution of bIPS to steering in the context of highspeed translation, we allowed our agent fly to freely explore a virtual arena where the fly’s trajectory was determined by bouts of straight walking (forward runs) and rapid turns (saccades) away from the walls, similar to real flies exploring a comparable environment (**Fig. 6a, 5-S1d,** see **Methods**). As expected by model design, ‘silencing’ individual DNp15 cells predicted the angular bias observed in real flies under individual DNp15 silencing (**Fig. 6b**, see also **Fig. 1i**). Silencing individual bIPS neurons in the model induced biases during forward runs and performing the similar perturbation in real flies lead to similar results (**Fig. 6c, 6-S1a**). This finding demonstrated that asymmetries between left and right bIPS activity, consistent with the neuron’s translational optic flow selectivity (**Fig. 3-S3**), are sufficient to bias the course direction of walking flies (**Fig. 6c**). Consistent with the model, bilateral manipulations of activity of bIPS did not affect the course of forward runs of real flies (**Fig. 6d**).

**Figure 6:**
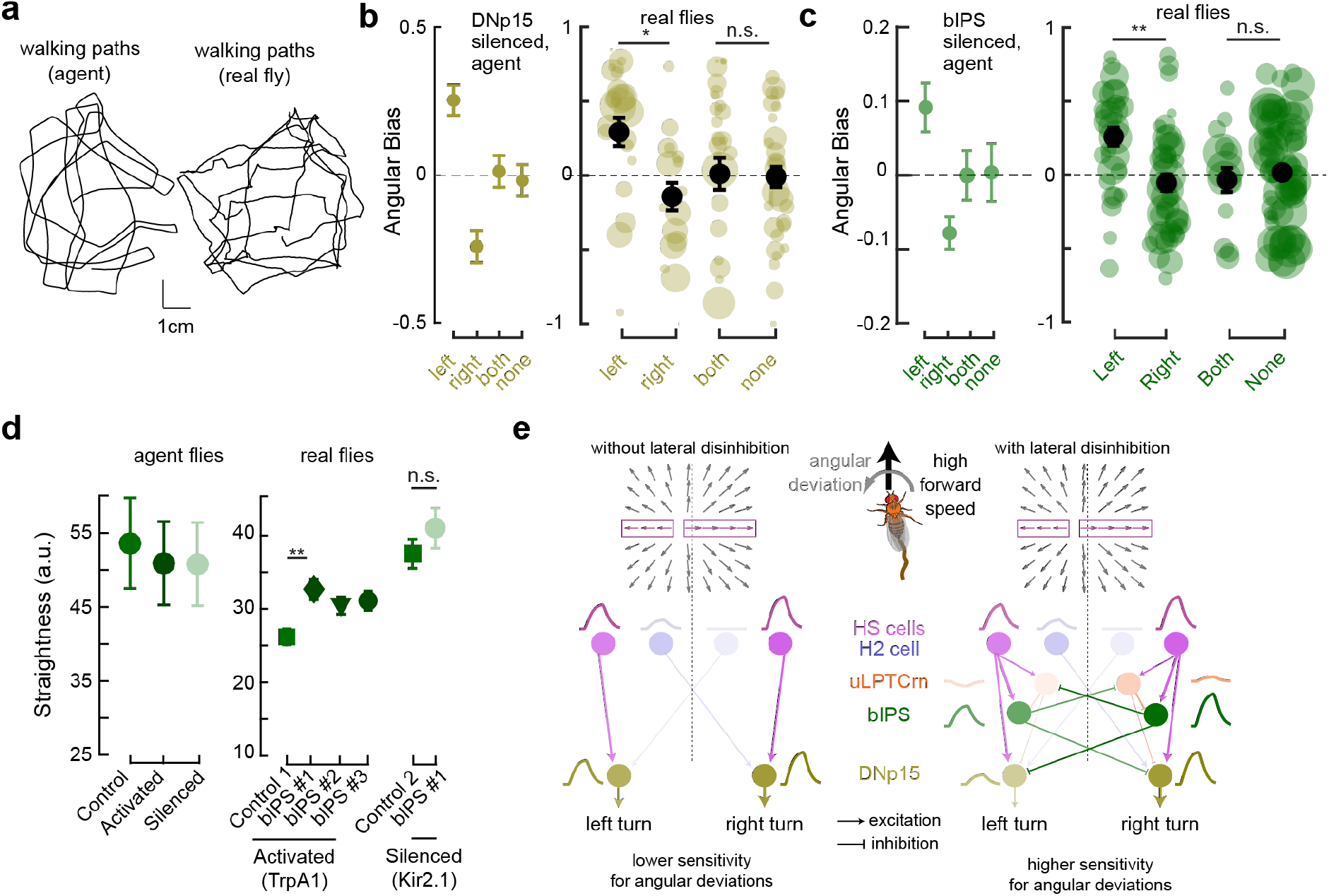
bIPS plays a role in steering control during high-speed walking. (a) Example walking paths of the agent (left) and the real fly (right) under prominent visual feedback. (b) Simulated (left) and real fly (right) angular bias during forward runs (see Methods) with DNp15 activity perturbed bilateral or unilaterally. Transparent dots depict mean bias of individual flies, black dots represent grand mean per genotype, weighted by the amount of walking bouts (dot size). Error bars represent SEM. Number of flies per condition, none n=25; left n=22; right n=22; both n=31. (c) Same as b), but for bIPS. Number of flies per condition, none n=65; left n=26; right n=38; both n=13. For (b) and (c): *p < 0.05; **p < 0.01 ***p < 0.001, Wilcoxon’s rank-sum test with Bonferroni correction for multiple comparisons. (d) Left, simulated straightness of the agent during forward runs (see Methods) with control (green), bilaterally activated (dark green) or silenced (light green) bIPS cells. Right, straightness of control flies (green), or flies with bilaterally activated (dark green) or silenced (light green) bIPS cells. Number of flies per condition, empty control>TrpA1 n=25; bIPS#1>TrpA1 n=27; bIPS#2>TrpA1 n=25; bIPS#3>Tr-pA1 n=29; empty control>Kir2.1 n=22; bIPS#1>Kir2.1 n=23. (e) Summary of the findings. The middle layer of the H2-HS network supports steering by enhancing the difference in the activity levels of the output layer across hemispheres. This function is driven by a competitive disinhibitory motif that is well posed to be recruited at high-speed walking. In this condition, binocular optic flow may contain prominent translational components, possibly overshadowing rotational cues that are critical for course control. We propose that the competitive disinhibition at the middle layer plays an important role in modulating the output layer in a continuous and dynamic manner, leading to a more robust estimate of the deviations from a straight course.

In summary, our findings support the idea of a central multilayer network that monitors rotational components of optic flow during translation and that amplifies asymmetric activity between right- vs. left outputs by lateral, competitive disinhibition, thereby increasing the rotational signal even in the context of prominent translational optic flow (**Fig. 6e**). This reciprocal, lateral inhibition therefore supports robust optic flow processing and increases the performance of networks contributing to course control, two functions that are especially important while walking at high speed.

## Discussion

In this study we investigated how visuomotor circuits discriminate sensory signals in challenging circumstances imposed by an animal’s own locomotion. Through a combination of full-brain EM reconstruction, modeling, optical imaging and behavior experiments, we discovered a multi-layer network in the *Drosophila* central brain responsible for robust optic flow processing and steering of exploring flies based. The first layer consists of optic flow sensitive visual projection neurons, which form two networks with distinct sets of interneurons and outputs. Both networks exhibit specific and recurrent connections between LPTCs and a set of GABAergic inhibitory interneurons. Furthermore, there are critical reciprocal and lateral interactions among the middle layer and across the hemispheres. The output layers receive direct and indirect connections from the input layer, with the indirect pathway strongly influenced by lateral disinhibition of the middle layer. This configuration allows the output layer to generate a robust estimate of the fly’s rotations, as indicated by their visual consequence. Our findings unveils a network motif of competitive disinhibition^4^ that enhances the detection of asymmetries in optic flow across the eyes, enabling course control even in the presence of highly symmetric bilateral visual motion cues (**Fig. 6e**).

### Parallel networks process visual feedback in the fly brain

Until now, the few identified neck motor and descending outputs of HS and VS cells have been shown to connect to either HS or VS cells^30,37,57,58^ (CNMN6^59^ being the single exception). As such, HS and VS cells have been proposed to contribute to movement control through largely parallel pathways. However, if downstream circuits of these LPTCs do not converge, how can flies accurately estimate their self-movement when their natural maneuvers during locomotion involve multiple axes of rotation and translation? By identifying the major synaptic partners of HS, H2 and VS cells in the central brain (**Fig. 2**) we have found that these LPTCs operate largely in parallel, connecting to distinct sets of descending, neck motor and central brain output neurons. However, HS, H2 and VS cells also converge onto anatomically distinct classes of GABAergic interneurons in the posterior slope. These interneurons recurrently connect to different and complementary combinations of HS, H2 and VS, suggesting the integration of orthogonal axes of optic flow at the central brain (**Fig. 3**). We propose that this partially overlapping recurrent motif enables accurate processing of naturalistic optic flow, modulating the concerted activity of LPTCs based on distinct visuomotor conditions, and other internal contexts. Indeed, the identified inhibitory classes receive inputs from motor and premotor areas of the fly CNS, including the VNC, PS, and GNG (**Fig. 2-S3**). This feedback structure indicates strong processing of multimodal signals to differentially modulate the activity of LPTCs and their postsynaptic outputs. These outputs likely contribute to different motor programs and higher-order functions. For example, outputs of the H2-HS network such as DNa02^47^, DNp15^30,37^ and VCNMN^57,60,61^ have all been proposed to control rotations of the body and the head along the yaw axis, while outputs of the VS subnetwork such as DNp20 and ocelli have been implicated in gaze stabilization control along the roll axis^60,62,63^. Each network project to neck-, leg-, and haltere-related regions in the VNC, implicating LPTCs in the control of head and body movement independently but likely within the context of head-body coordinated (gaze) motor programs occurring during exploratory walking^39,64^ and flight^65^. Additionally, output pathways that bridge LPTC activity with higher-order brain regions, such as lateral accessory lobe and central complex via PS047, can provide accurate internal self-motion estimates for heading perception and path integration during navigation^66–69^.

### GABAergic inhibition modulates asymmetries in visual motion processing

Three cell types in the intermediate layers of H2-HS subnetwork, namely bIPS, uLPTCrn and H2rn cells, exhibit optic flow sensitivities that are only partially explained by their upstream LPTCs. This finding indicates that additional inputs influence their optic flow sensitivity (**Fig. 3**). We have identified one group of such inputs: reciprocal connections among the three cell types. These connections play a critical role in transforming optic flow signals within the middle layer. For example, when fast acting GABAergic inputs to bIPS are disrupted, the negative response caused by non-preferred directions of optic flow is eliminated (**Fig. 4**). This effect is similar to the modulation of non-preferred visual motion responses in the fly early visual system^70,71^. Furthermore, the same perturbation reduces bIPS’ selectivity to symmetric horizontal optic flow. Our constrained model suggests that this reduction is partially caused by the inhibitory population of uLPTCrns, thereby likely affecting the enhanced sensitivity to rotational optic flow by the output of the network. However, and contrary to the expectations based by the connectivity structure, rapid GABAergic inputs do not influence bIPS’s responses to regressive, back-to-front optic flow. Although bIPS receive strong connections from H2, signals from these synapses are not transmitted under non-behaving conditions. One possibility is that the GABAergic population of H2rns controls this signaling through other GABA receptors in bIPS. Alternatively, the gating of regressive signals from H2 may involve other mechanisms, including neuromodulation. Further investigation will explore the conditions under which bIPS respond to regressive information, which may render the neuron more sensitive to asymmetric optic flow stimuli. Nevertheless, this finding highlights the importance of combining EM connectivity with physiology to uncover context-dependent modulation of brain circuits.

### Reciprocal inhibition in optic flow networks supports corrective steering

A straightforward circuit for angular course control based on optic flow can be created by connecting two rotational optic flow detectors (e.g., HS cells) separately to two rotational actuators (e.g., DNp15). Such a system performs well when there is a significant difference in activity levels between the two sensors. However, it performs poorly in detecting rotations when the animal is moving forward at high speeds, which is surprisingly the condition where HS cells contribute the most to corrective steering^35,36^. Furthermore, locomotion requires continuous adjustments due to internal sources of noise, such as muscular and sensorimotor noise, as well as uncertainties in the environment. These adjustments cause the optic flow to be non-stationary, even when the fly intends to maintain a fixed course^72^.

We propose two critical roles for the inhibitory motifs identified in the H2-HS network. Firstly, it allows comparing the activity state of LPTCs across the two hemispheres and provides cross-inhibition to silence competing inhibitory neurons, leading to a binary choice of turning left or right. Similar circuit motifs with cross-inhibitory neurons in the hindbrain of Zebrafish^73^ and the vertebrate superior colliculus^74^ bias angular movements in opposite directions when activated or inhibited asymmetrically. Our experiments with bIPS yield analogous results (**Fig. 6c**). Secondly, the inhibitory motifs can adjust the gain of LPTCs during highspeed locomotion, allowing them to flexibly respond to changes in external and internal conditions. Notably, bIPS provides prominent contralateral inputs to inhibitory feedback CH cells, which project from the posterior slope back the lobula plate. CH cells receive more than 450 synapses from bIPS (**Fig. 2-S1**). It is possible that bIPS corresponds to the previously hypothesized *Horizontal unknown* cell, which provides binocular information to CH cells through contralateral inhibition^75^, thus playing a major role in integrating horizontal motion from both eyes. Given its selectivity and inputs from motor and premotor regions, bIPS is well suited for monitoring the fly’s motor state, modulating descending neurons, feedback pathways, and central pathways during periods of straightforward walking. This class of cells also instructs optic flow processing, self-motion estimation and steering in the presence of angular drifts. Therefore, bIPS contributes to the modulation of optic flow processing in a multilayer manner.

### Estimation of self-motion from the eyes down to premotor and motor centers

Optic flow-sensitive neurons have long been proposed to be integral components of vital visuomotor circuits that enable animals to accurately perceive heading and control their gaze and steering. In flies, while we have made substantial progress in understanding how light reaching the photoreceptors are processed to detect visual motion and stimulate HS, H2, VS cells and other LPTCs^28,76^, our knowledge of how downstream pathways are organized and process optic flow has remained far more limited. With the comprehensive description of the HS, H2 and VS networks outlined here, and the visual and behavioral analysis of neurons within these networks, we now possess a framework to establish mechanistic links between visual activity in these networks and the ongoing locomotion control of a freely moving fly. Given the common characteristics of optic flow and associated visual neurons processing across different animal species^77,78^ and the universal constrains on locomotive systems, the findings reported here have the potential to shed light on how analogous systems in other animals are functionally organized to process naturalistic optic flow in support of navigation.

## Acknowledgements

We thank Ruchi Parekh and Janelia CAT for introducing us to FAFB and training us in EM tracing. Kaylynn Coates and Feng Li for reviewing some of our manually traced neurons in FAFB. Stephen Huston for early tracing of VCNMN and some cLPTCrn cells with Shahrozia Imtiaz and for helping identify neck motor neurons. Anna Li and Rachel Wilson for their help in reconstructing DNa02. Shigehiro Namiki and Gwyneth Card for their help in identifying descending neurons. Jens Goldammer for identifying potential split-Gal4 candidates for H2rn cells and helping identify S-Neurons. Hideo Otsuna, Greg Jefferis and Phillip Schlegel for introducing us to their tools for computational neuroanatomy and for helping with EM data analysis. Arthur Zhao for his input on LPTC naming, response predictions, and feedback on the manuscript. Nils Eckstein and Jan Funke for sharing neurotransmitter predictions for LPTC partners prior to publication. Gaby Maimon and Cheng Lyu for sharing sytGCaMP7f flies prior to publication. Marion Silies for sharing the UAS-GCaMP6f, UAS-FLP recombinant line. Ece Sönmez for help with testing Split-Gal4 combinations and help with validating FlpStop recombinants. CR Fly Platform for assisting fly stock generation, maintenance, and GABA stainings. Lalanti Venkatasubramanian for her help with the initial bIPS>TrpA1 behavior experiments. Terufumi Fujiwara for help with finding and testing potential Split-Gal4 combinations for H2rn and uLPTCrn and help with building the 2-photon calcium imaging set up. André Marques and Wynne Stagnaro for sharing unpublished work about multisensory processing in LPTCrn and bIPS cells. Past and present Chiappe Lab members for useful discussions and feedback on experiments and the manuscript. We thank the Princeton FlyWire team and members of the Murthy and Seung labs, as well as members of the Allen Institute for Brain Science, for development and maintenance of FlyWire (supported by BRAIN Initiative grants MH117815 and NS126935 to Murthy and Seung). We also acknowledge members of the Princeton FlyWire team and the FlyWire consortium for neuron proofreading and annotation. This work was supported by the Champalimaud Foundation and the research infrastructure Congento, LISBOA-01-0145-FEDER-022170. MECh is supported by European Research Council Starting Grant ERC-2017-STG-759782 537.

## Author Contributions

In alphabetical order:

**<u>Davi D Bock</u>**: Generation of FAFB dataset and allowing access pre-publication. Resources & Data Curation.

**<u>Margarida Brotas</u>**: Reconstruction and review of 1^st^ and 2^nd^ order LPTC partners and cross-matching LPTC partners across EM datasets. Investigating differences in anatomy, connectivity, and neurotransmitter profile among LPTC partners.

**<u>M Eugenia Chiappe</u>**: Conceptualization. Methodology. Supervision. Resources. Funding acquisition. Writing – original draft. Writing – Review & Editing. Visualization.

**<u>Tomás Cruz</u>**: Design, running and analysis of DNp15>hsFLP,Kir experiments. Provide input, feedback and initial scripts to parse and analyze behavior data. Behavioral agent simulations with the modeled network.

**<u>Mert Erginkaya</u>**: Conceptualization. Reconstruction and review of LPTC synaptic partners and some LPTCs in FAFB. Reconstruction and review of 2^nd^ order inputs/outputs of LPTC networks. Cross-matching LPTC partners across EM datasets. Analysis of EM anatomy & connectivity. Generation of all transgenic lines. GABA stainings (with the help of Fly Platform). Designing, performing, and analyzing calcium imaging experiments. Design and analysis of behavior experiments. Visualization of data and writing the paper.

**<u>Aljoscha Nern</u>**: Generation of transgenic lines that label LPTCrns. Re-injection of uLPTCrn Gal4 line. Input for split candidates for bIPS. EM-LM cell identification and naming for several cell types.

**<u>Michael B Reiser</u>**: Conceptualization. Reconstruction of LPTCs in FAFB and proofreading LPTCs in FlyWire and Hemibrain (by Reiser Lab tracers). Naming LPTCs. Help with generating optic flow stimulus patterns in LED arena (Software, from Matthew Isaacson). Regular feedback on EM tracing status & priorities, and feedback on calcium imaging experiments.

**<u>Kathrin Steck</u>**: Reconstruction and review of LPTC partners in FAFB. Reconstruction and review of 2^nd^ order inputs/outputs of LPTC networks.

**<u>Filipa Torrão</u>**: Investigating differences in connectivity among LPTC partners. Analysis of local connectivity structure within bIPS dendrites (**Fig. 6-S2**)

**<u>Nélia Varela</u>**: Generation of transgenic lines that label bIPS, H2rn, H2, HS. Immunostainings and images for several transgenic lines. Performing behavioral experiments. Immunostainings for individual bIPS>hsFLP,Kir and DNp15>hsFLP,Kir and MCFO flies.

## Inclusion and Diversity Statement

While citing references scientifically relevant for this study, we worked to promote gender balance in our reference list as much as possible

## Tables

**Supplementary Table 1:** Cross-matching of LPTC outputs across EM dataset.

| H2 Output Cells | FlyWire rank | FAFB rank | Hemibrain rank |
| --- | --- | --- | --- |
| PS047 | 3 | 1 | 3 |
| VCH | 1 | 2 | 2 |
| DCH | 2 | 3 | 1 |
| DNp15 | 4* | 4 | 4 |
| bIPS | 5 | 5 | 5 |
| CNMNx | 34* | 6 | 11** |
| DNb03 | 6 | 7 | 7 |
| DNb03 | 8 | 8 | 8 |
| DNg41 | 9 | 9 | 6** |
| uLPTCrn | 11 | 10 | 9 |
| PS126 | 10 | 12 | 28 |
| PS099 | 7 | 31 | 48 |
| PS081 | 14 | 16 | 10 |
| HS Output Cells | FlyWire rank | FAFB rank | Hemibrain rank |

| bIPS | 1 | 1 | 2 |
| --- | --- | --- | --- |
| DNp15 | 2* | 2 | 1 |
| PS047 | 4 | 3 | 3 |
| CNMNx | 21* | 4 | 4** |
| VCNMN | 5 | 5 | 7** |
| DNa02 | 3 | 6 | 9 |
| uLPTCrn | 10 | 7 | 5 |
| IVLPT8 | 8 | 8 | 35 |
| uLPTCrn | 16 | 9 | 6 |
| uLPTCrn | 13 | 10 | 10 |
| PLP195 | 6 | 11 | 29 |
| DNgX | 7 | 26 |  |
| DNa04 |  |  | 8 |
| VS Output Cells | FlyWire rank | FAFB rank | Hemibrain rank |
| DNp20 | 1* | 1 | 2 |
| DNb06 | 2 | 2 |  |
| cLPTCrn | 4 | 3 | 10 |
| CNMN6 | 6 | 4 | 6 |
| CNMN7 | 31 | 5 |  |
| cLPTCrn | 7 | 6 | 20 |
| uLPTCrn | 40 | 7 | 5 |
| PS126 | 3 | 8 | 7 |
| S-Neuron | 45 | 9 | 8 |
| S-Neuron | 38 | 10 | 9 |
| DNge043 | 5 | 48 | 3 |
| DNp22 | 8 | 70 |  |
| PS135 | 9 | 37 | 1 |
| cLPTCrn | 10 | 12 | 16 |
| PS169 | 48 |  | 4 |

\* FlyWire segmentation or synapse predictions are of low quality
\*\* Uncertain hemibrain matching, since it is based on a small neural arbor cut off at the limits
of the dataset

**Supplementary Table 2:** List of neurons within the LPTC central brain network.

| # | FAFB ID | FlyWire segment ID* | FlyWire soma coordinates | Hemibrain ID | Hemibrain cell type |
| --- | --- | --- | --- | --- | --- |
| 1 | Right uLPTCrn<br>2918805 PG ME | 720575940628599516 | 114786, 73911, 1623 |  | PS070-PS074 |
| 2 | Right<br>uLPTCrn<br>11965240<br>ME | 720575940631296543 | 113632,<br>74374, 1687 |  | PS070-PS074 |
| 3 | Right DNg41<br>15463855<br>MB | 720575940617611643 | 99290,<br>75988, 3794 | 5813024629** |  |
| 4 | Right<br>uLPTCrn<br>2549438 ME | 720575940631333955 | 109930,<br>75338, 1666 |  | PS070-PS074 |
| 5 | Right<br>uLPTCrn<br>2544668 ME | 720575940612144047 | 112472,<br>73317, 1957 |  | PS070-PS074 |
| 6 | GNG neuron<br>2948699 ME | 720575940648969337 | 121455,<br>91088, 2979 |  |  |
| 7 | DCH neuron<br>(right)<br>1077175<br>ZSG-GA | 720575940640039989 | 131955.27,<br>53517.50,<br>1223.00 | 1466485353 | DCH |
| 8 | VCH neuron<br>(right)<br>1078536<br>GA-ZSG | 720575940627138562 | 134785,<br>53191, 958 | 5813024201 | VCH |
| 9 | Putative<br>CNMN<br>5218081 ME | 720575940622008560*** | 105319,<br>78004, 4717 | 2180780757** |  |
| 10 | VCN002 DB | 720575940635776760*** | 124703,<br>91385, 3579 | 2428058771** |  |
| 11 | DNa02_AL | 720575940629327659 | 101150,<br>42526, 2117 | 1140245595 | DNa02 |
| 12 | neuron<br>7828867 BG | 720575940621564329 | 104740,<br>50998, 5510 |  | PLP195 |
| 13 | DNgX<br>999559 ME | 720575940614730914 | 105915,<br>76095, 4569 | 1874217622** |  |
| 14 | Putative<br>Right<br>cLPTCrn<br>410927 SI | 720575940642061709 | 112863,<br>73654, 2042 |  | PS079 |
| 15 | Right<br>cLPTCrn<br>410809 SI | 720575940617997396 | 113277,<br>73467, 2192 |  | PS079 |
| 16 | Right<br>bLPTCrn<br>416268 ME | 720575940608760110 | 112920,<br>74883, 1730 |  | PS078 |
| 17 | Right<br>bLPTCrn<br>986687 ME | 720575940620555696 | 111749,<br>73943, 2041 |  | PS078 |
| 18 | Right<br>uLPTCrn<br>2500746 ME | 720575940606873522 | 112559,<br>74568, 1653 |  | PS070-PS074 |
| 19 | Right<br>uLPTCrn<br>1002426 ME | 720575940647708537 | 114241,<br>73636, 1682 |  | PS070-PS074 |
| 20 | Right DNp15<br>5224930 ME | 720575940610301413*** | 120043,<br>60033, 5393 | 5813078134** | DNp15 |
| 21 | Right bIPS<br>neuron<br>3029744 ME | 720575940622581173 | 125645,<br>88652, 4814 | 1932497911 |  |
| 22 | Right PS047<br>16769498<br>ME | 720575940638306163 | 98755,<br>67802, 2032 | 1621806893 | PS047 |
| 23 | Left bIPS<br>neuron<br>969206 ME | 720575940623618708 | 141214,<br>90010, 4743 | 1963869636 |  |
| 24 | H2 neuron<br>(left)<br>5232903 ME | 720575940632427603 | 164810,<br>54592, 5192 | 5813078454 | H2 |
| 25 | Right DNb03<br>8416446530<br>ME | 720575940620776880 | 105915,<br>58345, 1981 |  | DNb03 |
| 26 | Right H2rn<br>11956981<br>ME | 720575940621865991 | 127153,<br>91186, 3928 |  |  |
| 27 | Right DNb03<br>413171 MB | 720575940634163682 | 107706,<br>58941, 2096 |  |  |
| 28 | Right H2rn<br>11930922<br>ME | 720575940622082451 | 128029,<br>91037, 3888 |  |  |
| 29 | Right H2rn<br>9730354 ME | 720575940624756836 | 126229,<br>90972, 3889 |  |  |
| 30 | Right H2rn<br>8438890 ME | 720575940617566769 | 127758,<br>91886, 3927 |  |  |
| 31 | Right H2rn<br>8178074 ME | 720575940642076576 | 127930,<br>91367, 4018 |  |  |
| 32 | Putative<br>VSN 851433<br>ZSG | 720575940624931564 | 89397,<br>61348, 6667 | 5813033533 | VS |
| 33 | Putative<br>VSN 804093<br>ZSG | 720575940633923298 | 82426,<br>57153, 6848 | 1558230909 | VS |
| 34 | neuron<br>5207415 ME | 720575940613940074 | 83118,<br>55640, 4101 | 1221457679 | IVLPT8 |
| 35 | Putative<br>HSN 830794<br>ZSG | 720575940628031249 | 91612,<br>60674, 6567 | 2211443902 | HSN |
| 36 | Putative<br>HSE 827035<br>ECS ZSG | 720575940642723981 | 98857,<br>55902, 5783 | 1807537598 | HSE |
| 37 | Putative<br>HSS 408972<br>SJH ZSG | 720575940622312965 | 94341,<br>61845, 6407 | 2179731270 | HSS |
| 38 | VSN-Like<br>982898 ZSG | 720575940626477498 | 86531,<br>61966, 6790 | 5813024262 | VS |
| 39 | putative<br>VSN 793033<br>ZSG | 720575940615269794 | 82514,<br>59151, 6836 | 1868619183 | VS |
| 40 | Putative<br>VSN 815777<br>ZSG | 720575940622831740 | 84905,<br>55521, 6704 | 5813025091 | VS |
| 41 | Putative<br>VSN 852994<br>JLME ZSG | 720575940633017939 | 82465,<br>56046, 6799 | 1557885051 | VS |
| 42 | Putative<br>VSN 807402<br>ZSG | 720575940626457406 | 85387,<br>60389, 6811 | 1868255513 | VS |
| 43 | putative<br>VSN 804540<br>ZSG | 720575940605688492 | 84584,<br>57582, 6777 | 1992161820 | VS |
| 44 | Right S-Neuron<br>4906099 ME | 720575940629666971 | 126241,<br>28340, 3824 |  | OCG07 |
| 45 | Right S-Neuron<br>4905400 ME | 720575940625314547 | 127007,<br>30053, 4080 |  | OCG07 |
| 46 | Right S-Neuron<br>4880581 ME | 720575940609826499 | 127788,<br>30884, 4072 |  | OCG07 |
| 47 | Right<br>DNOVS1<br>9937342 ME | 720575940629390123*** | 115510,<br>45447, 4960 | 1745333830** | DNp20 |
| 48 | Right DNb06<br>9857697 ME | 720575940637308605 | 126678,<br>68176, 1317 | 1715999506 | DNb06 |
| 49 | Right PS126<br>4905394 ME | 720575940629701484 | 131516,<br>30646, 3039 | 1591156798 | PS126 |
| 50 | Left<br>bLPTCrn<br>9157628 ME | 720575940631876577 | 149420,<br>77362, 1632 |  | PS078 |
| 51 | Right<br>bLPTCrn<br>17646150<br>ME | 720575940624461543 | 113818,<br>75477, 1710 |  | PS078 |
| 52 | Right<br>bLPTCrn<br>869476 ME | 720575940628284931 | 113237,<br>74767, 1898 |  | PS078 |
| 53 | Putative<br>Right<br>bLPTCrn<br>1934541 ME | 720575940605344434 | 113968,<br>76236, 1733 |  | PS078 |
| 54 | Cervical<br>Nerve<br>Neuron<br>1337778 SI | 720575940633404627 | 118851,<br>65265, 5421 |  |  |
| 55 | Cervical<br>Nerve<br>Neuron<br>1167859 SI | 720575940621583286 | 119957,<br>62874, 5368 |  |  |
| 56 | Left<br>bLPTCrn<br>9369685 ME | 720575940610505881 | 148617,<br>76818, 1461 |  | PS078 |
| 57 | Right<br>cLPTCrn<br>968975 ME | 720575940625557069 | 110736,<br>74923, 2028 |  | PS079 |
| 58 | Left<br>cLPTCrn<br>414149 ME | 720575940607991819 | 146347,<br>76067, 1244 |  | PS079 |
| 59 | Left<br>cLPTCrn<br>2725485 ME | 720575940629929273 | 147252,<br>76791, 1398 |  | PS079 |
| 60 | Left<br>cLPTCrn<br>1006244 SI | 720575940612306641 | 146666,<br>75707, 1607 |  | PS079 |
| 61 | Right<br>cLPTCrn<br>414194 ME | 720575940620867254 | 112846,<br>70948, 2282 |  | PS079 |
| 62 | Right<br>cLPTCrn<br>1051520 SI | 720575940629267531 | 111884,<br>74761, 2203 |  | PS079 |
| 63 | Left<br>cLPTCrn<br>967831 ME | 720575940624324708 | 147231,<br>75756, 1219 |  | PS079 |
| 64 | Putative<br>Right<br>cLPTCrn<br>9336219 ME | 720575940631645009 | 111149,<br>73533, 1804 |  | PS079 |
| 65 | Left<br>cLPTCrn<br>4881422 ME | 720575940626420220 | 146247,<br>75467, 1279 |  | PS079 |
| 66 | Right<br>cLPTCrn<br>969249 ME | 720575940620263088 | 111942,<br>73451, 1739 |  | PS079 |
| 67 | Left<br>cLPTCrn<br>952121 ME | 720575940616362123 | 148068,<br>76482, 1602 |  | PS079 |
| 68 | Left<br>cLPTCrn<br>412520 ME | 720575940621845119 | 147390,<br>76088, 1339 |  | PS079 |
| 69 | Right<br>cLPTCrn<br>2560372 ME | 720575940628688246 | 113125,<br>72872, 1901 |  | PS079 |
| 70 | Right<br>cLPTCrn<br>411566 ME | 720575940619872751 | 113686,<br>72642, 2212 |  | PS079 |
| 71 | Left<br>cLPTCrn<br>1027141 SI | 720575940635634680 | 145378,<br>76025, 1378 |  | PS079 |
| 72 | Right<br>cLPTCrn<br>968122 ME | 720575940621147425 | 114031,<br>73227, 1620 |  | PS079 |

\* As of 230422
\*\* Uncertain hemibrain matching, due to incomplete hemibrain skeleton
\*\*\* Bad or incomplete segmentation in FlyWire

**Supplementary Table 3:** List of fly lines used in each figure.

| Figure(s) | Panel(s) | Target cell type(s) | Genotype(s) |
| --- | --- | --- | --- |
| 1 | d, e | DNp15 | w<sup>1118</sup>; R11E07-p65.AD / 20XUAS-jGCaMP7f;<br>R77F05-Gal4.DBD / 20xUAS-6XmCherry;<br>13XLexAop-jGCaMP7f; R77F05-LexA; +; |
| 1 | f, g | DNp15 | hsFLP; R11E07-p65.AD / +; R77F05-Gal4.DBD /<br>10XUAS>Stop>eGFPKir2.1; |
| 1-S1 |  | DNp15 | w<sup>1118</sup>; R11E07-p65.AD / 20XUAS-jGCaMP7f;<br>R77F05-Gal4.DBD / 20XUAS-6XmCherry;<br>13XLexAop-jGCaMP7f; R77F05-LexA; +; |
| 1-S1 | f, g | DNp15 | hsFLP; R11E07-p65.AD / +; R77F05-Gal4.DBD /<br>10XUAS>Stop>eGFPKir2.1; |
| 1,1-S1,3,3-S2,3-S3 |  | HS | w<sup>1118</sup>; R27B03-p65.AD / 20XUAS-6XmCherry;<br>R39E01-Gal4.DBD / UAS-sytGCaMP7f;<br>w<sup>1118</sup>; 20XUAS-jGCaMP7f / +; VT058487-Gal4 / UAS-CD4::tdTomato<br>w<sup>1118</sup>; R39E01-LexA, 13XLexAop2-6XmCherry / +;<br>VT058487-Gal4 / UAS-sytGCaMP7f |
| 3,3-S2,3-S3 |  | H2 | w<sup>1118</sup>; R23C12-p65.AD / 20XUAS-6XmCherry;<br>R32A11-Gal4.DBD / UAS-sytGCaMP7f;<br>w<sup>1118</sup>; R23C12-p65.AD / R39E01-LexA, 13XLexAop2-6XmCherry; R32A11-Gal4.DBD / UAS-sytGCaMP7f |
| 3,3-S2,3-S3 |  | bIPS | w<sup>1118</sup>; R25F07-p65.AD / 20XUAS-6XmCherry;<br>VT014724-Gal4.DBD / UAS-sytGCaMP7f;<br>w<sup>1118</sup>; R25F07-p65.AD / 20XUAS-jGCaMP7f;<br>VT014724-Gal4.DBD / UAS-CD4::tdTomato |
| 3,3-S2,3-S3 |  | H2rn | w<sup>1118</sup>; VT040346-p65.AD / 20XUAS-6XmCherry;<br>VT049923-Gal4.DBD / UAS-sytGCaMP7f; |
| 3,3-S2,3-S3 |  | uLPTCrn | w<sup>1118</sup>; 20XUAS-6XmCherry / +; R50G03-Gal4 / UAS-sytGCaMP7f;<br>w<sup>1118</sup>; R39E01-LexA, 13XLexAop-6XmCherry / +;<br>R50G03-Gal4 / UAS-sytGCaMP7f |
| 3-S1 | a | bIPS | w<sup>1118</sup>; R25F07-p65.AD / UAS-mCD8::GFP;<br>VT014724-Gal4.DBD / +; |
| 3-S1 | a | H2rn | w<sup>1118</sup>; VT040346-p65.AD / UAS-mCD8::GFP;<br>VT049923-Gal4.DBD / +; |
| 3-S1 | a | uLPTCrn | w<sup>1118</sup>; Otd-nls:FLPo / +; R50G03-Gal4 / pJFRC56-10XUAS>Stop>eGFPKir2.1; |
| 3-S1 | a | LPTCrn | w<sup>1118</sup>; VT045281-p65.AD / Otd-nls:FLPo; VT038829-Gal4.DBD / pJFRC56-10XUAS>Stop>eGFPKir2.1; |
| 3-S1 | a | HS | w<sup>1118</sup>; R27B03-p65.AD / 20XUAS-jGCaMP7f;<br>R39E01-Gal4.DBD / UAS-CD4::TdTomato; |
| 3-S1 | a | H2 | w<sup>1118</sup>; R23C12-p65.AD / UAS-2xEGFP; R32A11-Gal4.DBD / +; |
| 3-S1 | a | DNp15 | w<sup>1118</sup>; R11E07-p65.AD / 20XUAS-jGCaMP7f;<br>R77F05-Gal4.DBD / 20XUAS-6XmCherry; |
| 3-S1 | b | LPTCrn | w<sup>1118</sup> / MCFO-1; VT045281-p65.AD / +; VT038829-Gal4.DBD / MCFO-1; |
| 3-S1 | c | bIPS Rdl +/- | w<sup>1118</sup>; R25F07-p65.AD / 20XUAS-GCaMP6f, UAS-FLP; VT014724-Gal4.DBD / Rdl<sup>FlpStop ND</sup>; |
| 3-S1 | c | bIPS Rdl -/- | w<sup>1118</sup>; R25F07-p65.AD / 20XUAS-GCaMP6f, UAS-FLP; VT014724-Gal4.DBD, Rdl<sup>FlpStop ND</sup> / Rdl<sup>FlpStop ND</sup>; |
| 4 |  | bIPS Rdl +/+ | w<sup>1118</sup>; R25F07-p65.AD / 20XUAS-GCaMP6f, UAS-FLP; VT014724-Gal4.DBD / 20XUAS-6XmCherry; |
| 4 |  | bIPS Rdl -/- | w<sup>1118</sup>; R25F07-p65.AD / 20XUAS-GCaMP6f, UAS-FLP; VT014724-Gal4.DBD, Rdl<sup>FlpStop ND</sup> / Rdl<sup>FlpStop ND</sup>; |
| 4-S1 | a | LPTCrn | w<sup>1118</sup>; VT045281-p65.AD / UAS-mCD8::GFP;<br>VT038829-Gal4.DBD / +; |
| 4-S1 | a | uLPTCrn | w<sup>1118</sup>; UAS-mCD8::GFP / +; R50G03-Gal4 / +; |
| 4-S1 | a | bIPS | w<sup>1118</sup>; VT014724-p65.AD / UAS-mCD8::GFP;<br>VT039047-Gal4.DBD / +; |
| 4-S1 | a | H2rn | w<sup>1118</sup>; VT040346-p65.AD / UAS-mCD8::GFP;<br>VT049923-Gal4.DBD / +; |
| 4-S1 | b | LPTCrn ∩ GAD1 | w<sup>1118</sup>; UAS-mCD8::GFP / +; VT045281-Gal4.DBD / GAD1-p65.AD; |
| 4-S1 | b | LPTCrn ∩ ChaT | w<sup>1118</sup>; VT045281-p65.AD / UAS-mCD8::GFP; ChaT-Gal4.DBD / +; |
| 4-S1 | b | LPTCrn ∩ VGlut | w<sup>1118</sup>; VT045281-p65.AD / VGlut-Gal4.DBD; 40xUAS-mCD8::GFP; |
| 4-S1 | b | uLPTCrn ∩ GAD1 | w<sup>1118</sup>; UAS-mCD8::GFP / +; GAD1-p65.AD / R50G03-Gal4.DBD; |
| 4-S1 | b | uLPTCrn ∩ ChaT | w<sup>1118</sup>; R50G03-p65.AD / UAS-mCD8::GFP; ChaT-Gal4.DBD / +; |
| 4-S1 | b | uLPTCrn ∩ VGlut | w<sup>1118</sup>; R50G03-p65.AD / VGlut-Gal4.DBD; 40xUAS-mCD8::GFP; |
| 4-S1 | b | bIPS ∩ GAD1 | w<sup>1118</sup>; UAS-mCD8::GFP / +; GAD1-p65.AD / VT014724-Gal4.DBD; |
| 4-S1 | b | bIPS ∩ ChaT | w<sup>1118</sup>; VT014724-p65.AD / UAS-mCD8::GFP; ChaT-Gal4.DBD / +; |
| 4-S1 | b | bIPS ∩ VGlut | w<sup>1118</sup>; VGlut-p65.AD / UAS-mCD8::GFP; VT019845-Gal4.DBD / +; |
| 4-S1 | b | H2rn ∩ GAD1 | w<sup>1118</sup>; UAS-mCD8::GFP / CyO; GAD1-p65.AD / VT049923-Gal4.DBD; |
| 4-S1 | b | H2rn ∩ ChaT | w<sup>1118</sup>; VT040346-p65.AD / UAS-mCD8::GFP; ChaT-Gal4.DBD / +; |
| 4-S1 | b | H2rn ∩ VGlut | w<sup>1118</sup>; VGlut-p65.AD / UAS-mCD8::GFP; VT049923-Gal4.DBD / +; |
| 6 | b | DNp15 | hsFLP; R11E07-p65.AD / +; R77F05-Gal4.DBD / 10XUAS>Stop>eGFPKir2.1; |
| 6 | c | bIPS | hsFLP; R25F07-p65.AD / +; VT014724-Gal4.DBD / 10XUAS>Stop>eGFPKir2.1; |
| 6 | d | empty control >TrpA1 (control 1) | w<sup>1118</sup>; Empty-p65.AD / UAS-TrpA1(B).K; Empty-Gal4.DBD / +; |
| 6 | d | bIPS #1 >TrpA1 | w<sup>1118</sup>; R25F07-p65.AD / UAS-TrpA1(B).K; VT014724-Gal4.DBD / +; |
| 6 | d | bIPS #2 >TrpA1 | w<sup>1118</sup>; VT014724-p65.AD / UAS-TrpA1(B).K; VT039047-Gal4.DBD / +; |
| 6 | d | bIPS #3 >TrpA1 | w<sup>1118</sup>; VT039047-p65.AD / UAS-TrpA1(B).K; VT019845-Gal4.DBD / +; |
| 6 | d | empty control > Kir2.1 (control 2) | w<sup>1118</sup>; Empty-p65.AD / Otd-nls:FLPo ; Empty-Gal4.DBD / pJFRC56-10XUAS>Stop>eGFPKir2.1; |
| 6 | d | bIPS #1 > Kir2.1 | w<sup>1118</sup>; R25F07-p65.AD / Otd-nls:FLPo; VT014724-Gal4.DBD / pJFRC56-10XUAS>Stop>eGFPKir2.1; |
| 6-S1 | a | bIPS #1 | w<sup>1118</sup>; R25F07-p65.AD / UAS-mCD8::GFP; VT014724-Gal4.DBD / +; |
| 6-S1 | a | bIPS #2 | w<sup>1118</sup>; VT014724-p65.AD / UAS-mCD8::GFP; VT039047-Gal4.DBD / +; |
| 6-S1 | a | bIPS #3 | w<sup>1118</sup>; VT039047-p65.AD / UAS-mCD8::GFP;<br>VT019845-Gal4.DBD / +; |
| 6-S1 | b | bIPS | hsFLP; R25F07-p65.AD / +; VT014724-Gal4.DBD /<br>pJFRC56-10XUAS>Stop>eGFPKir2.1; |

## STAR ★ METHODS

### KEY RESOURCES TABLE

| Reagent or resource | Source | ID |
| --- | --- | --- |
| **Antibodies** |  |  |
| mouse anti-nc82 antibody | DSHB | RRID: AB_2314866 |
| chicken anti-GFP antibody | Abcam | RRID: AB_13970 |
| Rabbit polyclonal anti-GFP antibody | Thermo Fisher Scientific | Cat#A6455<br>RRID: AB_221570 |
| rabbit anti-GABA antibody | Sigma | RRID: A2052 |
| rabbit anti-DsRed antibody | Abcam | AB356483 |
| goat Alexa-488 anti-chicken | Invitrogen | RRID: A11038 |
| goat Alexa-594 anti-rabbit | Invitrogen | RRID: AB_150088 |
| goat Alexa-633 anti-mouse | Invitrogen | RRID: AB_141431 |

| Experimental Models: Organisms/Strains |  |  |
| --- | --- | --- |
| *Drosophila* UAS-2xEGFP | Bloomington<br>Drosophila Stock<br>Center (BDSC) | RRID: BDSC_6874 |
| *Drosophila* UAS-mCD8::GFP | BDSC | RRID: BDSC_5137 |
| *Drosophila* 40XUAS-mCD8::GFP | BDSC | RRID: BDSC_32195 |
| *Drosophila* UAS-CD4::tdTomato | BDSC | RRID: BDSC_35837 |
| *Drosophila* 20XUAS-6XmCherry in<br>VK00018 | BDSC | RRID: BDSC_52267 |
| *Drosophila* 20XUAS-6XmCherry in<br>attP2 | BDSC | RRID: BDSC_52268 |
| *Drosophila* 13XLexAop2-6XmCherry | BDSC | RRID: BDSC_52272 |
| *Drosophila* MCFO-1<sup>79</sup> | BDSC | RRID: BDSC_64085 |
| *Drosophila* 13XLexAop-jGCaMP7f | BDSC | RRID: BDSC_80910 |
| *Drosophila* 20XUAS-jGCaMP7f | BDSC | RRID: BDSC_80906 |
| *Drosophila* UAS-sytGCaMP7f | Gaby Maimon<sup>80</sup> | RRID: BDSC_94619 |
| *Drosophila* 20XUAS-GCaMP6f, UAS-FLP | Molina-Obando et<br>al.<sup>81</sup> | N/A |
| *Drosophila* hs-FLP | BDSC | RRID: BDSC_23649 |
| *Drosophila* Otd-nls:FLPo | Asahina et al<sup>82</sup> | N/A |
| *Drosophila* pJFRC56-10XUAS>Stop>eGFPKir2.1 in attP2 | Gerry Rubin | N/A |
| *Drosophila* UAS-TrpA1(B).K | BDSC | RRID: BDSC_26263 |
| *Drosophila* Rdl<sup>FlpStop ND</sup> | BDSC | RRID: BDSC_67682 |
| *Drosophila* VT014724-Gal4.DBD,<br>Rdl<sup>FlpStop ND</sup> | this study | N/A |
| *Drosophila* GAD1-p65.AD | BDSC | RRID: BDSC_60322 |
| *Drosophila* VGlut-p65.AD | BDSC | RRID: BDSC_82986 |
| *Drosophila* VGlut-Gal4.DBD | Diao et al. 2015<sup>83</sup> | N/A |
| *Drosophila* ChaT-Gal4.DBD | BDSC | RRID: BDSC_60318 |
| *Drosophila* R39E01-LexA | BDSC | RRID: BDSC_52776 |
| *Drosophila* R39E01-LexA, 13XLexAop2-6XmCherry | this study | N/A |
| *Drosophila* R77F05-LexA | BDSC | RRID: BDSC_54265 |
| *Drosophila* R50G03-Gal4 | Aljoscha Nern | N/A |
| *Drosophila* VT058487-Gal4 | Vienna Drosophila<br>Resource Center<br>(VDRC) | N/A |
| *Drosophila* w<sup>1118</sup>; R11E07-p65.AD;<br>R77F05-Gal4.DBD | BDSC | RRID: BDSC_75953 |
| *Drosophila* w<sup>1118</sup>; Empty-p65.AD;<br>Empty-Gal4.DBD | BDSC | RRID: BDSC_79603 |
| *Drosophila* R23C12-p65.AD | BDSC | RRID: BDSC_70130 |
| *Drosophila* R25F07-p65.AD | BDSC | RRID: BDSC_70622 |
| *Drosophila* R27B03-p65.AD | BDSC | RRID: BDSC_70164 |
| *Drosophila* R50G03-p65.AD | Aljoscha Nern | N/A |
| *Drosophila* VT014724-p65.AD | BDSC | RRID: BDSC_72019 |
| *Drosophila* VT039047-p65.AD | BDSC | RRID: BDSC_73268 |
| *Drosophila* VT040346-p65.AD | BDSC | RRID: BDSC_74542 |
| *Drosophila* VT045281-p65.AD | BDSC | RRID: BDSC_75164 |
| *Drosophila* R32A11-Gal4.DBD | BDSC | RRID: BDSC_69228 |
| *Drosophila* R39E01-Gal4.DBD | BDSC | RRID: BDSC_70217 |
| *Drosophila* R50G03-Gal4.DBD | Aljoscha Nern | N/A |
| *Drosophila* VT014724-p65.DBD | BDSC | RRID: BDSC_72146 |
| *Drosophila* VT019845-Gal4.DBD | BDSC | RRID: BDSC_73315 |
| *Drosophila* VT038829-Gal4.DBD | BDSC | RRID: BDSC_73679 |
| *Drosophila* VT039047-Gal4.DBD | BDSC | RRID: BDSC_73724 |
| *Drosophila* VT045281-Gal4.DBD | BDSC | RRID: BDSC_72221 |
| *Drosophila* VT049923-Gal4.DBD | BDSC | RRID: BDSC_73921 |
| **Software and Algorithms** |  |  |
| MATLAB | MathWorks, Inc. | RRID:SCR_001622 |
| R | Open-source | RRID:SCR_001905 |
| natverse<sup>84</sup> (R package) | Open-source | https://github.com/natverse |
| ScanImage | Vidrio Technologies | RRID:SCR_014307 |
| Gephi | Open-source | RRID:SCR_004293 |
| Fiji | Open-source | RRID:SCR_002285 |
| CATMAID | Open-source | RRID:SCR_006278 |

### CONTACT FOR REAGENT AND RESOURCE SHARING

All data and analysis code are available upon request directed to Eugenia Chiappe).

### EXPERIMENTAL MODEL AND SUBJECT DETAILS

#### Drosophila husbandry

*Drosophila melanogaster* were reared on a standard fly medium and kept on a 12hr light/12hr dark cycle at 25 °C. All experiments were performed with 2-to-4-days old flies. Calcium imaging experiments were carried out with female and behavior experiments with male flies. The complete set of transgenic flies and their specific sources are listed in the *Key Resources Table*. Full genotypes of flies used in each experiment are described in **Supplementary Table 3**.

### METHOD DETAILS

#### Generation of recombinant lines

For the R39E01-LexA,13xLexAop-6xmCherry recombinant, transgenic lines carrying R39E01-LexA and 13xLexAop-6xmCherry were crossed according to standard procedures. Putative recombinant progeny was scored based on eye color and ectopic mCherry labeling in the ventral cuticle and verified by immunostaining.

For the VT014724-p65.DBD, Rdl^FlpStopND^ recombinant, we could not use eye color since FlpStop cassette does not contain the *mini-white* gene. We generated stable lines from single progeny and verified recombinant lines using PCR, with primers targeting the FLPStop cassette in the ND orientation as in^53^.

#### EM datasets and neuron identification

##### EM datasets

Two different EM datasets were used: A full female adult *Drosophila* brain imaged using serial section transmission EM (FAFB^40^) and another, dense reconstruction of a partial adult female brain imaged using FIBSEM (focused ion-beam scanning electron microscopy), referred to as Hemibrain^42^. In FAFB, neuron skeletons were either manually traced using CATMAID^85,86^, following the procedure described in^40^ or automatically segmented within the FlyWire^41^ environment together with chemical synapse predictions^87^. None of the EM datasets contain information about electrical synapses (i.e., gap junctions).

##### Neural reconstructions

Manual tracing of neurons for ‘identification’ and for ‘completion’ has been done in FAFB as described in^88^. Briefly, reconstructing the full arbor of a neuron results in tracing for ‘completion’, while tracing the soma and the major axonic and dendritic branches that contain microtubules results in tracing for ‘identification’. Tracing for identification saves considerable tracing time by leaving out the many small, microtubule-less branches called twigs. This strategy is often enough to uniquely identify a neural class. For many FAFB neurons, we were able to readily obtain complete reconstructions in FlyWire with small differences in their twigs. Some neurons, however, were badly segmented in earlier instances of FlyWire and had to be reconstructed manually in FAFB. For example, DNp15 neurons have a darker cytosol and very striated dendritic branching, which resulted in incomplete automatic segmentation and connectivity in FlyWire and Hemibrain datasets. Therefore, we manually traced Right DNp15 for completion and performed input profile analysis on the FAFB dataset (**Fig. 2-S3f**). HS, VS and H2 cells were all traced to completion. Strong LPTC partners were traced for identification, most of which were then traced to completion.

A recent connectome analysis of the three EM datasets found that the FAFB dataset is left-right inverted^89^. This means the Right hemisphere neurons we reconstructed in FAFB/FlyWire are actually the Left neurons in real world coordinates. Since inter-hemisphere connectivity is highly stereotyped within and across EM datasets^89^, and doesn’t affect any of the results we present here in this study, we retain the inverted naming of neurons reconstructed in FAFB. This allows for a more intuitive comparison with the Hemibrain dataset, which largely encompasses the Right side of the brain.

##### Tracing synaptic connections

In addition to their anatomy, we have tagged the locations of all incoming (postsynaptic) and outgoing (presynaptic) chemical synapses of HS, VS, H2 cells and several classes of strong LPTC partners in FAFB. As a rule, we tagged all synaptic connections between any pair of traced neurons. Therefore, even if a neuron does not have all their synapses tagged, we obtain the total number of synaptic connections between all neurons in our FAFB dataset, provided they are fully, or near-fully traced.

##### Selection of LPTC partners to reconstruct

We traced all HS, all H2 and >90% of VS inputs for identification (**Fig. 1-S2c**). For LPTC outputs, which are ~10x more numerous than inputs, we chose a pseudo-random sampling approach. In FAFB, most of the strongly connected partners make multiple axo-dendritic connections around the same presynaptic bouton. We focused on neurons that send multiple dendrites to LPTCs within a small area by visual inspection. We then prioritized tracing these ‘putative strong’ partners for identification. In addition, we checked the strongest LPTC partners in Hemibrain and FlyWire datasets and reconstructed them in FAFB if they weren’t traced for identification already. This allowed us to reconstruct all of the strongest LPTC partners with minimal manual tracing.

#### EM to Light Microscopy (LM) matching

We used natverse R libraries^84^, in particular *templatebrains* and *rcatmaid* to register FAFB neurons onto a common template (JFRC2010) that’s used in large driver line libraries curated by the Janelia FlyLight team^90–92^. We then transformed our aligned FAFB skeletons onto a 2-D ColorMIP image^93^ and ran a similarity search between our neurons and Gal4 lines within FlyLight. Gal4 lines with high similarity were shortlisted by manual inspection. The shortlisted lines were then used in combinations to generate the Split-Gal4 lines with sparse expression^94^. Stability and sparseness of each split combination was assessed by checking GFP expression of several brains under a confocal microscope.

#### Immunohistochemistry

##### Immunostainings and microscopy

2-4 days old adult *Drosophila melanogaster* brains were dissected in phosphate buffer saline (PBS). Immunostaining protocol was slightly modified from^95^ as follows: After dissection, brains were fixed with 4% paraformaldehyde (PFA) diluted in PBS for 20 minutes in room temperature (or 1 hour on ice). Before the application of the primary antibodies, brains were incubated for 15 minutes in 10% normal goat serum (or 1 hour in 5% normal goat serum) with 0.1% Triton X-100 in PBS.

For checking GFP and GCaMP expression, Chicken anti-GFP (Abcam, CAT #ab13970, 1:1000 dilution) was used with Mouse anti-nc82 (Developmental Studies Hybridoma Bank, 1:10 dilution). In flies that additionally express TdTomato or mCherry, Rabbit anti-DsRed (Abcam, CAT #ab356483, 1:500) was also used. For GABA stainings, Chicken anti-GFP was used together with Rabbit anti-GABA (1:100; Sigma-Aldrich CAT #A2052). Secondary fluorescent antibodies were AlexaFluor594 Goat anti-Rabbit (Abcam, CAT #ab150088, 1:500), AlexaFluor488 Goat anti-Chicken (Abcam, CAT #ab150173, 1:500), AlexaFluor633 Goat anti-Mouse (Life Technologies, CAT #A-21050, 1:500). In a small number of immunostainings, Rabbit anti-GFP and AlexaFluor488 Goat anti-Rabbit were used instead.

Confocal sections were acquired using a Zeiss LSM710 confocal microscope at 1 or 2 µm intervals and maximum projections of image stacks were performed in Fiji/ImageJ (https://imagej.net/software/fiji/).

#### Calcium Imaging

##### Fly preparation for immobilized calcium imaging

2-4 days old adult female flies were used in all experiments. To keep their body size relatively constant, 2-6 flies were anaesthetized with CO_2_ after eclosion and housed in vials. On the day of imaging, to minimize the potential influence of sensory reafference signals coming from the limbs, flies were cold-anaesthetized, and their legs were removed using fine forceps from the femur-tibia joint for the forelegs and coxa-femur joint for the middle and hind legs. The femur of the forelegs were kept intact since it proved difficult to reliably wax the coxa without waxing the eyes and the proboscis. Openings in all the legs were then covered with beeswax. Proboscis was gently extended using fine forceps and waxed from below to minimize the movement of the brain during imaging. The fly was mounted on a custom-made physiology holder by temporarily pinning the thorax onto a small tungsten wire (A-M Systems #716100) using UV-cured glue (Bondic). The mounting is aided by a custom-built micromanipulator and a triple camera array to position the head in the same orientation across flies. After each experiment, high resolution images of the fly head from the side and bottom were taken to measure the head pitch, yaw and roll angles as described in^50^. Once the desired head orientation is achieved, the fly head and the body were waxed onto the physiology holder, and the UV-glued pin was removed. The cuticle from the back of the head was removed with extra sharp forceps (Dumont 5SF). Fat body tissues were removed by gentle suctioning while trachea and the muscles 1 and 16 of the fly head were removed by fine forceps. In this preparation, flies have no mobility in their legs and proboscis, very limited mobility in their wings and abdomen, and high mobility in their halteres and antennae.

##### Imaging setup

Flies ready for imaging were mounted under an upright microscope (Movable Objective Microscope, Sutter) with a 40× water-immersion objective lens (CFI Apo 40XW NIR, Nikon). Calcium imaging was performed with a custom-built galvo-galvo 2-photon laser scanning system. We used a Chameleon Ultra II Ti-Sapphire femtosecond laser (Coherent) tuned to 930 nm for GCaMP and tdTomato excitation (6 mW under the objective lens) and 780 nm for mCherry excitation when available. The longer excitation wavelength is not visible to the fly. Emission was collected on GaAsP PMT detectors (Hamamatsu, H10770PA-40) through a 535/50 nm bandpass filter for the green channel and 605/70 nm for red channel (Chroma). A 128×128 pixels slice image was acquired with ScanImage at the dorsal part of the posterior slope, where the axon terminals of all neurons that were imaged overlap. To record activity in a consistently large region across different cell types, the acquisition frame rate varied between 12.2-15.0 Hz. When mCherry or tdTomato was co-expressed with GCaMP, higher resolution 3D z-stacks were taken of the imaged regions before or after each experiment. The external solution, which perfused the preparation constantly in room temperature, contained 103 mM NaCl, 3 mM KCl, 5 mM TES, 8 mM D-trehalose, 10 mM D-glucose, 26 mM NaHCO_3_, 1 mM NaH_2_PO_4_, 4 mM MgCl_2_ and 2 mM CaCl_2_ in MilliQ water (270–275 mOsm) and was bubbled with 95% O_2_ / 5% CO_2_ (pH ~7.3).

##### Visual display

The visual display consists of a 32 × 96 array of blue light-emitting diodes (LEDs, 465 nm, Bright LED Electronics) that are modeled after Generation 3 (G3) modular display^96^ with four layers of blue filter (Rosco R385) to minimize light leaking into photomultiplier tubes used in calcium imaging. For a centrally positioned fly, its visual field subtended 216° in azimuth and 72° in elevation (pixel size of 2.25°). In an initial set of experiments, we prepared flies for imaging and measured the head pitch angle as described in^50^. The display was tilted 74° relative to the horizon to match the average pitch angle of the imaged flies.

##### Visual stimuli

Starfield stimuli^97^ were generated using MATLAB (MathWorks, Natick, MA) scripts (from https://github.com/misaacson01/Motion_Maker_G4), which were slightly modified to be compatible with the G3 display. We virtually extended our display onto a 2D spherical volume and populated this volume with 500, equally sized dots (5° solid angle diameter) at random and uniformly distributed positions. We illuminated the pixels on the LED display that overlapped with the dots in this spherical volume. Instead of having fully on/off edges, we used 16 gray-scale intensity levels which allowed for edges in visual patterns to appear to move more smoothly across frames. This increases the apparent resolution of the display while minimizing motion-independent flicker responses in recorded neurons. All dots had a nominal contrast of 100%.

Each trial consists of an identical, stationary visual pattern shown to the fly, followed by clockwise and counterclockwise movements of the pattern in a particular body axis of rotation (pitch, yaw, and roll) or movements of the pattern in translational axes (thrust and sideslip) that lasts for 2 seconds, flanked by stationary periods. Each trial lasted for 13 seconds (still-motion-still-motion-still for 2-2-5-2-2 seconds) with 5 seconds of inter-trial interval where the visual display was unlit (dark). See **Supplementary Video 1** for all optic flow patterns. In speed tuning experiments, only one direction of motion was presented, as such, trials lasted for 6 seconds (still-motion-still for 2-2-2 seconds). See **Supplementary Video 2** for speed tuning stimuli. Every visual pattern was shown 3-12 times for each fly, and the order of the trials were randomized.

##### Choice of calcium indicators

We initially tested several GCaMP variants to reliably record calcium activity across all the cell types imaged in this study. We found sytGCaMP7f^80^ to give us the strongest signal-to-noise ratio across most cell types. In addition, sytGCaMP7f guarantees imaging from the axon terminals, which allowed us to confidently distinguish calcium activity coming from Right bIPS axons versus Left bIPS dendrites. Therefore, we used sytGCaMP7f in most of our recordings. DNp15 does not have axon terminals in the brain, so we used cytoplasmic GCaMP7f^98^ instead. In FLPStop experiments we used cytoplasmic GCaMP6f^99^ which was readily recombined with UAS-FLP and used in FlpStop experiments previously^81^. In these experiments, we distinguished dendritic and axonic signals in bIPS using the anatomical landmarks provided by TdTomato expression as well as the direction selectivity of the recorded ROIs.

##### Data acquisition and alignment for imaging

Before quantifying fluorescence intensities, imaging frames were motion-corrected in MATLAB by translating each frame in the x and y plane using efficient subpixel image registration^100^ to match the time-averaged template from that session. Multiple recordings from the same fly were registered to the same template if the positional shift between recordings was small. In a small number of flies the analysis was performed on raw image traces, where the movement correction resulted in abnormal shifts of imaging frames in most of the trials despite negligible brain movement. All trials where the imaged neurons move out of the imaging plane were discarded.

#### Behavior

##### Exploratory walking arena

The virtual reality setup used in the exploratory walking experiments has been previously described^39^. Briefly, single 2-4 days old male flies with their wings clipped since eclosion walked freely in a 90 mm circular arena with heated walls. We projected an array of random dots that provide prominent visual feedback and translate together with the fly using real-time tracking of the fly’s center of mass and orientation.

To label single bIPS or DNp15 cells, flies were placed under 30°C heat shock for up to 15 minutes after their wings were removed post-eclosion. The brains of these flies were dissected immediately after each experiment to confirm cell labeling.

##### Agent fly simulations

A simulated rigid-body agent moved forward or executed a saccade in alternating activity and immobility intervals (**Fig. 5-S1d**). The distribution of the length of activity bouts was similar to the real flies’ distribution^39^. The agent explored a simulated 90 mm circular arena with hot walls. Saccades were modeled as highly stereotyped rapid events in the angular velocity of the agent. The probability and amplitude/direction of a saccade depended on the distance to the arena wall. The time intervals within an activity bout without saccades were considered forward runs. Forward runs had constant translational speed (20 mm/s) except in **Fig. 5f** where the forward velocity was varied. During forward runs, non-zero slow rotations are represented by an “injected” 1/f noise (η). The size of η was scaled by the parameter ‘Noise Level’ (*w_noise_*) to match the straightness of real flies walking in dark (**Fig. 5-S1f**).

Rotations of the agent are guided by a visual controller based on measured visual flow from both eyes. In short, four channels containing arrays of a “two-quadrant” type Hassenstein-Reichardt (HR) correlator detected self-generated visual motion cues^101^. Self-generated visual stimuli were modeled first by high-pass filtering the visual pattern at each spatial position (τ = 50ms). This was followed by a half-wave rectification step. Next, the visual signal was low-pass filtered (first order, τ = 15ms), and then multiplied with an unfiltered signal from a neighboring spatial location. This was done twice in a mirror-symmetrical manner, followed by subtraction, yielding a fully opponent direction-selective output signal, which was then scaled by the parameter ‘Visual Weight’ (*w_visual_*) to match straightness of real flies experiencing visual motion cues (**Fig. 5-S1g**). The two eyes of the agent are represented by independent arrays of HR, each detecting progressive (FtoB) and regressive (BtoF) visual rotations. Combinations of progressive and regressive signals were then used to simulate the response profiles of HS and H2 cells.

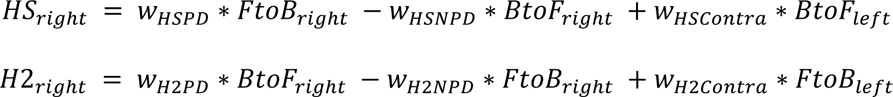

The weights for HS and H2 responses were adjusted based on a subset of horizontal visual motion responses obtained (**Fig. 5-S1a**). Following HS and H2, the intermediate layer of inhibitory neurons – bIPS (B) and uLPTCrns (U) was simulated according to the architecture outlined in **Fig. 5a**.

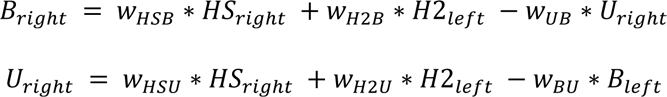

Like HS and H2, the weights for bIPS and uLPTCrn responses were also adjusted based on real visual responses (**Fig. 5-S1b**). Finally, all simulated neurons converge onto a simulated DNp15 modeled after the H2-HS network architecture in **Fig. 2**.

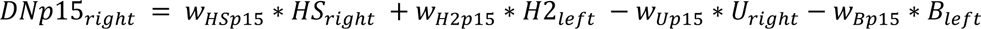

The weights for DNp15 were adjusted to match the measured horizontal visual motion responses in **Fig. 1-S1c** by iteratively changing the weights of the input layer (HS and H2) and then changing the weights of the intermediate layer (B and U). The final table of weights is presented in **Fig. 5-S1c**.

The agent’s rotation depends on the activity difference between Left and Right DNp15.

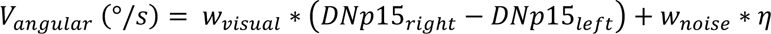

Where *w_visual_*, *w_noise_* and η correspond to the visual weight, noise level and injected 1/f noise, respectively. In the current framework, neuronal silencing was mimicked by turning the signal from specific neurons to zero, while activation was mimicked by turning the signal from specific neurons to 0.1. Asymmetry indices (AI) were calculated by comparing the activity between left and right neurons for each neural population:

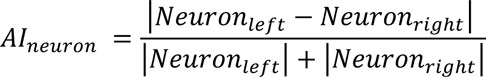

#### Data analysis

##### EM connectivity matrices and synaptic clustering

We used *natverse*^84^ R libraries to load HS, VS and H2 cells from FAFB and prune their dendritic arbor in the optic lobes, leaving only the central brain projections. We then loaded the list of all their synaptic partners, and together with HS, VS and H2 cells, we generated an all-by-all connectivity matrix where each point corresponds to the number of synapses from a row neuron to a column neuron.

To cluster the neurons that are more closely connected with each other, we first summed input and output synapses between each pair of neurons, thereby generating a symmetric connectivity table. We then binarized this table, setting all non-zero connections to 1. This prevents very strong pairwise connections to take over and distort network clustering. Conversely, it increases the weight of the weakest connections (i.e., <5 synapses). Therefore, we convert all weak connections to zero before binarization. We compute the distance matrix using this binarized connectivity and perform hierarchical clustering with Ward’s method to generate the connectivity dendrogram in **Fig. 2-S1**. The dendrogram was used to reorder the initial connectivity matrix where neurons that are clustered together are placed next to each other. We also ran the same clustering analysis without binarization nor removing weak partners (**Fig. 2-S2**) to confirm the broad network structure does not critically depend on these subjective parameters.

##### Network connectivity graphs

After generating the clusters, neurons within each cluster are loaded in CATMAID^86^. Neurons that belong to the same class (largely based on anatomy) are grouped together, generating a graph structure where each node is a neuron class and directed edges between nodes represent the number of synaptic connections. The adjacency matrix of this graph structure is then imported to Gephi where the edge and node parameters (color, size etc.) are adjusted for better visualization. The end point shape of each edge was determined by their predicted primary neurotransmitter^52^.

To measure the prominence of a neural class within each subnetwork, we plotted the weighted degree, which is the total number of incoming and outgoing synapses for each node inside that subnetwork (**Fig. 2b**).

##### Anatomical clustering of LPTC inputs

To quantify the anatomical similarities among LPTC input cell types (**Fig. 2-S3d**), we performed NBLAST^102^ on the main backbone (primary neurite tracts with the primary dendrites) of LPTC input neurons. The smallest distal dendrites (twigs) were removed since they are highly variable and make NBLAST less sensitive to the stereotyped features of cells. Hierarchical clustering was then performed on Euclidean distance matrices of NBLAST scores using Ward’s method.

##### Synaptic input profiles

To plot the neuropil input profile of LPTC input neurons (**Fig. 2-S3e, Fig. 3c**), we first obtained the list of upstream partners from FlyWire for each cell type. We only analyzed predicted synaptic sites with a cleft score of >50 to minimize false positives. We defined the input neuropil of each upstream cell as the brain region that houses most of their postsynaptic (dendritic) sites. Upstream cells that lack a soma or have less than 200 postsynaptic sites were visually inspected. The ones that project to the neck connective were scored as VNC/Neck. The remaining fragments without a cell soma were excluded from the analysis. For the DNp15 input profile (**Fig. 2-S3f**) we fully reconstructed Right DNp15 in FAFB and manually tagged all its synapses since it was badly segmented in FlyWire. We then traced ~89% of its synaptic partners for identification and defined their primary input neuropil. To plot the neurotransmitter input profile of bIPS (**Fig. 4a**) we regrouped the list of Right bIPS input neurons based on their predicted neurotransmitter^52^.

##### Distance-to-root and inter-synaptic distance analyses on bIPS

We retrieved the spatial distribution of Right bIPS inputs from FAFB using *natverse* R libraries^84^. The location of each input synapse from Right HS, H2rn, uLPTCrn and Left H2 cells were projected onto a graph representation of bIPS neuron using *nabor* library. We defined the root point of bIPS as the soma-axon-dendrite junction, which is where all the dendritic arbors of bIPS converge before crossing the midline (**Fig. 4-S2a**). We then computed the geodesic distance between each input synapse and the root point using *igraph* library (**Fig. 4-S2b,c**). This analysis is loosely based on the electrotonic distance analysis within EPG neurons performed by Hulse et. al.^103^. Geodesic distances between H2rn inputs and the nearest H2 or HS input were calculated in a similar manner (**Fig. 4-S2d**).

##### Proportion of local circuitry motifs

The non-random locations of bIPS input synapses suggest they may be organized in distinct microcircuits as they contact bIPS. We assumed only one synaptic input from a presynaptic cell takes part in a microcircuit. We therefore defined 1.865 µm as the maximum inter-synapse distance (**Fig. 4-S2d**, black line) based on the distances between H2rns with H2. We then counted the number of times a cell type (cell X) forms a microcircuit with H2 (**Fig. 4-S2e** and **4-S2g**) or HS (**Fig. 4-S2f** and **4-S2h**) as they give input to bIPS (**Fig. 4-S2d-f**) or DNp15 (**Fig. 4-S2f-h**), normalized by the total number of occurrences per motif.

##### Calcium responses to visual motion

After each imaging session, we manually defined regions of interest (ROI) that correspond to the axon terminals or dendrites of recorded neurons. For lines that label single neurons (e.g., H2, bIPS), only one ROI was defined. For lines that label multiple neurons, multiple ROIs were first defined. After confirming the visual responses of different ROIs were not qualitatively different, we chose a single ROI with the largest signal-to-noise ratio to represent the cell type. As such, we recorded the responses of no more than two ROIs per fly, one per hemisphere.

To calculate calcium evoked responses in these ROIs, we calculated mean pixel intensity (fluorescence) for each trial (F). Baseline fluorescence (F_0_) was defined as the F value of the last second before the first visual motion presentation. Change in calcium response was measured as:

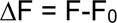

Since ΔF/F measurements of calcium activity could create positive artifacts following negative deviations from baseline observed in several cell types we imaged^104^, we instead Z normalized the ΔF data to compare optic flow responses within and across cell types. We downsampled the imaging data from each fly to the lowest recorded frame rate (12.2Hz). Recordings from left hand side cells were flipped to match the responses from right side recordings (and vice versa for H2 recordings). Matching is done by converting the stimulus identity (e.g. roll CW, pitch CW and yaw CW) into its midline-symmetric version (roll CCW, pitch CCW and yaw CCW), similar to what is described in Weir et al.^97^. Finally, all repetitions of a given stimulus were averaged to get a single mean response per stimulus per ROI.

##### Optic flow response indices

To measure the relative sensitivity of a neuron to a particular pattern of optic flow, we calculated the response as the area under the curve (auc) during the entire 2 seconds of visual motion stimulation or calculated the mean response within the last 200 ms of visual motion (peak). The preferred (PD) and non-preferred (NPD) direction of visual motion for all axes of movement were defined based on Right HS cell responses. Optic Flow (OF) responses (**Fig. 3c**) were plotted as the difference between calcium responses of the preferred (PD) and non-preferred (NPD) direction related to a given stimulus (e.g., CW minus CCW). The OF responses in **Fig. 3d** were normalized by the peak OF response per ROI.

To measure the discrimination index (DI), we first calculated the peak OF response for a given optic flow pattern (e.g., Yaw). Here, PD and NPD responses were defined per cell type:

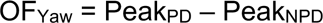

The DI between two optic flow patterns (e.g., Progressive and Yaw) is calculated as the difference between OF values divided by the sum of the absolute OF values:

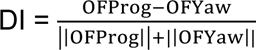

##### Classification of walking paths, walking straightness, and angular deviations

Walking paths are classified into spike-like saccadic events that correspond to rapid turns and forward segments as described in Cruz et al.^39^. Straightness of forward segments are calculated by first measuring local deviations from an ideal path within a running window of 333 ms for each point of the fly trajectory. Straightness is defined as the total distance traveled within each window divided by the sum of local deviations. Angular path deviation for each forward segment is defined as the cumulative angular deviation of the body divided by the total distance traveled. Angular bias is the binarized version of angular path deviation (**Fig. 1g, 5-S1e**).

### QUANTIFICATION AND STATISTICAL ANALYSIS

#### Statistics

We performed a two-sided Wilcoxon rank-sum (here, abbreviated as MWW) test for comparisons between two independent groups with Bonferroni correction for multiple comparisons where applicable.

### DATA AND CODE AVAILABILITY

Analysis codes and data in this paper are available upon request to the Lead Contact.

## Supplementary figure and video legends

**Figure 1-S1: DNp15’s responses to optic flow and their effect on body saccades** (**a**) Mean responses to rightward horizontal binocular optic flow moving in different speeds for Right HS (magenta) and right DNp15 (olive). The speed of the stimuli presented in the other panels of this figure, and of the rest of the figures in this study is highlighted in red. Gray shaded area indicates the duration of visual stimulation. Each trace corresponds to the grand mean of z-scored calcium responses. The mean response to 5-10 repetitions per stimulus from a single region of interest (ROI) at the axon terminals per fly was used to compute the garnd mean. Shaded area represents standard error of the mean. (**b**) Speed tuning curves calculated as the area under the curve during stimulus motion, normalized by the maximum response per ROI. (**c**) Responses of HS and DNp15 cells to different binocular and monocular optic flow stimuli. FTB= front-to-back, BTF= back-to-front visual motion (**d**) Top, schematic illustrating directions of body translations and the resulting optic flow without rotating the head. Bottom, right HS (magenta) and DNp15 (olive) responses to translational optic flow. (**e**) Discrimination index for HS and DNp15 between different patterns of optic flow. Each circle represents an individual ROI. Colored horizontal line and black vertical line represents mean and 95% CI. (HS: N=12 flies, n=16 ROIs, DNp15: N=6 flies n=7 ROIs, *P < 0.05; **P < 0.01 ***P < 0.001, Wilcoxon’s rank-sum test). (**f**) Immunostainings showing the stochastic expression of kir2.1:GFP in both, none, left or right DNp15 cells. (**g**) Effect of the stochastic silencing on saccade bias (see **Methods**). Number of flies per condition, none n=25; both n=31; left n=22; right n=22. The size of circles indicates the number saccades.

**Figure 1-S2. LPTC axon terminals send outputs to and receive inputs from central neurons** (**a**) Left, schematic of the data coverage of the fly brain (dark gray) available from two large-scale fly brains imaged under EM resolution, FAFB and Flywire (top) and Hemibrain (bottom). Right, reconstructions of the right HSE cell in the three EM datasets. FAFB contains skeletonized neurons while FlyWire and Hemibrain provide volumetric reconstructions of cells. Postsynaptic (input) and presynaptic (output) site locations are marked with cyan and red dots, respectively. (**b**) Example EM section at the axon terminals of HSE (yellow) and a recurrent synaptic partner (green). Arrowheads indicate the location of presynaptic active zones. Scale bar, 1 μm. (**c**) Top, number of inputs (i.e., upstream partners) for H2, HS and VS cell classes in the posterior slope region of the fly central brain. Numbers in parentheses indicate number of cells present in each cell class. Presynaptic neurons traced for identification are colored based on the EM dataset. Synapses from untraced neurons are colored in gray. Bottom, same as Top, but for outputs (i.e., downstream partners). (**d**) Top, distribution of the number of synaptic connections between LPTCs and their inputs. Color code indicates EM dataset and tracing status. Each distribution is normalized by total number of synapses. Inset shows a zoomed version of the region outlined by the dashed rectangle. Neurons with more than 10 synapses (black line) are considered strong inputs. Bottom, same as top, but for LPTC outputs. Neurons with more than 35 synapses (black line) are considered strong outputs.

**Figure 1-S3: LPTCs have few strong, and many weak synaptic partners.** (**a**) Number of inputs H2 (top), HS (middle), and VS (bottom) cells receive from central neurons. Presynaptic central neurons are ordered from strongest to weakest. Color code indicates EM data-set. Inset, input numbers normalized by the strongest input cell (**b**) same as, **a**), but for LPTC outputs. (**c**) Pairwise matching of the strongest H2, HS (n=3) and VS (n=8) outputs across EM datasets. Cells are ordered based on total number of inputs they receive from each LPTC class. Matching is primarily based on a cell’s anatomy, aided by its synaptic connectivity. Black lines represent matches where both pairs are within the top 10 strongest outputs in their respective EM dataset. Gray lines indicate that only one of the pairs is within the top 10. Dashed lines indicate uncertain matching due to the incomplete or bad segmentation in Hemibrain. See **Supplementary Table 1** for the cell identities.

**Figure 2-S1: The synaptic connectivity matrix among LPTC and their partners is structured.** The synaptic connectivity matrix of HS, VS, H2 cells and their strongest synaptic partners in the central brain. Neurons are colored based on anatomical similarities, color code as in Fig. 2 (see **Supplementary Table 2** for the complete list of neurons and nomenclature). The number of synaptic connections between pairs is indicated by the grayscale color code. In addition, connections are either colored red or blue based on the hierarchical clustering analysis, which represent whether connections belong to the H2-HS or VS networks, respectively (dendrogram, see **Methods**).

**Figure 2-S2: The structure of the connectivity matrix is largely unaffected by synaptic thresholds.** The pairwise synaptic connectivity matrix among HS, VS, H2 cells their synaptic partners in the central brain, without applying any thresholds to the connectivity strength (232 neurons total). Like in **Figure 2-S1**, number of synaptic connections between pairs is indicated by the grayscale color code. The dendrogram shows the hierarchical clustering analysis of the connectivity matrix (see **Methods**). Colored lines in the dendrogram indicate the cluster which the neuron belongs to in **Figure 2-S1**. The location of HS, H2 and VS cells within the matrix is highlighted.

**Figure 2-S3: The middle layers of the H2-HS and VS networks are characterized by distinct classes of interneurons.** (**a**) Percentage of total number of inputs to the left H2 (left), right HS (middle) and right VS cells (right) grouped by class (color code). LPTC inputs that do not belong to the five major interneuron classes are grouped together as “other”. (**b**) Posterior view of the morphology of single cell examples from each LPTC input interneuron class. Red and cyan dots mark the location of pre- and postsynaptic sites, respectively. Regions of innervation are outlined in gray. (**c**) Anatomical reconstructions of bilateral populations of neurons per class. Top, posterior views; bottom, dorsal views. Each neural class forms a bilateral, mirror symmetric population (**d**) NBLAST-based anatomical clustering of the main backbones of the LPTC input interneurons. (**e**) Left, schematic of the fly brain with key neuropils sending projections to LPTC inputs. Right, percentage of total synaptic inputs of bLPTCrn (right) and cLPTCrn (right) cells in FlyWire grouped by the main brain regions where they receive dendritic input. (**f**) Same as **e**) but for the right DNp15 from FAFB. Input neurons from H2-HS network shown in Figure 3a are highlighted with colored lines. IPS - inferior posterior slope; SPS - superior posterior slope; GNG - gnathal ganglion; LOP - lobula plate; ME - medulla; VNC - ventral nerve cord.

**Figure 3-S1: Transgenic lines used to gain access to H2-HS network neurons. (a)** Maximum intensity projections of transgenic lines labeling H2-HS network neurons. The specific cell class is indicated above each image (see **Methods** for a list of the transgenic lines). Neuropil (nc82) staining is shown in magenta, membrane targeted GFP or cytoplasmic GCaMP expression is shown in green. (**b**) Single-cell labeling using Multicolor FlpOut (MCFO) shows that the LPTCrn-Split Gal4 line contains both cLPTCrn (top) and uLPTCrn (bottom) cells. (**c**) Maximum intensity projections of transgenic lines that are heterozygous (left) or homozygous (right) mutants for *Rdl* in bIPS neurons. Neuropil staining is shown in magenta, cytoplasmic GCaMP expression is shown in green, TdTomato expression, resulting from a successfully disruption of the Rdl gene with the FLP-Stop cassette, is shown in red. Note that one copy of Rdl-FlpStop gene is enough to express TdTomato in bIPS. Scale bar for all panels: 100 μm.

**Figure 3-S2: Responses of H2-HS network neurons to different optic flow patterns.** (**a**) Top, schematics illustrating rotations of the fly’s body along six degrees of freedom. Bottom, calcium responses to the corresponding visual consequences of the body rotations from the fly’s perspective in right HS (magenta), left H2 (blue), right bIPS (green), right H2rn (red) and right uLPTCrn (orange) cells. Gray shaded area shows the window of stimulation. Each trace corresponds to the grand mean± SEM (shade) of z-scored calcium responses obtained across flies. In each fly, mean responses were obtained from 5-10 repetitions of per stimulus from a single ROI at the axon terminals of the cells. (**b**) Left, responses of right HS, left H2, right bIPS, right H2rn and right uLPTCrn to horizontal optic flow moving CW at the indicated speeds. The speed of the rotational and translational optic flow presented in other figures is highlighted in red. Calcium responses are plotted the same way as in **a**). Right, speed tuning curves for each cell type. (**c**) Same as **a**), but for monocular stimuli. FTB= front-to-back, BTF= back-to-front visual motion (**d**) Same as **a**), but for binocular symmetric optic flow. For all panels: HS: N=12 flies n=16 ROIs, H2: N =8 flies n=11 ROIs, bIPS: N=15 flies n=16 ROIs, H2rn: N=5 flies n=8 ROIs, uLPTCrn: N=5 flies n=5 ROIs.

**Figure 3-S3: Responses of H2-HS network neurons to translational optic flow** (**a**) Schematic illustrating possible body traveling directions (translations) and the corresponding optic flow without rotations of the head or body. (**b-f**) Calcium responses of right HS (**b**), left H2 (**c**), right bIPS (**d**), right H2rn (**e**) and right uLPTCrn (**f**) to translational optic flow patterns arranged in the same order as in **a**). Gray shaded area depicts the stimulation window. Each trace corresponds to the grand mean ± SEM (shaded area) of z-scored calcium responses obtained across flies. For each fly, mean responses to 5-10 repetitions per stimulus were obtained from a single ROI at the axon terminals. (**g**) Progressive-Sideslip discrimination index of neurons shown in panels **b-f**) (see **Methods**). Each circle represents an individual ROI. Colored horizontal line and black vertical line represents mean and 95% CI. (**h**) Same as **g**), but for Diagonal Translation vs. Sideslip discrimination index. For all panels: HS: N=10 flies n=10 ROIs, H2: N =8 flies n=9 ROIs, bIPS: N=9 flies n=9 ROIs, H2rn: N=5 flies n=8 ROIs, uLPTCrn: N=8 flies n=10 ROIs, *p < 0.05; **p < 0.01 ***p < 0.001, Wilcoxon’s rank-sum test with Bonferroni correction for multiple comparisons).

**Figure 4-S1: Central LPTC input neurons are GABAergic** (**a**) Representative confocal images showing immunoreactivity against GABA and GFP in cell bodies of each class of LPTC input neuron, as indicated on top of the images. Dashed circles outline cell bodies. Scale bar is 100 μm for full brain images and 10 μm for insets. (**b**) Reconstitution of the expression of each class of LPTCinput neurons by genetic intersection between LPTC input hemidrivers (columns) and GAD1 (top), ChaT (middle), or Vglut hemidrivers (bottom). Membrane-targeted GFP expression is shown in green, neuropil (nc82) staining is shown in magenta. Scale bar is 50 μm. Note the strong expression of each cell type under GAD1 hemidriver and weak to no expression under ChaT and VGlut hemidrivers. A complete list of the genotypes can be found in **Supplementary Table 3**.

**Figure 4-S2: The synaptic inputs of bIPS are organized non-randomly.** (**a**) Example of inputs to bIPS and the root point of the neuron. (**b**) Geodesic distance between bIPS root point and synaptic inputs from H2rn (n=192), HS (n=380), H2 (n=164), uLPTCrn (n=238), or a random subsample of the remaining inputs (n=192, see **Methods**). (**c**) Cumulative distributions from (**b**). (**d**) Distance between an HS, H2 or other input synapse to bIPS and the closest H2rn input to bIPS. The vertical black line indicates the maximum inter-synapse distance used to defined local connectivity. (**e-h**) Proportion of connectivity motifs that involve H2 (**e,g**) or HS (**f,h**) that converge onto bIPS (**e-f**) or DNp15 (**g-h**) with an additional neuron (cell X) color coded by cell type. Schematics at the top of each panel show the three cells involved. Numbered schematics show the connectivity motif plotted per graph.

**Figure 5-S1: Calibration and parameters of the visually guided agent.** (**a**) Visual responses of simulated and real HS and H2 cells for different preferred (PD), non-preferred (NPD) and contralateral (Con) weights. (**b**) Visual responses of simulated and real bIPS and uLPTCrn cells for different HS, H2 and cross-connection weights. (**c**) Final table with manually adjusted weights based on visual responses from **a**) and **b**). (**d**) Schematic of the state flow of the modeled agent, and the associated distributions for saccades (inset, top) and activity bout size (inset, bottom) estimated from the real data. (**e**) Definition of the quality parameters, straightness, path deviation and angular bias. (**f**) Simulated straightness under dark conditions, as a function of 1/f noise level. Black line represents the mean straightness observed in real flies. (**g**) Simulated straightness under light conditions with visual feedback, as a function of visual weight. Black line represents the mean straightness observed in real flies. (**h**) Simulated straightness under dark (black) and light (gold) conditions for the chosen noise and visual weights. Shaded areas represent the standard error of straightness observed in real flies under light and dark conditions (Cruz et al. 2021)^39^

**Figure 6-S1: Transgenic lines used in behavioral experiments** (**a**) Posterior view of maximal projection stacks of three different Split-Gal4 lines labeling bIPS. Neuropil (nc82) staining is shown in magenta and membrane targeted GFP expression is shown in green. Scale bar, 200 μm. (**b**) Posterior view of maximal stack projection of example brains with stochastic silencing of bIPS. Neuropil (nc82) staining is shown in magenta and membrane-targeted, Kir2.1-bound GFP expression is shown in green. Scale bar, 100 μm. For the complete list of genotypes of all transgenic lines, please see **Supplementary Table 3**.

**Supplementary video 1: Optic flow stimuli used in this study.** Patterns of optic flow used in this study. Note that the LED arena is curved such that the middle of the video screen is in front of the fly while the edges are on either side of the fly (Fig. 1c).

**Supplementary video 2: Speed tuning stimuli used in this study.** Example of binocular horizontal rotational optic flow presented at different speeds.

## References

1. Mahajan, N. R. & Mysore, S. P. Donut-like organization of inhibition underlies categorical neural responses in the midbrain. Nat. Commun. 2022 131 13, 1–17 (2022).

2. Hu, Y. et al. A Neural Basis for Categorizing Sensory Stimuli to Enhance Decision Accuracy. Curr. Biol. 30, 4896–4909.e6 (2020).

3. Koyama, M. & Pujala, A. Mutual inhibition of lateral inhibition: a network motif for an elementary computation in the brain. Curr. Opin. Neurobiol. 49, 69–74 (2018).

4. Jovanic, T. et al. Competitive Disinhibition Mediates Behavioral Choice and Sequences in Drosophila. Cell 167, 858–870 (2016).

5. Zhao, W. et al. A disinhibitory mechanism biases Drosophila innate light preference. Nat. Commun. 2019 101 10, 1–11 (2019).

6. Gibson, J. J. The Perception of the Visual World. The Philosophical Review vol. 60 (The Riverside Press, 1950).

7. Koenderink, J. J. Optic flow. Vision Res. 26, 161–179 (1986).

8. Wurtz, R. H. Optic flow: A brain region devoted to optic flow analysis? Curr. Biol. 8, 554–556 (1998).

9. Zhang, Y., Huang, R., Nörenberg, W. & Arrenberg, A. B. A robust receptive field code for optic flow detection and decomposition during self-motion. Curr. Biol. 32, 2505–2516.e8 (2022).

10. Kohn, J. R., Heath, S. L. & Behnia, R. Eyes matched to the prize: The state of matched filters in insect visual circuits. Front. Neural Circuits 12, 26 (2018).

11. Franz, M. O. & Krapp, H. G. Wide-field, motion-sensitive neurons and matched filters for optic flow fields. Biol. Cybern. 83, 185–197 (2000).

12. Barnes, W. J. P., Horseman, B. G. & Macauley, M. W. S. The Detection and Analysis of Optic Flow by Crabs: from Eye Movements to Electrophysiology. in The Crustacean Nervous System (ed. Wiese, K.) 468–485 (Springer, Berlin, Heidelberg, 2002). doi:10.1007/978-3-662-04843-6_35.

13. Britten, K. H. Mechanisms of Self-Motion Perception. https://doi.org/10.1146/annurev.neuro.29.051605.112953 31, 389–410 (2008).

14. Hausen, K. The Lobula-Complex of the Fly: Structure, Function and Significance in Visual Behaviour. in Photoreception and Vision in Invertebrates 523–559 (Springer, Boston, MA, 1984). doi:10.1007/978-1-4613-2743-1_15.

15. Matsuda, K. & Kubo, F. Circuit Organization Underlying Optic Flow Processing in Zebrafish. Front. Neural Circuits 15, 73 (2021).

16. Paulk, A. C., Phillips-Portillo, J., Dacks, A. M., Fellous, J. M. & Gronenberg, W. The processing of color, motion, and stimulus timing are anatomically segregated in the bumblebee brain. J. Neurosci. 28, 6319–6332 (2008).

17. Rasmussen, R. N., Matsumoto, A., Arvin, S. & Yonehara, K. Binocular integration of retinal motion information underlies optic flow processing by the cortex. Curr. Biol. 31, 1165–1174.e6 (2021).

18. Lappe, M. et al. Perception of self-motion from visual flow. Trends Cogn. Sci. 3, 329–336 (1999).

19. Warren, J., Kay, B. A., Zosh, W. D., Duchon, A. P. & Sahuc, S. Optic flow is used to control human walking. Nat. Neurosci. 2001 42 4, 213–216 (2001).

20. Warren, W. H., Morris, M. W. & Kalish, M. Perception of Translational Heading From Optical Flow. J. Exp. Psychol. Hum. Percept. Perform. 14, 646–660 (1988).

21. Angelaki, D. E. & Hess, B. J. M. Self-motion-induced eye movements: Effects on visual acuity and navigation. Nat. Rev. Neurosci. 6, 966–976 (2005).

22. Cullen, K. E. & Taube, J. S. Our sense of direction: Progress, controversies and challenges. Nat. Neurosci. 20, 1465–1473 (2017).

23. Theobald, J. Insect Flight: Navigating with Smooth Turns and Quick Saccades. Curr. Biol. 27, R1125–R1127 (2017).

24. Kim, A. J., Fitzgerald, J. K. & Maimon, G. Cellular evidence for efference copy in Drosophila visuomotor processing. Nat. Neurosci. 18, 1247–1255 (2015).

25. Fenk, L. M., Kim, A. J. & Maimon, G. Suppression of motion vision during course-changing, but not course-stabilizing, navigational turns. Curr. Biol. 31, 4608–4619.e3 (2021).

26. Duffy, C. J. MST neurons respond to optic how and translational movement. J. Neurophysiol. 80, 1816–1827 (1998).

27. Fujiwara, T., Cruz, T. L., Bohnslav, J. P. & Chiappe, M. E. A faithful internal representation of walking movements in the Drosophila visual system. Nat. Neurosci. 20, 72–81 (2017).

28. Borst, A., Haag, J. & Reiff, D. F. Fly motion vision. Annu. Rev. Neurosci. 33, 49–70 (2010).

29. Farrow, K., Haag, J. & Borst, A. Nonlinear, binocular interactions underlying flow field selectivity of a motion-sensitive neuron. Nat. Neurosci. 9, 1312–1320 (2006).

30. Pokusaeva, V. O., Satapathy, R., Symonova, O. & Jösch, M. Gap junctions arbitrate binocular course control in flies. bioRxiv 2023.05.31.543181 (2023) doi:10.1101/2023.05.31.543181.

31. Haag, J. & Borst, A. Orientation tuning of motion-sensitive neurons shaped by vertical-horizontal network interactions. *J. Comp. Physiol. A Neuroethol. Sensory, Neural*, Behav. Physiol. 189, 363–370 (2003).

32. Schnell, B. et al. Processing of horizontal optic flow in three visual interneurons of the Drosophila brain. J. Neurophysiol. 103, 1646–1657 (2010).

33. Cruz, T., et al. Motor context coordinates visually guided walking in Drosophila. bioRxiv 1–49 (2019) doi:10.1101/572792.

34. Zhao, A. et al. Eye structure shapes neuron function in Drosophila motion vision. bioRxiv 2022.12.14.520178 (2022).

35. Busch, C., Borst, A. & Mauss, A. S. Bi-directional Control of Walking Behavior by Horizontal Optic Flow Sensors. Curr. Biol. 28, 4037–4045.e5 (2018).

36. Fujiwara, T., Brotas, M. & Chiappe, M. E. Walking strides direct rapid and flexible recruitment of visual circuits for course control in Drosophila. Neuron 110, 2124–2138.e8 (2022).

37. Suver, M. P., Huda, A., Iwasaki, N., Safarik, S. & Dickinson, M. H. An array of descending visual interneurons encoding self-motion in Drosophila. J. Neurosci. 36, 11768–11780 (2016).

38. Namiki, S., Dickinson, M. H., Wong, A. M., Korff, W. & Card, G. M. The functional organization of descending sensory-motor pathways in drosophila. Elife 7, (2018).

39. Cruz, T. L., Pérez, S. M. & Chiappe, M. E. Fast tuning of posture control by visual feedback underlies gaze stabilization in walking Drosophila. Curr. Biol. 31, 4596–4607 (2021).

40. Zheng, Z. et al. A Complete Electron Microscopy Volume of the Brain of Adult Drosophila melanogaster. Cell 174, 730–743.e22 (2018).

41. Dorkenwald, S. et al. FlyWire: Online community for whole-brain connectomics. bioRxiv 2020.08.30.274225 (2020) doi:10.1101/2020.08.30.274225.

42. Scheffer, L. K. et al. A connectome and analysis of the adult drosophila central brain. Elife 9, 1–74 (2020).

43. Hengstenberg, R. Gaze control in the blowfly Calliphora: a multisensory, two-stage integration process. Semin. Neurosci. 3, 19–29 (1991).

44. Parsons, M. M., Krapp, H. G. & Laughlin, S. B. A motion-sensitive neurone responds to signals from the two visual systems of the blowfly, the compound eyes and ocelli. J. Exp. Biol. 209, 4464–4474 (2006).

45. Eckert, H. & Dvorak, D. R. The centrifugal horizontal cells in the lobula plate of the blowfly, Phaenicia sericata. J. Insect Physiol. 29, 547–560 (1983).

46. Wei, H., Kyung, H. Y., Kim, P. J. & Desplan, C. The diversity of lobula plate tangential cells (LPTCs) in the Drosophila motion vision system. *J. Comp. Physiol. A Neuroethol. Sensory, Neural*, Behav. Physiol. 206, 139–148 (2020).

47. Rayshubskiy, A. et al. Neural control of steering in walking Drosophila. bioRxiv 2020.04.04.024703 (2020).

48. Namiki, S. & Kanzaki, R. Brain Premotor Centers for Pheromone Orientation Behavior. 243–264 (2020) doi:10.1007/978-981-15-3082-1_12.

49. Namiki, S., Wada, S. & Kanzaki, R. Descending neurons from the lateral accessory lobe and posterior slope in the brain of the silkmoth Bombyx mori. Sci. Reports 2018 81 8, 1–19 (2018).

50. Kim, A. J., Fenk, L. M., Lyu, C. & Maimon, G. Quantitative Predictions Orchestrate Visual Signaling in Drosophila. Cell 168, 280–294.e12 (2017).

51. Fischer, P. J. & Schnell, B. Multiple mechanisms mediate the suppression of motion vision during escape maneuvers in flying Drosophila. iScience 25, 105143 (2022).

52. Eckstein, N. et al. Neurotransmitter classification from electron microscopy images at synaptic sites in Drosophila. bioRxiv (2020) doi:10.1101/2020.06.12.148775.

53. Fisher, Y. E. et al. FlpStop, a tool for conditional gene control in Drosophila. Elife 6, (2017).

54. Ffrench-Constant, R. H., Roush, R. T., Mortlock, D. & Dively, G. P. Isolation of Dieldrin Resistance from Field Populations of Drosophila melanogaster (Diptera: Drosophilidae). J. Econ. Entomol. 83, 1733–1737 (1990).

55. Ffrench-Constant, R. H., Rocheleau, T. A., Steichen, J. C. & Chalmers, A. E. A point mutation in a Drosophila GABA receptor confers insecticide resistance. Nat. 1993 3636428 363, 449–451 (1993).

56. Hosie, A. M., Aronstein, K., Sattelle, D. B. & Ffrench-Constant, R. H. Molecular biology of insect neuronal GABA receptors. Trends Neurosci. 20, 578–583 (1997).

57. Haag, J., Wertz, A. & Borst, A. Central gating of fly optomotor response. Proc. Natl. Acad. Sci. U. S. A. 107, 20104–20109 (2010).

58. Wertz, A., Haag, J. & Borst, A. Local and global motion preferences in descending neurons of the fly. J. Comp. Physiol. A. Neuroethol. Sens. Neural. Behav. Physiol. 195, 1107–1120 (2009).

59. Wertz, A., Haag, J. & Borst, A. Integration of binocular optic flow in cervical neck motor neurons of the fly. *J. Comp. Physiol. A Neuroethol. Sensory, Neural*, Behav. Physiol. 198, 655–668 (2012).

60. Strausfeld, N. J., Seyan, H. S. & Milde, J. J. The neck motor system of the fly Calliphora erythrocephala - I. Muscles and motor neurons. J. Comp. Physiol. A 160, 205–224 (1987).

61. Huston, S. J. & Krapp, H. G. Visuomotor Transformation in the Fly Gaze Stabilization System. PLOS Biol. 6, e173 (2008).

62. Haag, J., Wertz, A. & Borst, A. Integration of Lobula Plate Output Signals by DNOVS1, an Identified Premotor Descending Neuron. J. Neurosci. 27, 1992– 2000 (2007).

63. Strausfeld, N. J. & Bassemir, U. K. Lobula plate and ocellar interneurons converge onto a cluster of descending neurons leading to neck and leg motor neuropil in Calliphora erythrocephala. Cell Tissue Res. 240, 617–640 (1985).

64. Kress, D. & Egelhaaf, M. Head and body stabilization in blowflies walking on differently structured substrates. J. Exp. Biol. 215, 1523–1532 (2012).

65. Schilstra, C. & Van Hateren, J. H. Stabilizing gaze in flying blowflies. Nat. 1998 3956703 395, 654–654 (1998).

66. Seelig, J. D. & Jayaraman, V. Neural dynamics for landmark orientation and angular path integration. Nat. 2015 5217551 521, 186–191 (2015).

67. Green, J. et al. A neural circuit architecture for angular integration in Drosophila. Nat. 2017 5467656 546, 101–106 (2017).

68. Turner-Evans, D. B. et al. The Neuroanatomical Ultrastructure and Function of a Biological Ring Attractor. Neuron 108, 145–163.e10 (2020).

69. Fisher, Y. E., Marquis, M., D’Alessandro, I. & Wilson, R. I. Dopamine promotes head direction plasticity during orienting movements. Nat. 2022 6127939 612, 316–322 (2022).

70. Mauss, A. S. et al. Neural Circuit to Integrate Opposing Motions in the Visual Field. Cell 162, 351–362 (2015).

71. Braun, A., Borst, A. & Meier, M. Disynaptic inhibition shapes tuning of OFF-motion detectors in Drosophila. Curr. Biol. 33, 2260–2269.e4 (2023).

72. Kress, D. & Egelhaaf, M. Gaze characteristics of freely walking blowflies Calliphora vicina in a goal-directed task. J. Exp. Biol. 217, 3209–3220 (2014).

73. Koyama, M. et al. A circuit motif in the zebrafish hindbrain for a two alternative behavioral choice to turn left or right. Elife 5, (2016).

74. Essig, J., Hunt, J. B. & Felsen, G. Inhibitory neurons in the superior colliculus mediate selection of spatially-directed movements. Commun. Biol. 2021 41 4, 1–14 (2021).

75. Haag, J. & Borst, A. Recurrent network interactions underlying flow-field selectivity of visual interneurons. J. Neurosci. 21, 5685–5692 (2001).

76. Silies, M., Gohl, D. M. & Clandinin, T. R. Motion-Detecting Circuits in Flies: Coming into View. (2014) doi:10.1146/annurev-neuro-071013-013931.

77. Borst, A. & Helmstaedter, M. Common circuit design in fly and mammalian motion vision. Nat. Neurosci. 18, 1067–1076 (2015).

78. Agrochao, M., Tanaka, R., Salazar-Gatzimas, E. & Clark, D. A. Mechanism for analogous illusory motion perception in flies and humans. Proc. Natl. Acad. Sci. U. S. A. 117, 23044–23053 (2020).

79. Nern, A., Pfeiffer, B. D. & Rubin, G. M. Optimized tools for multicolor stochastic labeling reveal diverse stereotyped cell arrangements in the fly visual system. Proc. Natl. Acad. Sci. U. S. A. 112, E2967–E2976 (2015).

80. Lyu, C., Abbott, L. F. & Maimon, G. Building an allocentric travelling direction signal via vector computation. Nat. 2021 6017891 601, 92–97 (2021).

81. Molina-Obando, S. et al. On selectivity in the drosophila visual system is a multisynaptic process involving both glutamatergic and GABAergic inhibition. Elife 8, (2019).

82. Asahina, K. et al. Tachykinin-expressing neurons control male-specific aggressive arousal in drosophila. Cell 156, 221–235 (2014).

83. Diao, F. et al. Plug-and-Play Genetic Access to Drosophila Cell Types using Exchangeable Exon Cassettes. Cell Rep. 10, 1410–1421 (2015).

84. Bates, A. S. et al. The natverse, a versatile toolbox for combining and analysing neuroanatomical data. Elife 9, (2020).

85. Schneider-Mizell, C. M. et al. Quantitative neuroanatomy for connectomics in Drosophila. Elife 5, 1–36 (2016).

86. Saalfeld, S., Cardona, A., Hartenstein, V. & Tomančák, P. CATMAID: Collaborative annotation toolkit for massive amounts of image data. Bioinformatics 25, 1984–1986 (2009).

87. Buhmann, J. et al. Automatic detection of synaptic partners in a whole-brain Drosophila electron microscopy data set. Nat. Methods 18, 771–774 (2021).

88. Bates, A. S. et al. Complete Connectomic Reconstruction of Olfactory Projection Neurons in the Fly Brain. Curr. Biol. 30, 3183–3199.e6 (2020).

89. Schlegel, P. et al. A consensus cell type atlas from multiple connectomes reveals principles of circuit stereotypy and variation. bioRxiv 2023.06.27.546055 (2023) doi:10.1101/2023.06.27.546055.

90. Jenett, A. et al. A GAL4-Driver Line Resource for Drosophila Neurobiology. Cell Rep. 2, 991–1001 (2012).

91. Tirian, L. & Dickson, B. J. The VT GAL4, LexA, and split-GAL4 driver line collections for targeted expression in the Drosophila nervous system. bioRxiv 198648 (2017) doi:10.1101/198648.

92. Meissner, G. W. et al. An image resource of subdivided Drosophila GAL4-driver expression patterns for neuron-level searches. bioRxiv 2020.05.29.080473 (2020) doi:10.1101/2020.05.29.080473.

93. Otsuna, H., Ito, M. & Kawase, T. Color depth MIP mask search: a new tool to expedite Split-GAL4 creation. bioRxiv 318006 (2018) doi:10.1101/318006.

94. Dionne, H., Hibbard, K. L., Cavallaro, A., Kao, J. C. & Rubin, G. M. Genetic Reagents for Making Split-GAL4 Lines in Drosophila. Genetics 209, 31–35 (2018).

95. Crickmore, M. A. & Vosshall, L. B. Opposing Dopaminergic and GABAergic Neurons Control the Duration and Persistence of Copulation in Drosophila. Cell 155, 881–893 (2013).

96. Reiser, M. B. & Dickinson, M. H. A modular display system for insect behavioral neuroscience. J. Neurosci. Methods 167, 127–139 (2008).

97. Weir, P. T. & Dickinson, M. H. Functional divisions for visual processing in the central brain of flying Drosophila. Proc. Natl. Acad. Sci. U. S. A. 112, E5523–E5532 (2015).

98. Dana, H. et al. High-performance calcium sensors for imaging activity in neuronal populations and microcompartments. Nat. Methods 16, 649–657 (2019).

99. Chen, T. W. et al. Ultrasensitive fluorescent proteins for imaging neuronal activity. Nat. 2013 4997458 499, 295–300 (2013).

100. Thurman, S. T., Guizar-Sicairos, M. & Fienup, J. R. Efficient subpixel image registration algorithms. Opt. Lett. Vol. 33, Issue 2, pp. 156-158 33, 156–158 (2008).

101. Eichner, H., Joesch, M., Schnell, B., Reiff, D. F. & Borst, A. Internal Structure of the Fly Elementary Motion Detector. Neuron 70, 1155–1164 (2011).

102. Costa, M., Manton, J. D., Ostrovsky, A. D., Prohaska, S. & Jefferis, G. S. X. E. NBLAST: Rapid, Sensitive Comparison of Neuronal Structure and Construction of Neuron Family Databases. Neuron 91, 293–311 (2016).

103. Hulse, B. K. et al. A connectome of the drosophila central complex reveals network motifs suitable for flexible navigation and context-dependent action selection. Elife 10, (2021).

104. Vanwalleghem, G., Constantin, L. & Scott, E. K. Calcium Imaging and the Curse of Negativity. Front. Neural Circuits 14, 607391 (2021).

