## Supplementary Figures for "A competitive disinhibitory network for robust optic flow processing in *Drosophila*"

### Figure 1-S1: DNp15’s responses to optic flow, and their effect on body saccades

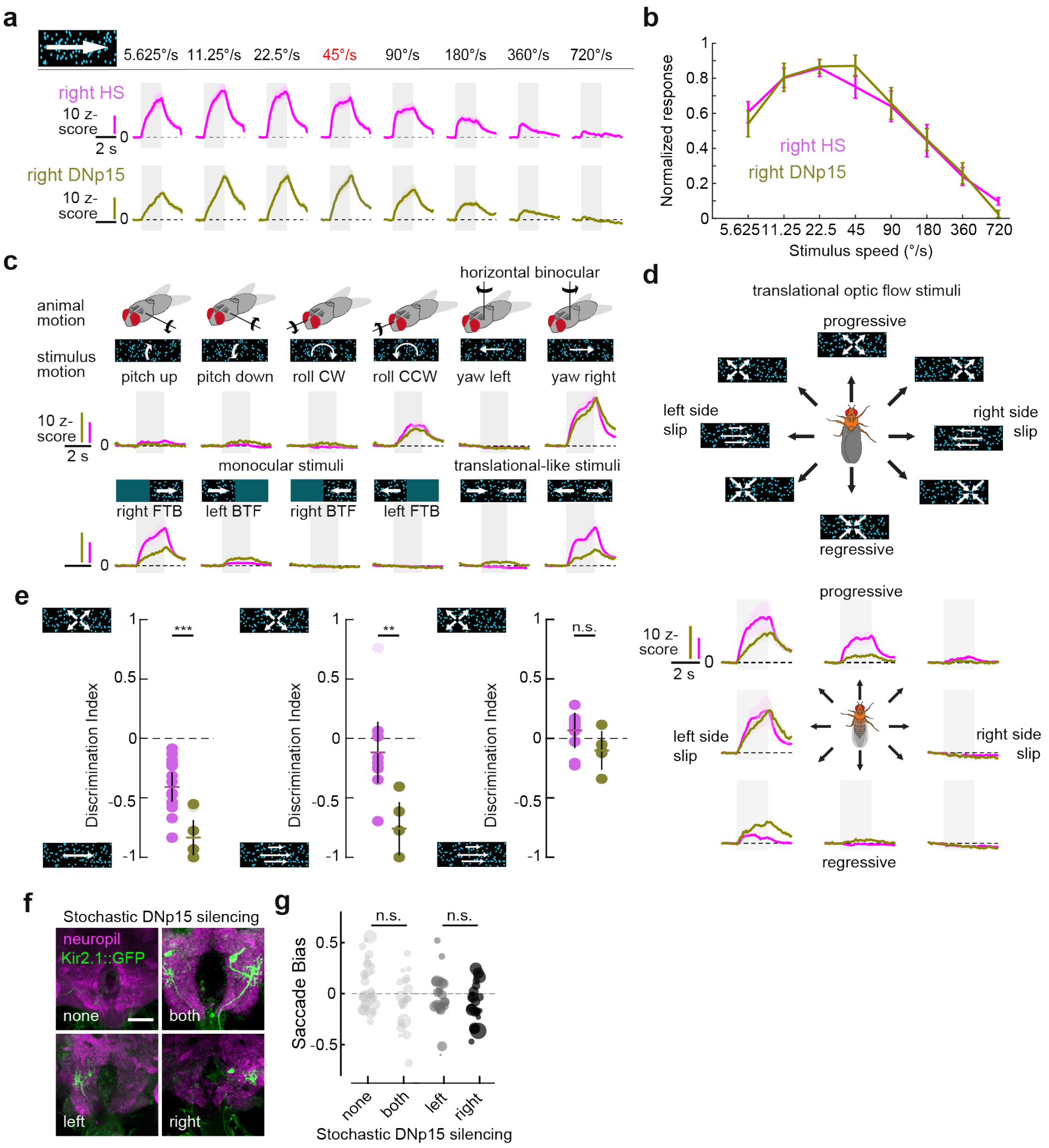


**Figure 1-S1: DNp15’s responses to optic flow and their effect on body saccades** (**a**) Mean responses to rightward horizontal binocular optic flow moving in different speeds for Right HS (magenta) and right DNp15 (olive). The speed of the stimuli presented in the other panels of this figure, and of the rest of the figures in this study is highlighted in red. Gray shaded area indicates the duration of visual stimulation. Each trace corresponds to the grand mean of z-scored calcium responses. The mean response to 5-10 repetitions per stimulus from a single region of interest (ROI) at the axon terminals per fly was used to compute the garnd mean. Shaded area represents standard error of the mean. (**b**) Speed tuning curves calculated as the area under the curve during stimulus motion, normalized by the maximum response per ROI. (**c**) Responses of HS and DNp15 cells to different binocular and monocular stimuli. FTB = front-to-back, BTF= back-to-front visual motion (**d**) Top, schematic illustrating directions of body translations and the resulting optic flow without rotating the head. Bottom, right HS (magenta) and DNp15 (olive) responses to translational optic flow. (**e**) Discrimination index for HS and DNp15 between different patterns of optic flow. Each circle represents an individual ROI. Colored horizontal line and black vertical line represents mean and 95% CI. (HS: N=12 flies, n=16 ROIs, DNp15: N=6 flies n=7 ROIs, *$P < 0.05$; **$P < 0.01$ ***$P < 0.001$, Wilcoxon’s rank-sum test). (**f**) Immunostainings showing the stochastic expression of kir2.1:GFP in both, none, left or right DNp15 cells. (**g**) Effect of the stochastic silencing on saccade bias (see **Methods**). Number of flies per condition, none n=25; both n=31; left n=22; right n=22. The size of circles indicates the number of saccades.

### Figure 1-S2. LPTC axon terminals send outputs to and receive inputs from central neurons

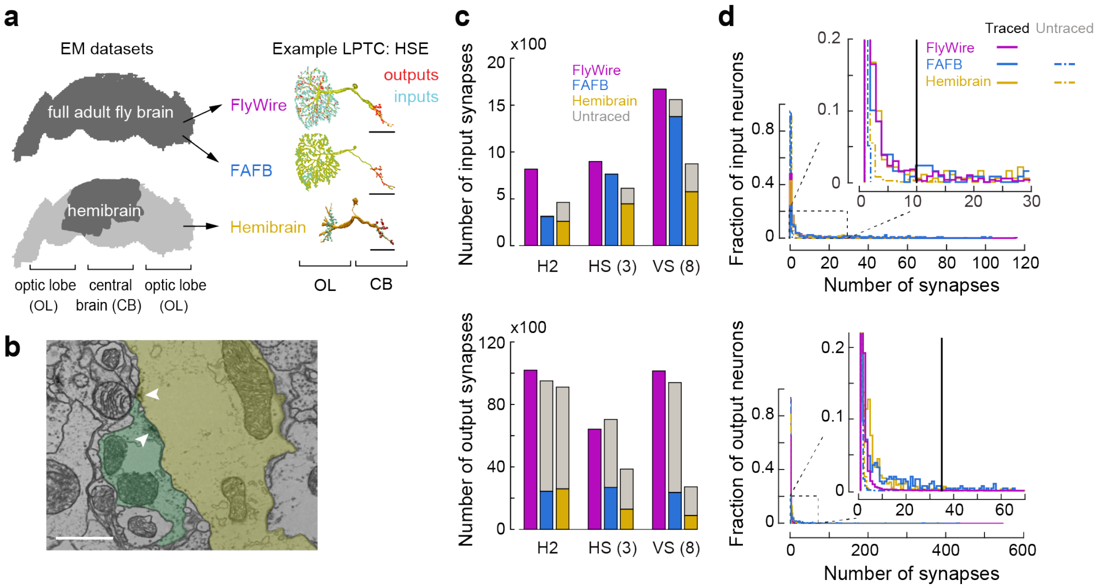


**Figure 1-S2. LPTC axon terminals send outputs to and receive inputs from central neurons** (**a**) Left, schematic of the data coverage of the fly brain (dark gray) available from two large-scale fly brains imaged under EM resolution, FAFB and Flywire (top) and Hemibrain (bottom). Right, reconstructions of the right HSE cell in the three EM datasets. FAFB contains skeletonized neurons while FlyWire and Hemibrain provide volumetric reconstructions of cells. Postsynaptic (input) and presynaptic (output) site locations are marked with cyan and red dots, respectively. (**b**) Example EM section at the axon terminals of HSE (yellow) and a recurrent synaptic partner (green). Arrowheads indicate the location of presynaptic active zones. Scale bar, 1 μm. (**c**) Top, number of inputs (i.e. upstream partners) for H2, HS and VS cell classes in the posterior slope region of the fly central brain. Numbers in parentheses indicate number of cells present in each cell class. Presynaptic neurons traced for identification are colored based on the EM dataset. Synapses from untraced neurons are colored in gray. Bottom, same as Top, but for outputs (i.e. downstream partners). (**d**) Top, distribution of the number of synaptic connections between LPTCs and their inputs. Color code indicates EM dataset and tracing status. Each distribution is normalized by total number of synapses. Inset shows a zoomed version of the region outlined by the dashed rectangle. Neurons with more than 10 synapses (black line) are considered strong inputs. Bottom, same as top, but for LPTC outputs. Neurons with more than 35 synapses (black line) are considered strong outputs.

### Figure 1-S3: LPTCs have few strong, and many weak synaptic partners

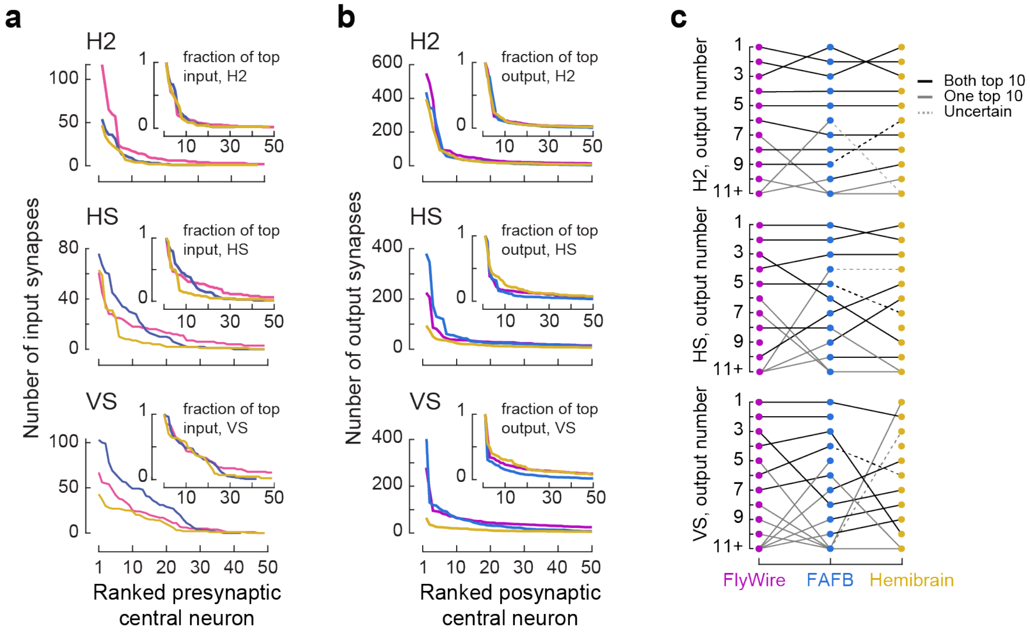


**Figure 1-S3: LPTCs have few strong, and many weak synaptic partners. (a)** Number of inputs H2 (top), HS (middle), and VS (bottom) cells receive from central neurons. Presynaptic central neurons are ordered from strongest to weakest. Color code indicates EM dataset. Inset, input numbers normalized by the strongest input cell **(b)** same as, **a**), but for LPTC outputs. **(c)** Pairwise matching of the strongest H2, HS (n=3) and VS (n=8) outputs across EM datasets. Cells are ordered based on total number of inputs they receive from each LPTC class. Matching is primarily based on a cell's anatomy, aided by its synaptic connectivity. Black lines represent matches where both pairs are within the top 10 strongest outputs in their respective EM dataset. Gray lines indicate that only one of the pairs is within the top 10. Dashed lines indicates uncertain matching due to the incomplete or bad segmentation in Hemibrain. See **Supplementary Table 1** for the cell identities.

### Figure2-S1: The synaptic connectivity matrix among LPTC and their partners is structured

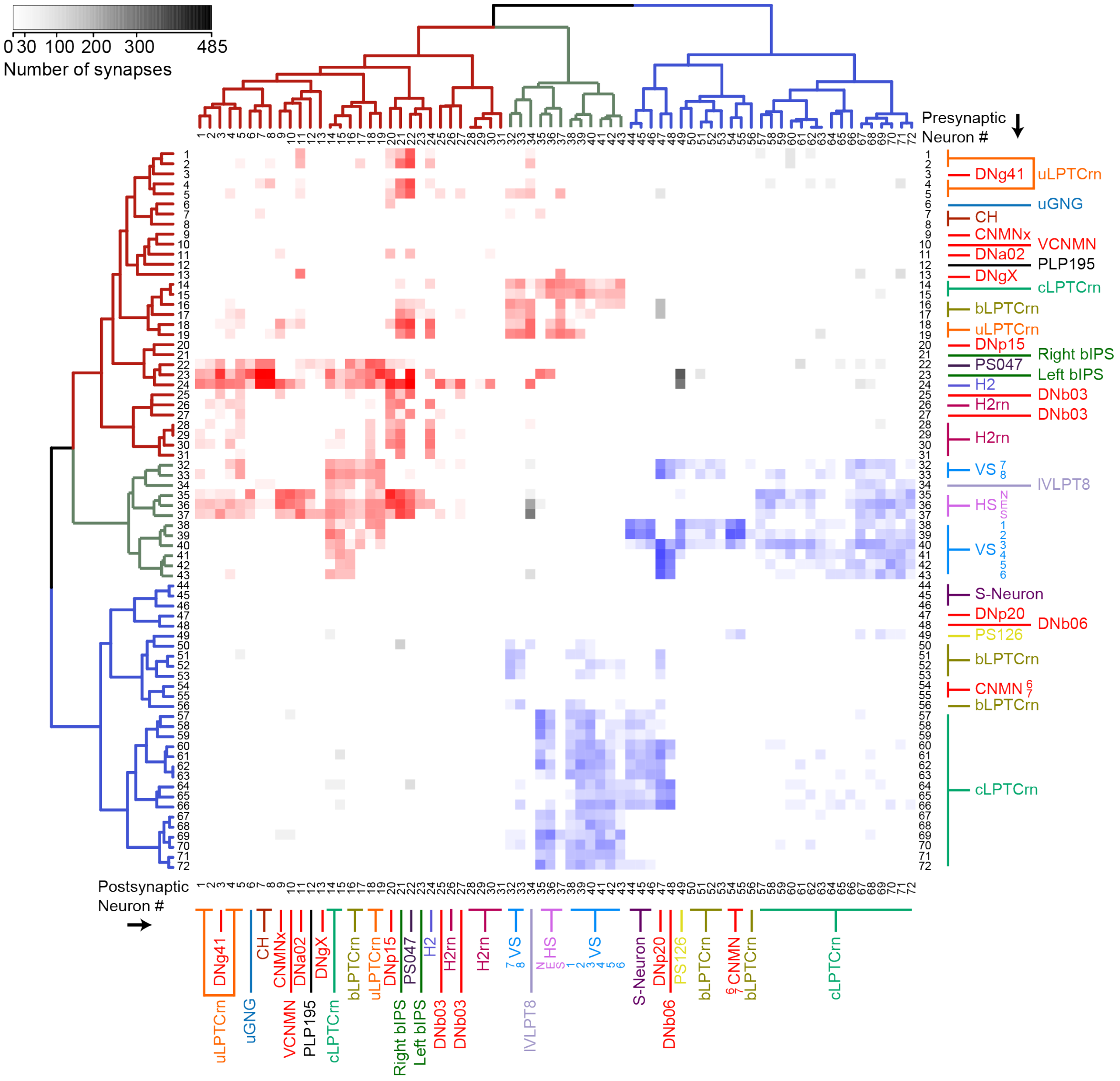


**Figure 2-S1: The synaptic connectivity matrix among LPTC and their partners is structured.** The synaptic connectivity matrix of HS, VS, H2 cells and their strongest synaptic partners in the central brain. Neurons are colored based on anatomical similarities, color code as in **Figure 2** (see **Supplementary Table 2** for the complete list of neurons and nomenclature). The number of synaptic connections between pairs is indicated by the grayscale color code. In addition, connections are either colored red or blue based on the hierarchical clustering analysis, which represent wether connections belong to the H2-HS or VS networks, respectively (dendrogram, see **Methods**).

**Figure 2-S2: The structure of the connectivity matrix is largely unaffected by synaptic thresholds**

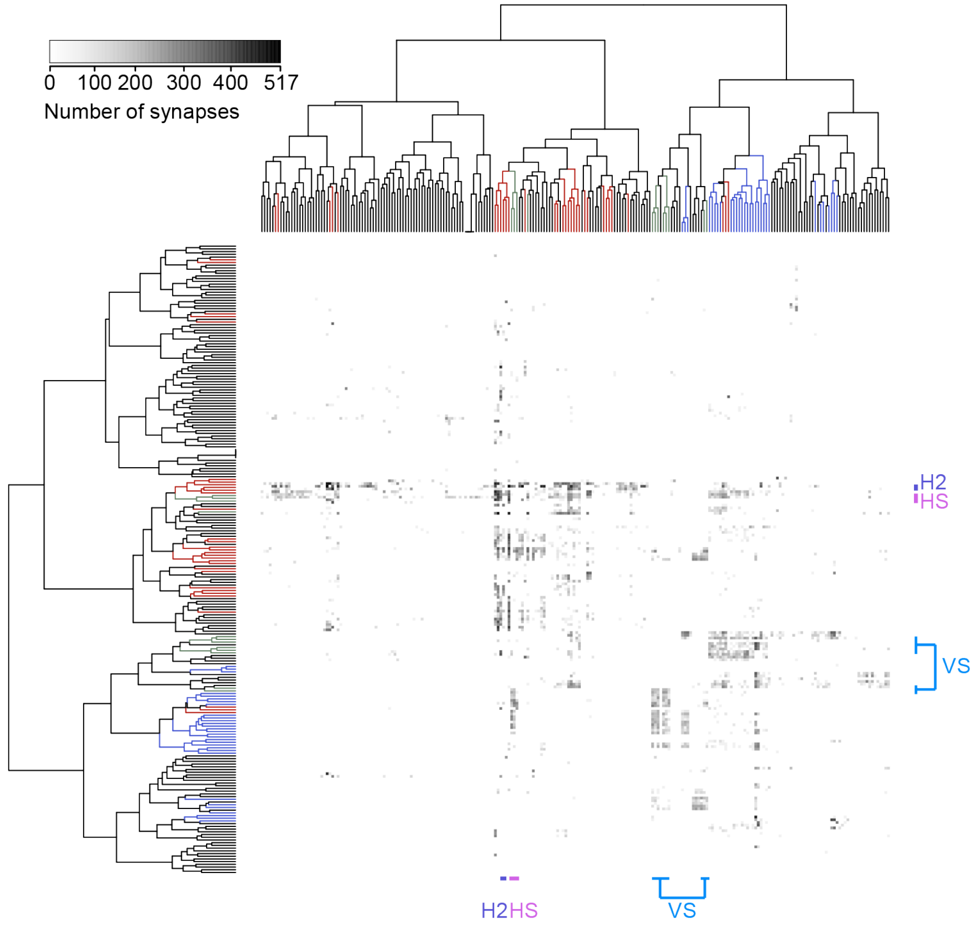


**Figure 2-S2: The structure of the connectivity matrix is largely unaffected by synaptic thresholds.** The pairwise synaptic connectivity matrix among HS, VS, H2 cells their synaptic partners in the central brain, without applying any thresholds to the connectivity strength (232 neurons total). Like in **Figure 2-S1**, number of synaptic connections between pairs is indicated by the grayscale color code. The dendrogram shows the hierarchical clustering analysis of the connectivity matrix (see **Methods**). Colored lines in the dendrogram indicate the cluster which the neuron belongs to in **Figure 2-S1**. The location of HS, H2 and VS cells within the matrix is highlighted.

### Figure2-S3: The middle layers of the H2-HS and VS networks are characterized by distinct classes of interneurons

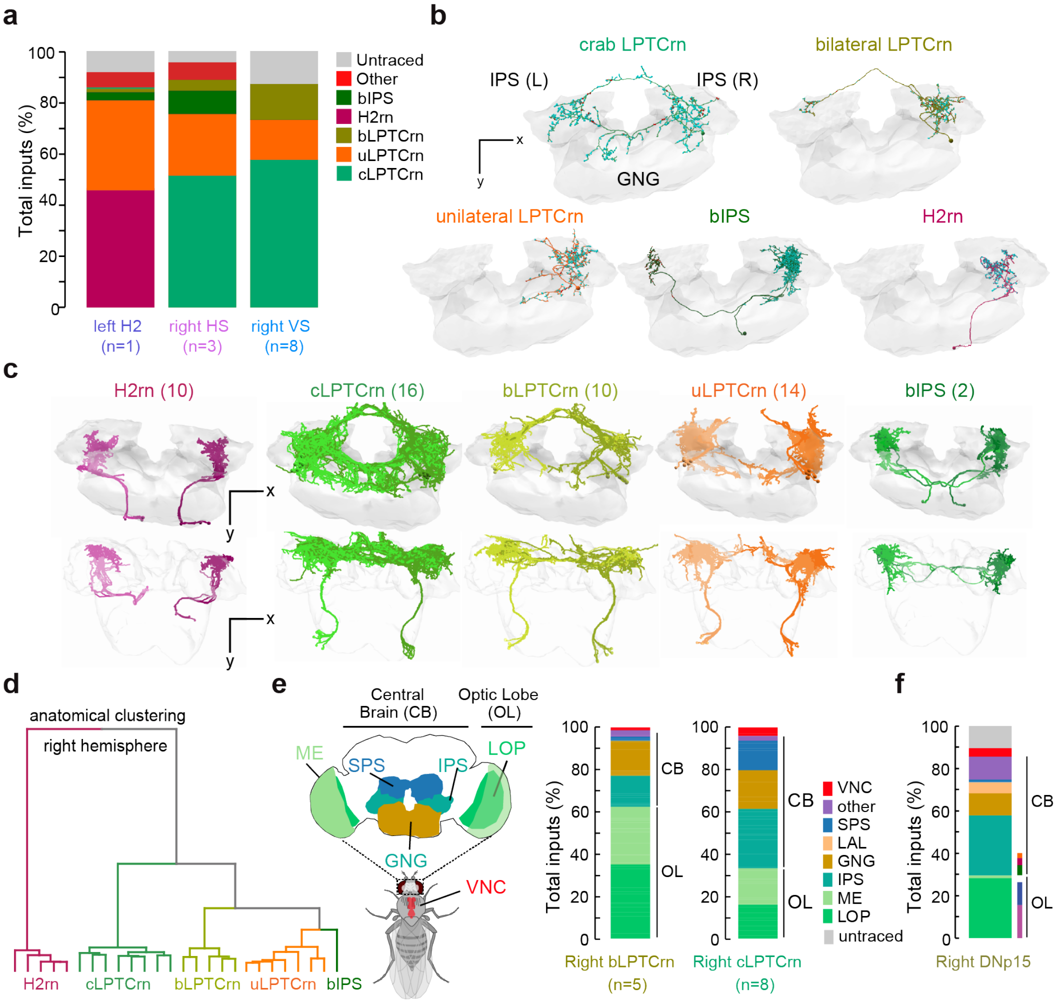


**Figure 2-S3: The middle layers of the H2-HS and VS networks are characterized by distinct classes of interneurons.** (**a**) Percentage of total number of inputs to the left H2 (left), right HS (middle) and right VS cells (right) grouped by class (color code). LPTC inputs that do not belong to the five major interneuron classes are grouped together as “other”. (**b**) Posterior view of the morphology of single cell examples from each LPTC input interneuron class. Red and cyan dots mark the location of pre- and postsynaptic sites, respectively. Regions of innervation are outlined in gray. (**c**) Anatomical reconstructions of bilateral populations of neurons per class. Top, posterior views; bottom, dorsal views. Each neural class forms a bilateral, mirror symmetric population (**d**) NBLAST-based anatomical clustering of the main backbones of the LPTC input interneurons. (**e**) Left, schematic of the fly brain with key neuropils sending projections to LPTC inputs. Right, percentage of total synaptic inputs of bLPTCrn (right) and cLPTCrn (right) cells in FlyWire grouped by the main brain regions where they receive dendritic input. (**f**) Same as **e**) but for the right DNp15 from FAFB. Input neurons from H2-HS network shown in **Figure 3a** are highlighted with colored lines. IPS - inferior posterior slope; SPS - superior posterior slope; GNG - gnathal ganglion; LOP - lobula plate; ME - medulla; VNC - ventral nerve cord.

### Figure 3-S1: Transgenic lines used to gain access to H2-HS network neurons

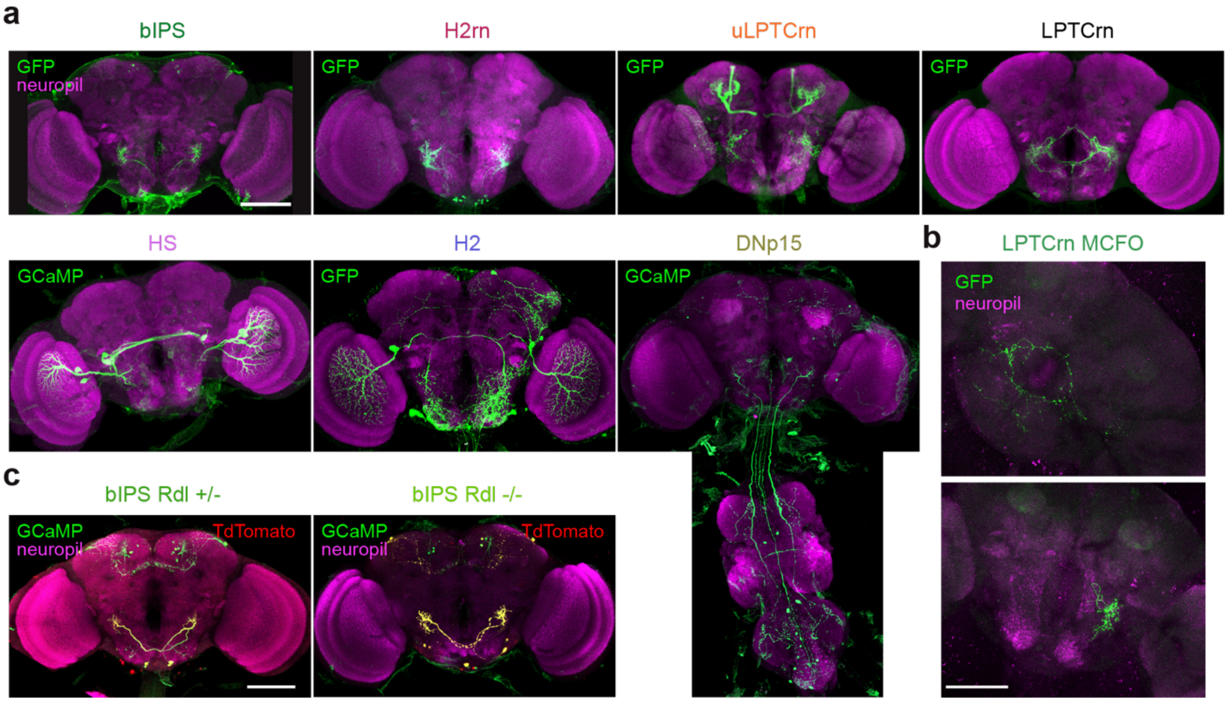


**Figure 3-S1: Transgenic lines used to gain access to H2-HS network neurons. (a)** Maximum intensity projections of transgenic lines labeling H2-HS network neurons. The specific cell class is indicated above each image (see **Methods** for a list of the transgenic lines). Neuropil (nc82) staining is shown in magenta, membrane targeted GFP or cytoplasmic GCaMP expression is shown in green. (**b**) Single-cell labeling using Multicolor FlpOut (MCFO) shows that the LPTCrn-Split Gal4 line contains both cLPTCrn (top) and uLPTCrn (bottom) cells. (**c**) Maximum intensity projections of transgenic lines that are heterozygous (left) or homozygous (right) mutants for *Rdl* in bIPS neurons. Neuropil staining is shown in magenta, cytoplasmic GCaMP expression is shown in green, TdTomato expression, resulting from a successfully disruption of the Rdl gene with the FLP-Stop cassette, is shown in red. Note that one copy of Rdl-FlpStop gene is enough to express TdTomato in bIPS. Scale bar for all panels: 100 µm.

### Figure3-S2: Responses of H2-HS network neurons to different optic flow patterns

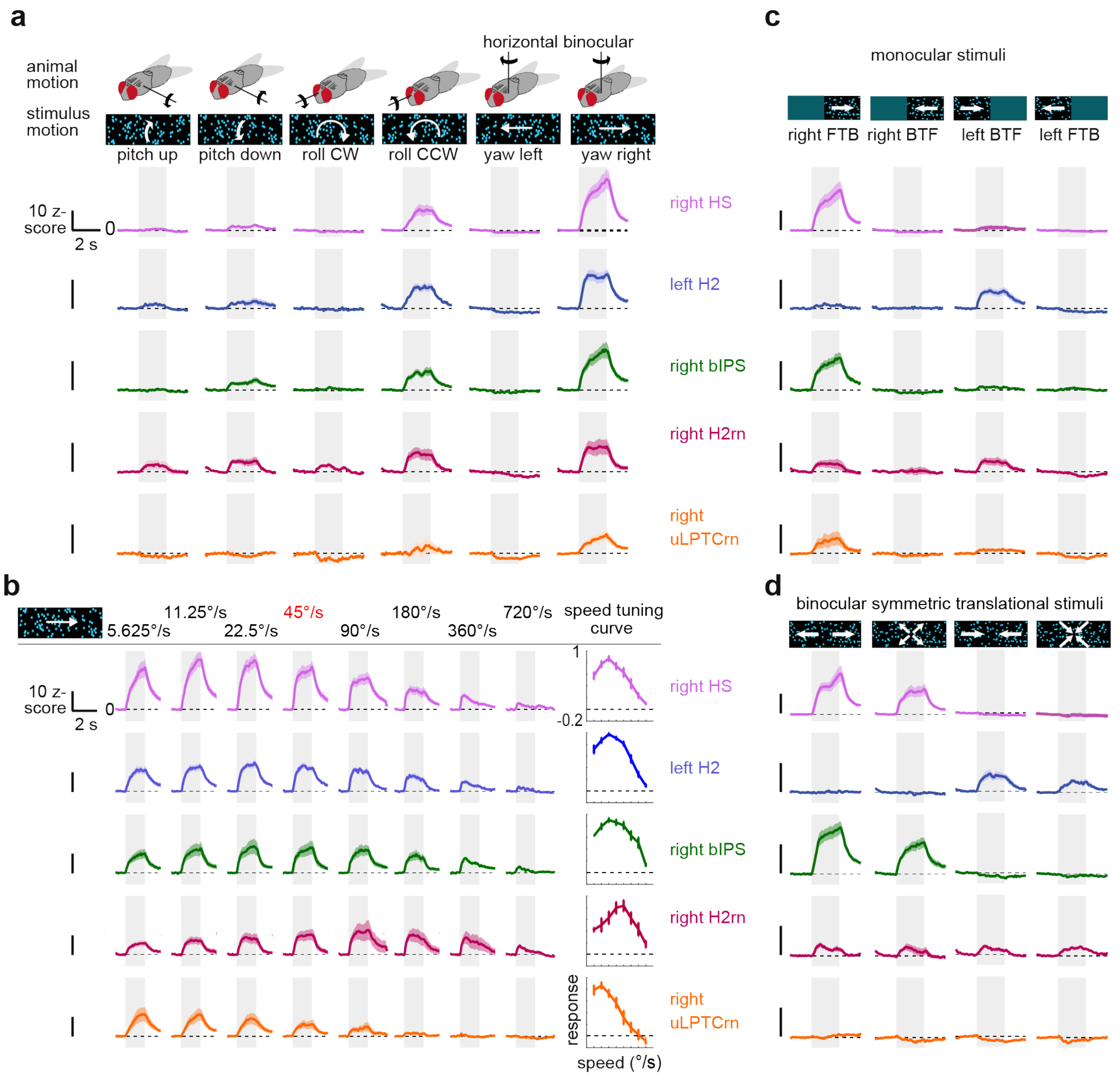


**Figure 3-S2: Responses of H2-HS network neurons to different optic flow patterns.** (**a**) Top, schematics illustrating rotations of the fly's body along six degrees of freedom. Bottom, calcium responses to the corresponding visual consequences of the body rotations from the fly's perspective in right HS (magenta), left H2 (blue), right bIPS (green), right H2rn (red) and right uLPTCrn (orange) cells. Gray shaded area shows the window of stimulation. Each trace corresponds to the grand mean± SEM (shade) of z-scored calcium responses obtained across flies. In each fly, mean responses were obtained from 5-10 repetitions of per stimulus from a single ROI at the axon terminals of the cells. (**b**) Left, responses of right HS, left H2, right bIPS, right H2rn and right uLPTCrn to horizontal optic flow moving CW at the indicated speeds. The speed of the rotational and translational optic flow presented in other figures is highlighted in red. Calcium responses are plotted the same way as in **a**). Right, speed tuning curves for each cell type. (**c**) Same as **a**), but for monocular stimuli. FTB = front-to-back, BTF= back-to-front visual motion(**d**) Same as **a**), but for binocular symmetric optic flow. For all panels: HS: N=12 flies n=16 ROIs, H2: N =8 flies n=11 ROIs, bIPS: N=15 flies n=16 ROIs, H2rn: N=5 flies n=8 ROIs, uLPTCrn: N=5 flies n=5 ROIs.

### Figure 3-S3: Responses of H2-HS network neurons to translational optic flow

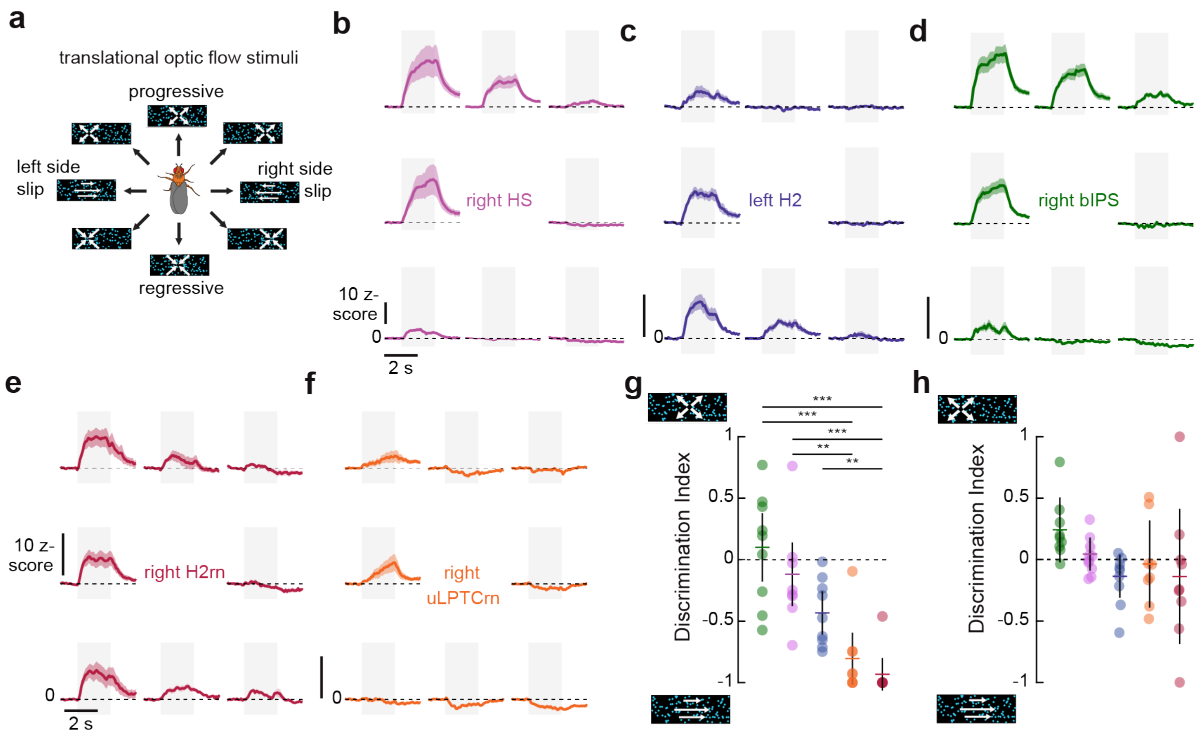


**Figure 3-S3: Responses of H2-HS network neurons to translational optic flow** (**a**) Schematic illustrating possible body traveling directions (translations) and the corresponding optic flow without rotations of the head or body. (**b-f**) Calcium responses of right HS (**b**), left H2 (**c**), right bIPS (**d**), right H2rn (**e**) and right uLPTCrn (**f**) to translational optic flow patterns arranged in the same order as in **a**). Gray shaded area depicts the stimulation window. Each trace corresponds to the grand mean ± SEM (shaded area) of z-scored calcium responses obtained across flies. For each fly, mean responses to 5-10 repetitions per stimulus were obtained from a single ROI at the axon terminals. (**g**) Progressive-Sideslip discrimination index of neurons shown in panels **b-f**) (see **Methods**). Each circle represents an individual ROI. Colored horizontal line and black vertical line represents mean and 95% CI. (**h**) Same as **g**), but for Diagonal Translation vs. Sideslip discrimination index. For all panels: HS: N=10 flies n=10 ROIs, H2: N =8 flies n=9 ROIs, bIPS: N=9 flies n=9 ROIs, H2rn: N=5 flies n=8 ROIs, uLPTCrn: N=8 flies n=10 ROIs, *$p < 0.05$; **$p < 0.01$ ***$p < 0.001$, Wilcoxon's rank-sum test with Bonferroni correction for multiple comparisons).

### Figure 4-S1: Central LPTC input neurons are GABAergic

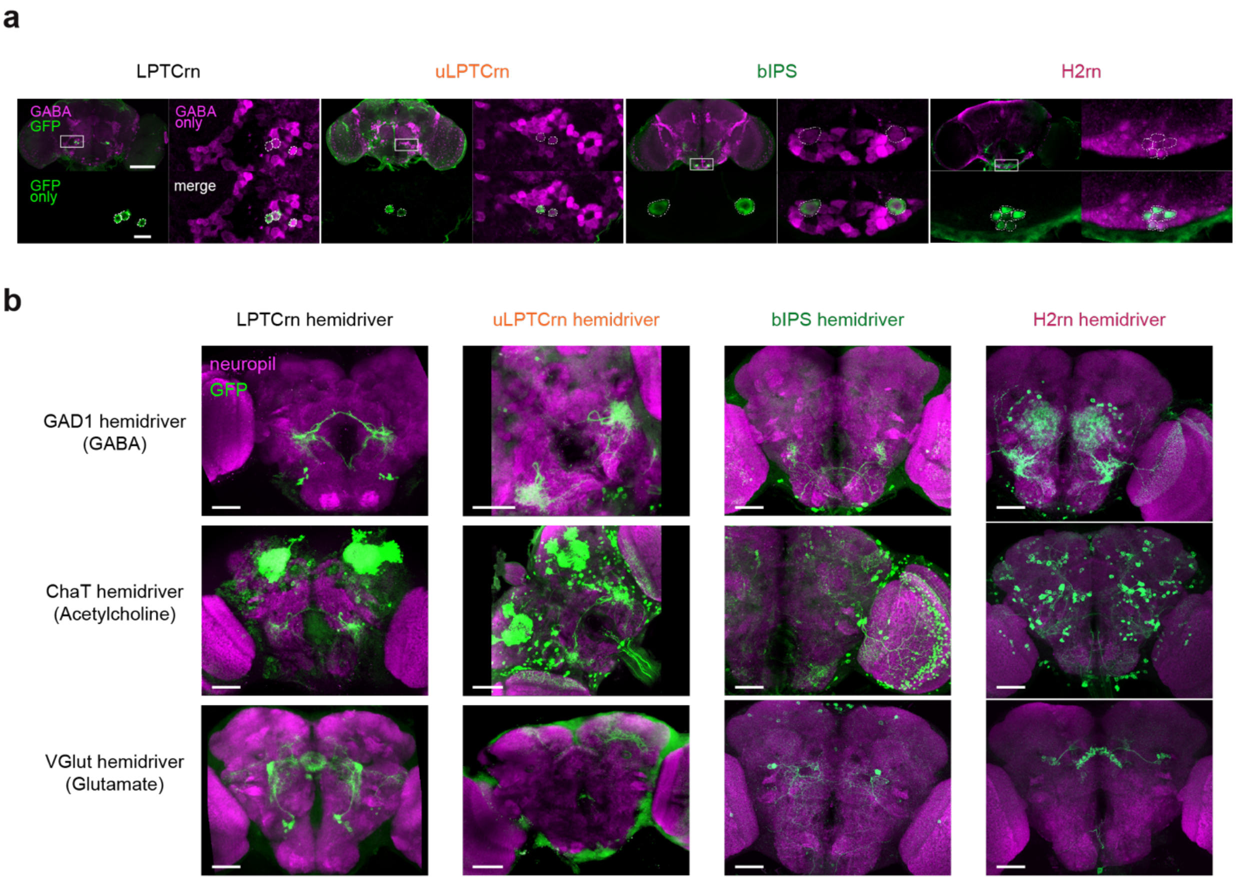


**Figure 4-S1: Central LPTC input neurons are GABAergic** (**a**) Representative confocal images showing immunoreactivity against GABA and GFP in cell bodies of each class of LPTC input neuron, as indicated on top of the images. Dashed circles outline cell bodies. Scale bar is 100 µm for full brain images and 10 µm for insets. (**b**) Reconstitution of the expression of each class of LPTC input neurons by genetic intersection between LPTC input hemidrivers (columns) and GAD1 (top), ChaT (middle), or Vglut hemidrivers (bottom). Membrane-targeted GFP expression is shown in green, neuropil (nc82) staining is shown in magenta. Scale bar is 50 µm. Note the strong expression of each cell type under GAD1 hemidriver and weak to no expression under ChaT and VGlut hemidrivers. A complete list of the genotypes can be found in **Supplementary Table 3**.

### Figure4-S2: The synaptic inputs of bIPS are organized non-randomly

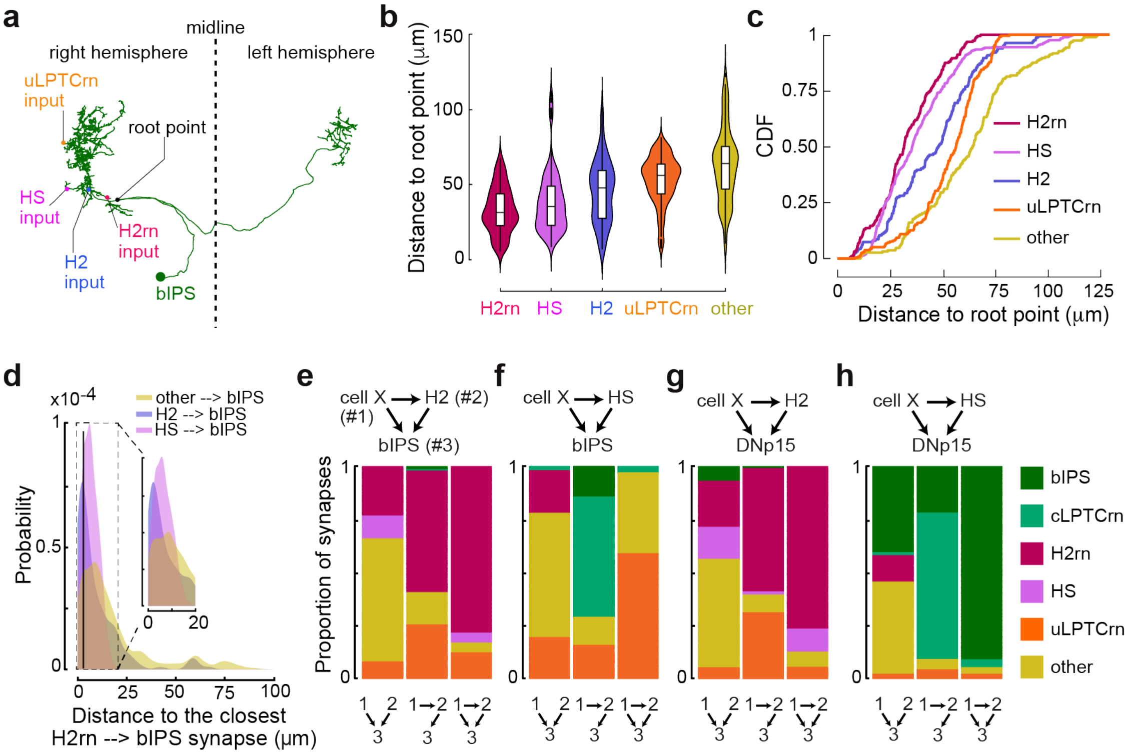


**Figure 4-S2: The synaptic inputs of bIPS are organized non-randomly.** (**a**) Example of inputs to bIPS and the root point of the neuron. (**b**) Geodesic distance between bIPS root point and synaptic inputs from H2rn (n=192), HS (n=380), H2 (n=164), uLPTCrn (n=238), or a random subsample of the remaining inputs (n=192, see **Methods**). (**c**) Cumulative distributions from (**b**). (**d**) Distance between an HS, H2 or other input synapse to bIPS and the closest H2rn input to bIPS. The vertical black line indicatesthe maximum inter-synapse distance used to define local connectivity (**e**-**h**) Proportion of connectivity motifs that involve H2 (**e**,**g**) or HS (**f**,**h**) that converge onto bIPS (**e**-**f**) or DNp15 (**g**-**h**) with an additional neuron (cell X) color coded by cell type. Schematics at the top of each panel show the three cells involved. Numbered schematics show the connectivity motif plotted per graph.

### Figure 5-S1: Calibration and parameters of the visually guided agent

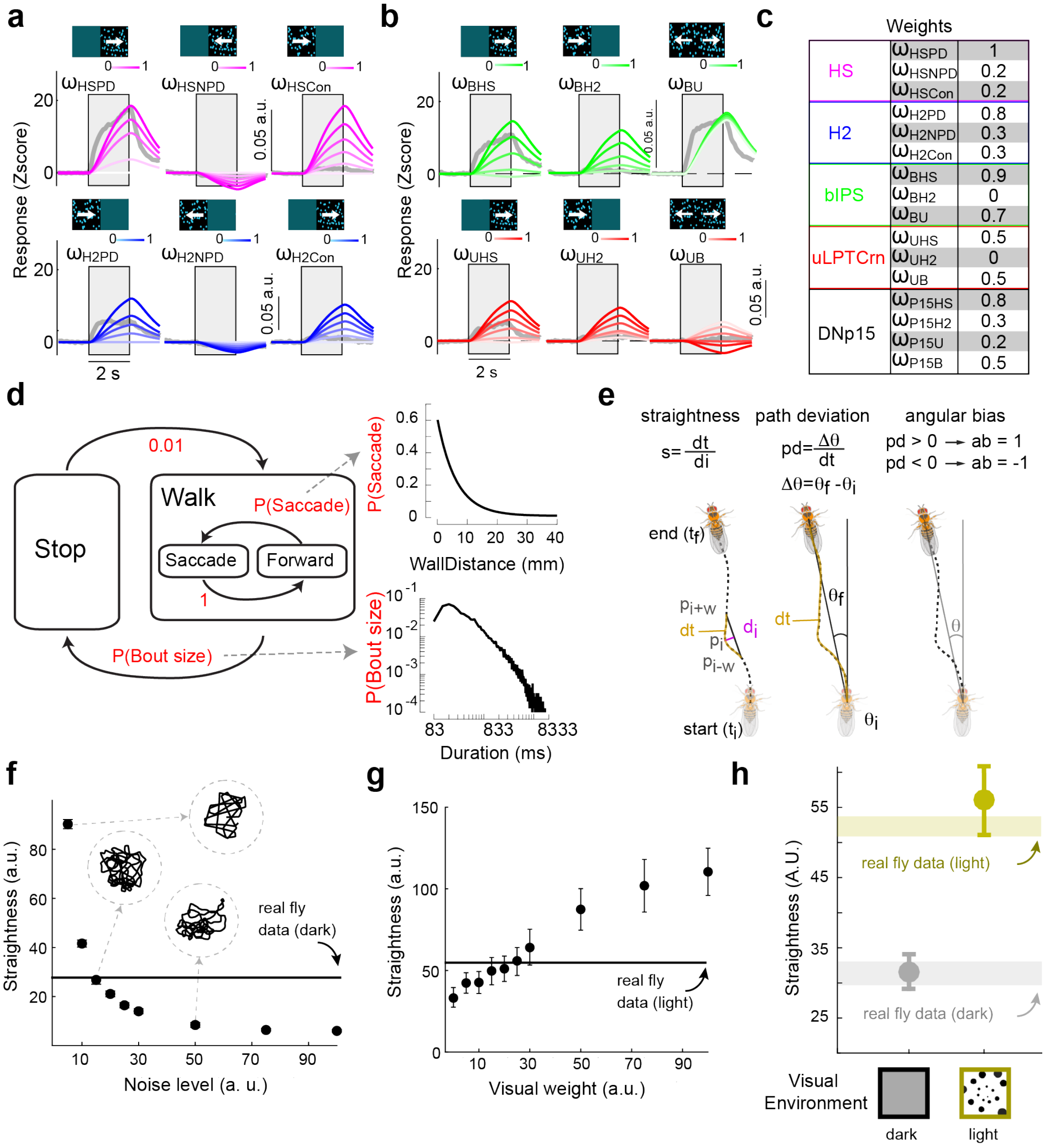


| | | |
|---|---|---|
| HS | $\omega_{HSPD}$ | 1 |
| | $\omega_{HSNPD}$ | 0.2 |
| | $\omega_{HSCon}$ | 0.2 |
| H2 | $\omega_{H2PD}$ | 0.8 |
| | $\omega_{H2NPD}$ | 0.3 |
| | $\omega_{H2Con}$ | 0.3 |
| bIPS | $\omega_{BHS}$ | 0.9 |
| | $\omega_{BH2}$ | 0 |
| | $\omega_{BU}$ | 0.7 |
| uLPTCrn | $\omega_{UHS}$ | 0.5 |
| | $\omega_{UH2}$ | 0 |
| | $\omega_{UB}$ | 0.5 |
| DNp15 | $\omega_{P15HS}$ | 0.8 |
| | $\omega_{P15H2}$ | 0.3 |
| | $\omega_{P15U}$ | 0.2 |
| | $\omega_{P15B}$ | 0.5 |

$s = \frac{dt}{di}$ $pd = \frac{\Delta\theta}{dt}$ $\Delta\theta = \theta_f - \theta_i$

$pd > 0 \rightarrow ab = 1$; $pd < 0 \rightarrow ab = -1$

**Figure 5-S1: Calibration and parameters of the visually guided agent.** (**a**) Visual responses of simulated and real HS and H2 cells for different preferred (PD), non-preferred (NPD) and contralateral (Con) weights. (**b**) Visual responses of simulated and real bIPS and uLPTCrn cells for different HS, H2 and cross-connection weights. (**c**) Final table with manually adjusted weights based on visual responses from **a**) and **b**). (**d**) Schematic of the state flow of the modeled agent, and the associated distributions for saccades (inset, top) and activity bout size (inset, bottom) estimated from the real data. (**e**) Definition of the quality parameters, straightness, path deviation and angular bias. (**f**) Simulated straightness under dark conditions, as a function of 1/f noise level. Black line represents the mean straightness observed in real flies. (**g**) Simulated straightness under light conditions with visual feedback, as a function of visual weight. Black line represents the mean straightness observed in real flies. (**h**) Simulated straightness under dark (black) and light (gold) conditions for the chosen noise and visual weights. Shaded areas represent the standard error of straightness observed in real flies under light and dark conditions (Cruz et al. 2021)

### Figure6-S1: Transgenic lines used in behavioral experiments

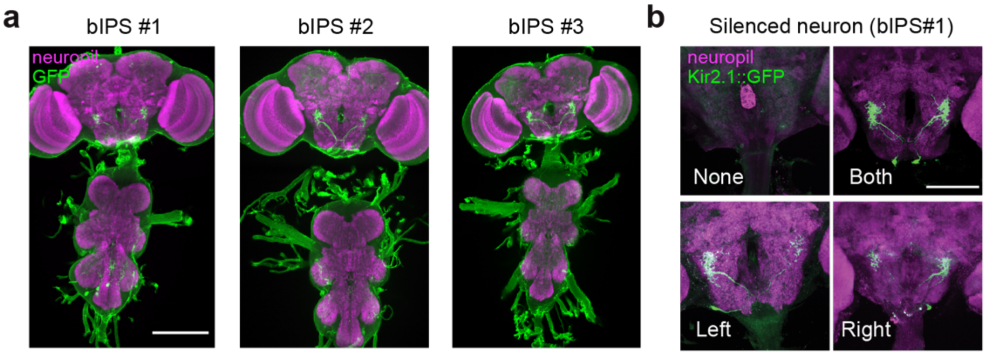


**Figure 6-S1: Transgenic lines used in behavioral experiments** (**a**) Posterior view of maximal projection stacks of three different Split-Gal4 lines labeling bIPS. Neuropil (nc82) staining is shown in magenta and membrane targeted GFP expression is shown in green. Scale bar, 200 µm. (**b**) Posterior view of maximal stack projection of example brains with stochastic silencing of bIPS. Neuropil (nc82) staining is shown in magenta and membrane-targeted, Kir2.1-bound GFP expression is shown in green. Scale bar, 100 µm. For the complete list of genotypes of all transgenic lines, please see **Supplementary Table 3**.
